# A mathematical theory of relational generalization in transitive inference

**DOI:** 10.1101/2023.08.22.554287

**Authors:** Samuel Lippl, Kenneth Kay, Greg Jensen, Vincent P. Ferrera, L.F. Abbott

## Abstract

Humans and animals routinely infer relations between different items or events and generalize these relations to novel combinations of items. This allows them to respond appropriately to radically novel circumstances and is fundamental to advanced cognition. However, how learning systems (including the brain) can implement the necessary inductive biases has been unclear. Here we investigated transitive inference (TI), a classic relational task paradigm in which subjects must learn a relation (*A* > *B* and *B* > *C*) and generalize it to new combinations of items (*A* > *C*). Through mathematical analysis, we found that a broad range of biologically relevant learning models (e.g. gradient flow or ridge regression) perform TI successfully and recapitulate signature behavioral patterns long observed in living subjects. First, we found that models with item-wise additive representations automatically encode transitive relations. Second, for more general representations, a single scalar “conjunctivity factor” determines model behavior on TI and, further, the principle of norm minimization (a standard statistical inductive bias) enables models with fixed, partly conjunctive representations to generalize transitively. Finally, neural networks in the “rich regime,” which enables representation learning and has been found to improve generalization, unexpectedly show poor generalization and anomalous behavior. We find that such networks implement a form of norm minimization (over hidden weights) that yields a local encoding mechanism lacking transitivity. Our findings show how minimal statistical learning principles give rise to a classical relational inductive bias (transitivity), explain empirically observed behaviors, and establish a formal approach to understanding the neural basis of relational abstraction.

## 1 Introduction

Humans and animals have a remarkable ability to generalize to circumstances that are radically different from their prior experience. They are able to do so, in part, by learning relationships between different events or items and extending these relationships to novel combinations of components [1]. Such relational generalization is important across a broad range of domains: for example, subjects can reason about social relationships between individuals they have never seen interact [2–4], take a novel route between familiar locations [5], or apply familiar tools to novel problems [6]. Accordingly, relational cognition has been implicated in a variety of cognitive abilities, including social cognition [7], spatial navigation [8], and logical and causal cognition [9].

It is unclear how living subjects, and learning systems more generally, can learn the kinds of abstractions needed for relational generalization. To generalize from limited experience, subjects (whether they are humans, animals, or learning models) need an “inductive bias”: a disposition towards certain behaviors among the many that are consistent with past experience [10]. Much attention has been devoted to clarifying suitable inductive biases on standard statistical tasks, which require generalization to nearby data points (“near transfer” or “in-distribution generalization”) [11–16]. An important instance of such a “statistical inductive bias” is norm minimization, which selects the model parameters with the smallest weight norm and is at the core of many successful learning models [17–19]. Both theoretical and practical insights suggest that such a solution is likely to generalize well in distribution [20, 21]. In contrast, our understanding of how learning models perform relational tasks, which require generalization to radically different circumstances (“far transfer,” [22] “out-of-distribution generalization,” [23] or, more specifically, “generalization to the unseen” [24]), has been much more limited [25]. Addressing this gap is essential to understanding how living subjects perform these tasks.

To investigate this question, we studied transitive inference [TI; Fig. 1a; 26–28], a classical cognitive task that tests if humans or animals can generalize transitively. In this task, subjects are presented with pairs of items (Fig. 1b) and must pick the “larger” item according to an implicit hierarchy (*A* > *B* > … > *G*, Fig. 1a). Importantly, subjects are not informed of this task structure [cf. 29, 30] and, further, only receive feedback about the correct response on adjacent pairs (*AB, BC*, …, *FG*). Hence, they must infer the underlying relation and use transitive inference to determine the correct response to non-adjacent pairs (*B* > *E*, Fig. 1c). Notably, humans [32] and a variety of animals (ranging from monkeys [33, 34] and rodents [35] to wasps [36] and fish [37]) perform TI successfully. Moreover, they show consistent behavioral patterns (in their reaction time and accuracy, Fig. 1d), which may be informative of the underlying neural implementation. First, subjects’ performance improves with increasing separation in the hierarchy [**“symbolic distance effect**,**”** e.g. 32, 38–42]. Second, subjects’ performance tends to be better for trials involving more terminal (e.g. A and G) rather than more intermediate (e.g. D and E) items [**“terminal item effect**,**”** e.g. 31, 43–45]. Third, some studies indicate better performance on training trials (which have a symbolic distance of 1) than on test trials with a symbolic distance of 2 [29, 42, 46–48, Fig. 1e], a limited violation of the symbolic distance effect which we here refer to as the **memorization effect**. Finally, subjects’ performance is often better for item pairs towards the beginning of the hierarchy than item pairs towards the end (the **“asymmetry effect**,**”** which we do not address but return to in the discussion) [e.g. 29].

**Figure 1:**
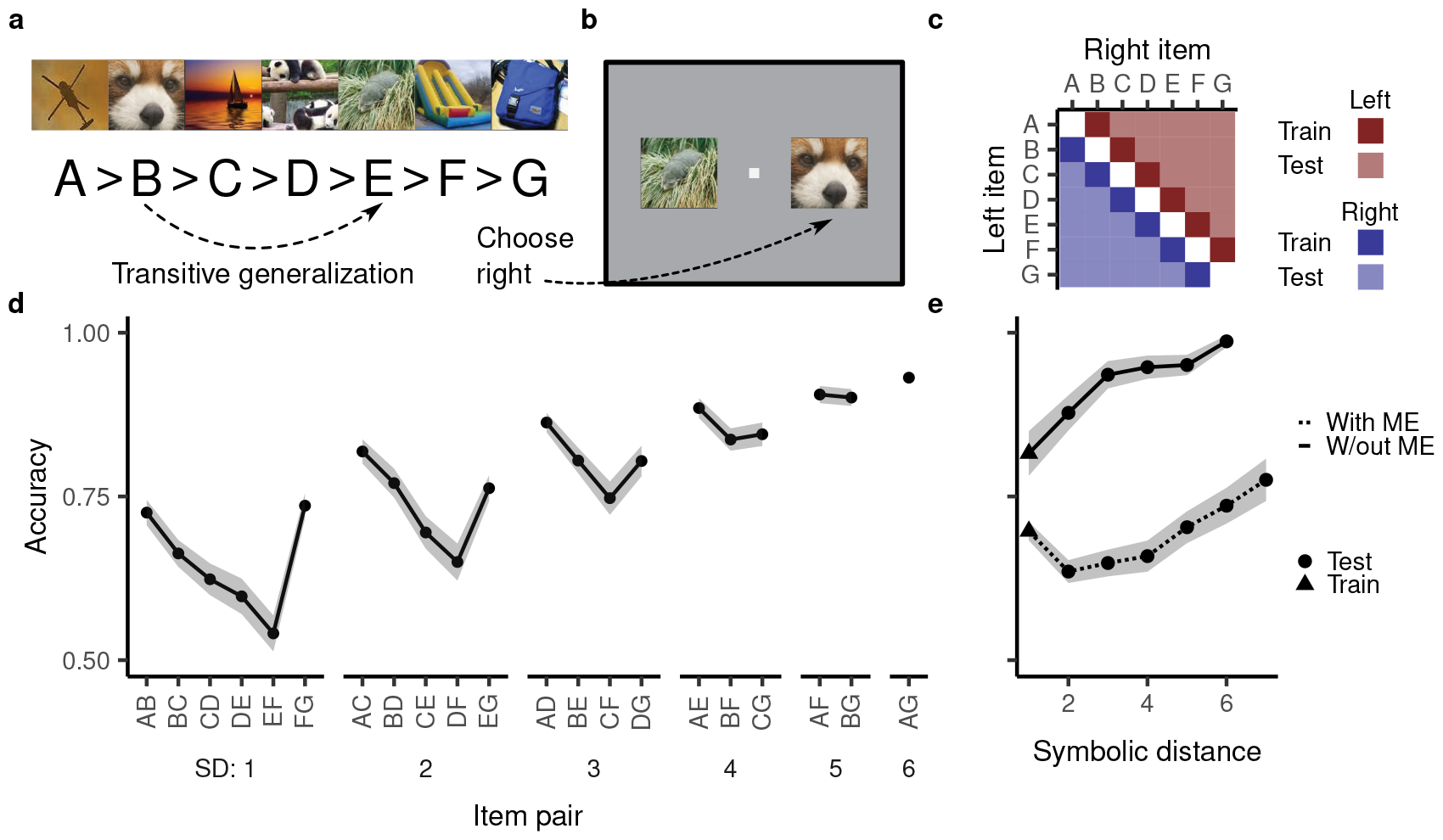
Transitive inference and behavioral patterns observed in subjects performing this task. **a**, Example stimuli taken from [31]. An example generalization (*B* > *E*) is highlighted. **b**, In a given trial, the subject is asked to choose between the two presented items. They are rewarded if the chosen item is “larger” according to the underlying hierarchy. This panel depicts a test trial where successful performance would consist in picking the item on the right. **c**, Schematic of the training and test cases. **d**, Example accuracy on all training and test cases (in this case by rhesus macaques). Terminal item, symbolic distance, and asymmetry effect are apparent in the plot. In the subsequent figures, we leave off the item pair labels but use the same ordering. Symbolic distance (SD) is the separation in the rank hierarchy. The data is reproduced from [31]. **e**, Symbolic distance effect with and without a memorization effect (ME). Data without ME reproduced from [31] and data with ME reproduced from [29]. Shaded regions in both panels indicate mean ± one standard error.

Many simple learning models have been shown to perform transitive inference. However, the inductive biases that give rise to this ability are not well understood. Various learning models associate a numerical “rank” with each presented item and choose whichever item has a higher rank [29, 31, 49–51]. Such models are pre-configured to generalize transitively and leave unclear how transitive generalization could arise from more basic learning principles. They also leave unclear how the brain, which is not constrained in this way [52], could implement TI. Intriguingly, several studies have found that generic neural networks can generalize transitively, suggesting that statistical learning principles can sometimes give rise to a suitable relational inductive bias [30, 53–55]. However, TI in neural networks has largely been studied through simulations, rather than analytically [cf. 56], raising the question of when and how statistical learning models can implement TI.

Here we show, via both analytical approaches and simulations, that a broad range of biologically relevant learning models generalize transitively and recapitulate the symbolic distance, terminal item, and memorization effect. We first consider models with “additive” representations that represent the two presented items independently, and show that they are constrained to implementing a transitive relation. If models additionally represent nonlinear conjunctions between items (as is important for many other tasks) but use norm minimization to determine their readout weights, they also generalize transitively. Remarkably, the same learning principle that underpins many instances of successful near transfer also enables this instance of successful far transfer. We further show that, for TI, the effect of a particular choice of internal representation can be characterized by a single scalar “conjunctivity factor.” Finally, we consider models which adapt their internal representation to a given task, an ability thought critical to human and animal cognition. Surprisingly, we find that this impairs performance on TI and leads to behavioral patterns that deviate from those in living subjects. Notably, this anomalous behavior is explained by the rich regime’s implementation of a different form of norm minimization, namely one over all weights in the network rather than just the readout weights.

At first glance, TI appears to be a complex task involving relational cognition and application of the transitive rule. Nevertheless, we can characterize a broad range of learning models performing this task in exact analytical terms, explaining how they give rise to the rich behavioral patterns observed in living subjects. In doing so, our investigation clarifies systematically how a learning principle that has largely been considered in the context of near transfer, also implements an important instance of far transfer and relational abstraction.

## 2 Model setup

We represent individual items as high-dimensional vectors. A trial input is a concatenation of the two vectors *X, Y* corresponding to the two presented items. We generally consider a learning model *f* which represents this input as a numerical vector *g*(*X, Y*) (in the simplest case, this could be the input vector itself). The model then computes a linear readout from that representation: *f* (*X, Y*) = *w* ∘ *g*(X, Y). A positive model output (*f* (*X, Y*) > 0) corresponds to *X* > *Y*, whereas a negative output (*f* (*X, Y*) < 0) corresponds to *X* < *Y*. We generally assume that *w* is learned from the training trials, whereas *g*(*X, Y*) remains fixed (for example arising from the representation in a neural network with random or pre-learned weights). In the final two sections, we investigate models that also learn *g*(*X, Y*).

Inputs where *f* (*X, Y*) = 0 lie on the model’s decision boundary. Accordingly, a higher magnitude of the margin (*f* (*X, Y*) if *X* > *Y* or − *f* (*X, Y*) if *X* < *Y*) corresponds to a larger distance from the decision boundary. We take this to indicate better performance, corresponding to higher choice accuracy and lower reaction time. This is a standard assumption in models of decision making [30, 57–59]. For example, in a drift diffusion model [60], the output *f* (*X, Y*) determines the model’s drift rate. The higher the drift rate, the less susceptible the model is to noise (improving accuracy) and the faster it will cross its threshold (improving reaction time) [61].

## 3 Results

### 3.1 An additive representation yields transitive generalization

To perform TI, or indeed any kind of relational inference, a learning model’s representation of items *X* and *Y*, *g*(*X, Y*), should reflect the fact that *X* and *Y* are separate items. In the most extreme case, this would amount to an *additive* representation, where *g*(*X, Y*) is a sum of two separate representations *g*_1_(*X*) and *g*_2_(*Y*): *g*(*X, Y*) = *g*_1_(*X*) + *g*_2_(*Y*). A simple instance of this is a model architecture where nodes respond exclusively either to *X* or *Y* (Fig. 2a).

**Figure 2:**
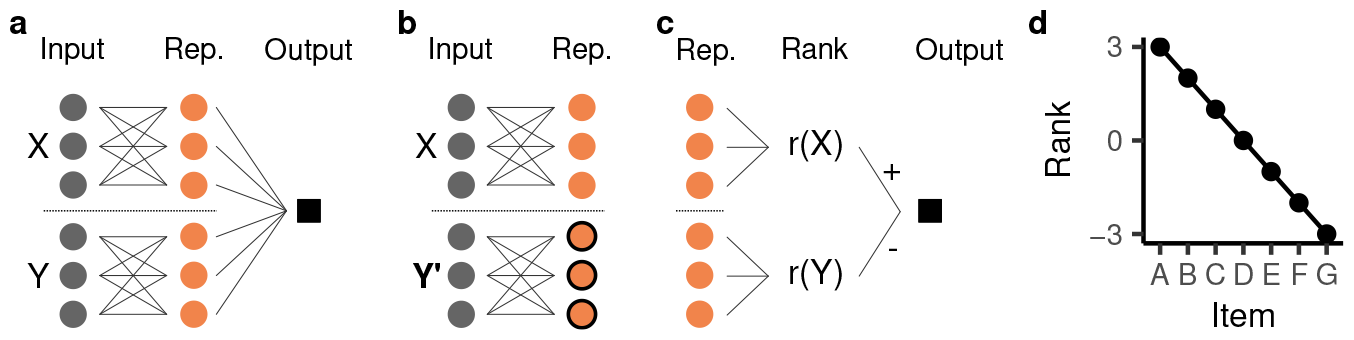
An additive representation yields transitive generalization and the symbolic distance effect. **a**, Schematic illustration of an additive representation (rep.). The first three orange nodes represent the first item (*X*) and the latter three represent the second item (*Y*). **b**, Replacing the second item only results in changes to the highlighted half of the units. **c**, The readout weights of the model can be grouped into those pertaining to item *X* and those pertaining to item *Y*. The model’s output can be understood as assigning a “rank” *r*(*X*) and *r*(*Y*) to each item and then computing *r*(*X*) − *r*(*Y*). **d**, Example of a model’s rank assignment.

A change in one of the two items will leave half of the additive representation unchanged (Fig. 2b), implementing a kind of compositionality [62]. A linear readout from an additive representation, *f* (*X, Y*) = *w* ∘ *g*(*X, Y*), is a sum of responses to each individual item, *w* ∘ *g*_1_(*X*) + *w* ∘ *g*_2_(*Y*), and therefore also additive. The consequences of this are especially clear in the case of a model without a choice bias, i.e. if *f* (*X, X*) = *w* ∘ *g*_1_(*X*) + *w* ∘ *g*_2_(*X*) = 0 (Appendix S1.1 considers the biased case). In this case, we know that −*w* ∘ *g*_1_(*X*) = *w* ∘ *g*_2_(*X*), and, as a result, the model’s decision function can be expressed as

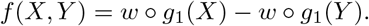

This means that the model necessarily learns to assign a scalar “rank” *r*(*X*) = *w* ∘ *g*_1_(*X*) = − *w* ∘ *g*_2_(*X*) to each item, and computes its decision by comparing the two ranks (Fig. 2c):

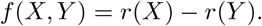

Note that *f* (*X, Y*) > 0, i.e. the decision to choose the left item, is equivalent to *r*(*X*) > *r*(*Y*).

As shown in previous work [e.g. 29, 31, 43], learning systems that are pre-configured to have such a ranking system yield both transitive generalization and the symbolic distance effect. This is because, to learn the training set, the model’s ranks must be monotonically decreasing:

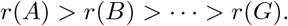

As an example, if the model output has a margin of one for all inputs composed of adjacent items, the rank must decrease by one for each successive item (Fig. 2d). A monotonically decreasing rank directly implies that non-adjacent items are also ordered correctly and, consequently, that the model will generalize transitively. Further, item pairs with larger symbolic distance will have a larger difference in their ranks, giving rise to a symbolic distance effect.

Critically, and in distinction to previous work, the above model structure (additive representation) does not explicitly pre-configure or assume a ranking system. Rather, the above analysis shows that any learning model that implements an additive input representation necessarily implements a ranking system and thus encodes a transitive relation. Further, the above analysis makes no assumptions about how the model learns. As long as the model correctly classifies the training cases, it will correctly classify all test cases, i.e. transitively generalize, and it will exhibit a symbolic distance effect.^1^

### 3.2 A single scalar fully characterizes a broad range of relational representations

The above analysis indicates that an additive representation would enable the brain to perform TI. However, neural representations are not thought to be fully additive [63]. Indeed, non-additive representations (Fig. 3a) are important for learning relevant tasks across a broad range of domains and, with some differences in implementation, are known under a correspondingly broad range of names, including conjunctive [64] or configural [65] representations, nonlinear features, and representations with mixed selectivity [63]. We next asked whether and how non-additive representations can support transitive generalization.

**Figure 3:**
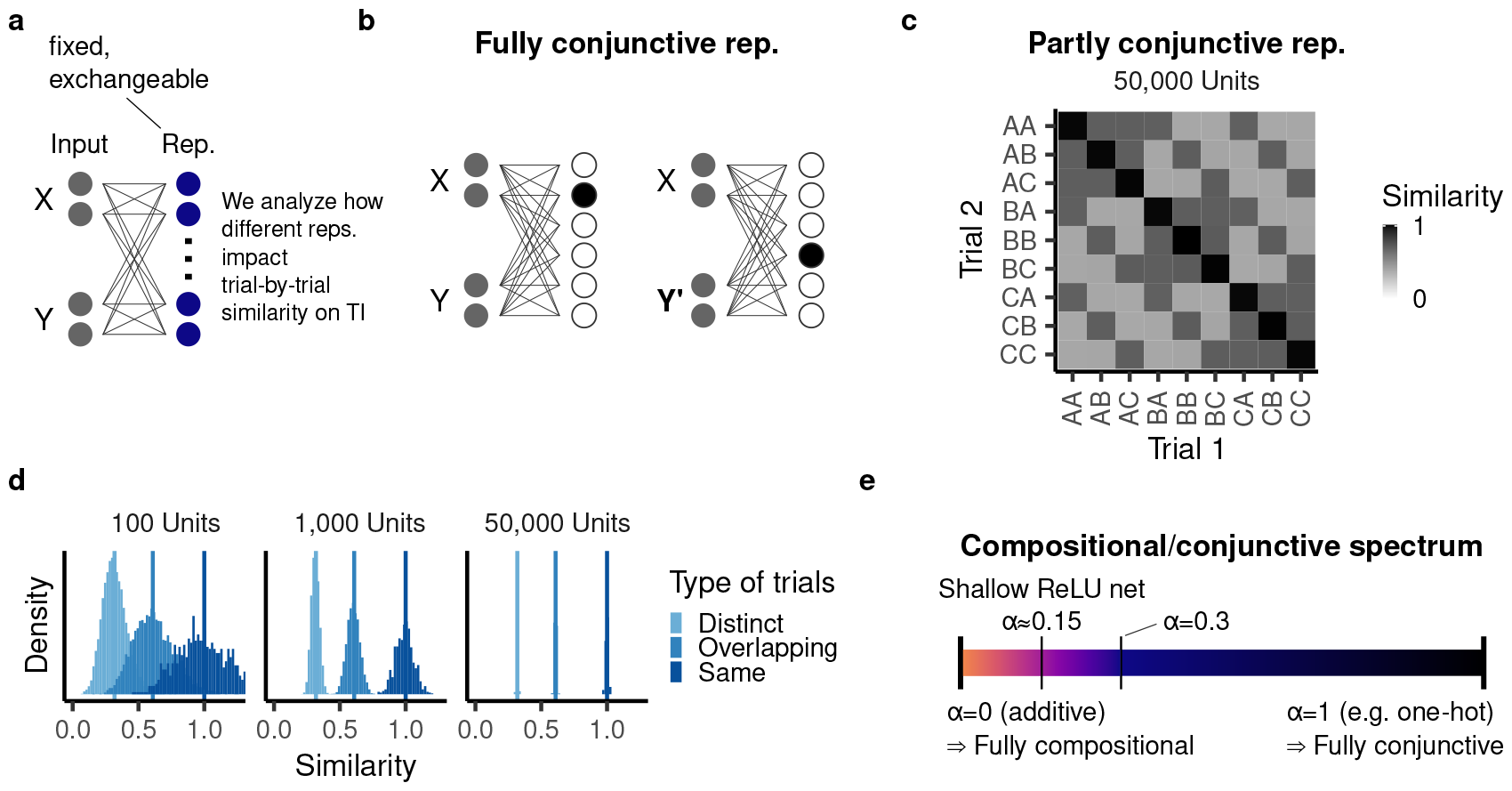
Non-additive representations of TI cases. **a**, A non-additive representation encodes nonlinear interactions between items. **b**, A one-hot representation represents each combination of items by a distinct node. **c**, Representational similarity (cross-correlation) between a subset of trials in a ReLU network with 50,000 units. **d**, Representational similarity between all possible trials in networks with different numbers of hidden units, organized according to the type of trials. **e**, The conjunctivity factor characterizes a given representation according to how similarly it represents overlapping trials. Additive representations lie at one end of the spectrum, whereas one-hot representations lie at the other end.

To begin, consider the most extreme conjunctive case: a one-hot representation in which each composition of items is represented by a different hidden unit (Fig. 3b). In this case, a change in one of the two items yields a completely different representation. Such a model is able to memorize the training cases, but cannot generalize transitively.

Many representations, whether in the brain or in other learning systems, are neither fully conjunctive nor fully additive, but rather lie in between these two extremes [63, 66–69]. To characterize this spectrum formally, we considered the representational similarity between two trials (*X, Y*) and (*X*^′^, *Y*′), as measured by their dot product, ⟨*g*(*X, Y*), *g*(*X*^′^, *Y*′)⟩ (leaving the representations *g*(*X, Y*) fixed). We assumed that the representational similarity between two trials only depends on whether these trials are distinct ((*X, Y*) and (*X*^′^, *Y*′)), overlapping ((*X, Y*) and (*X*^′^, *Y*) or (*X, Y*′)), or identical ((*X, Y*) and (*X, Y*)) (Fig. 3c). We call this the “exchangeability assumption.”

The exchangeability assumption captures the fact that model behavior should not depend on the particular (i.e. arbitrary) hierarchy in which the set of items is arranged [70]. To promote exchangeability, we assumed that all input items are equally correlated with each other. In this case, most commonly used nonlinear representations of that input satisfy exchangeability as well. As a paradigmatic learning model, we considered a neural network with a ReLU nonlinearity and random weights. By determining the expected value of the representational similarity analytically (Appendix S1.3), we found that the network’s hidden layer, in expectation, satisfies the exchangeability assumption (Fig. 3c). This is because even though the hidden layer computes a nonlinear transformation, it partially inherits the input’s similarity structure [71].

In network simulations, the empirical representational similarity exhibits some variance around the expected value due to the random weight initialization. However, this variance vanishes as the network’s hidden layer becomes wider (Fig. 3d).

Under the exchangeability assumption, we found that for learning models that only modify their readout weights, a single scalar, which we call the **“conjunctivity factor”** *α*, fully determines their TI task behavior, i.e. all models that have representations with the same *α* exhibit the same behavior on TI (see Appendix S1.2). The conjunctivity factor is given by:

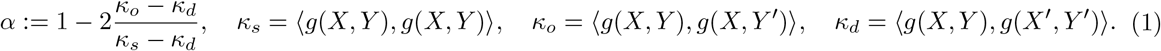

Here *κ*_*s*_, *κ*_*o*_, and *κ*_*d*_ are the similarity between identical, overlapping, and distinct pairs, respectively. Notably, we can extend the definition of the conjunctivity factor to non-exchangeable representations by taking the average over all identical, overlapping, or distinct pairs.

For an additive representation, half the units share their activation between different overlapping pairs. As a result, the similarity between overlapping pairs is halfway between that of distinct and that of identical pairs, corresponding to *α* = 0. At the other extreme, *α* = 1 indicates that overlapping pairs are encoded in the model with equal similarity to each other as completely distinct pairs (as is the case for a onehot representation). Consequently, *α* systematically characterizes the spectrum from fully additive to fully conjunctive representations (Fig. 3e).

In most commonly used representations, overlapping trials are more similar to each other than distinct trials and therefore have an intermediate value for *α* (“partly conjunctive” representations). This is because the input space represents overlapping trials as more similar than distinct trials and, as noted above, the hidden layer partly inherits the input’s similarity structure. Through analytical computation, we found that a random ReLU network with one hidden layer, for example, has *α* ≈ 0.15 (Eq. (S28); Fig. 3c,d).

Importantly, the network giving rise to the representation could have an arbitrary number of layers; all we need to know is the conjunctivity factor of the network’s final layer. In fact, in Appendix S1.3, we compute the conjunctivity factor of neural networks with arbitrary depth, finding that *α* increases with depth. Indeed, as the networks become deeper, *α* eventually approaches one. However, if the connectivity structure is modified so a part of each layer is additive (but is, in turn, connected to the next layer’s conjunctive representation as well), *α* can be reduced and, in fact, will eventually approach zero. This illustrates the crucial impact that network connectivity can have on representational structure.

For transitive inference, the conjunctivity factor raises two questions. First, how does *α* > 0 affect TI behavior? Second, if the learning model’s internal representation is modifiable rather than fixed, how does this affect the conjunctivity factor and subsequent TI behavior?

### 3.3 Norm minimization and partly conjunctive representations yield transitive generalization

Unlike an additive representation, a representation with *α* > 0 is not constrained to implementing a transitive relation. To understand how models with partly conjunctive representations perform on TI, we need to consider additional constraints. In particular, we analyzed models in which the learning of readout weights implements norm minimization (Fig. 4a), a paradigmatic statistical inductive bias (Fig. 4b). Under the exchangeability assumption, we were able to characterize model behavior on TI through exact analytical solutions. As noted above, to perform this analysis, we do not need to know the particular representation implemented, only its conjunctivity factor *α*.

**Figure 4:**
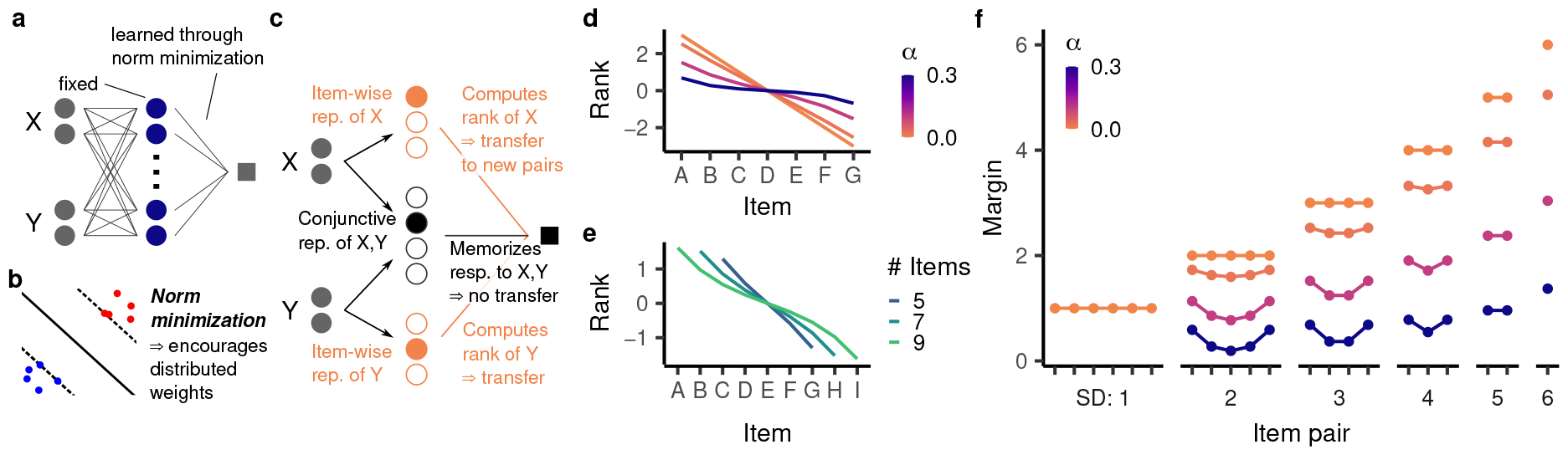
We analyze the behavior of models having readout weights trained with norm minimization. **a**, Schematic illustration of the setup. **b**, Norm minimization implements a useful inductive bias for generalization to nearby data points. On categorization tasks, it determines the hyperplane separating the two categories with the maximal margin. **c**, Intuitively, a partly conjunctive representation is given by an itemwise representation of *X* and *Y* concatenated with a fully conjunctive representation of *X* and *Y*. The readout from the item-wise representations computes a rank for each item that transfers to overlapping pairs. The readout from the fully conjunctive representation memorizes a response to a given pair and does not transfer to overlapping pairs. Because norm minimization encourages distributed weights, it finds a solution that partly uses the item-wise representation and hence computes a rank. This leads to transitive generalization. **d**,**e**, The emergent rank representation at the end of training (Eq. 2) for (d) seven items and different values for *α*; and (e) *α* = 0.1 and different numbers of items. **f**, For seven items and different values for *α*, the corresponding margin for all trials. Item pairs are arranged by their position in the hierarchy, as in Fig. 1e.

On the training cases, the minimal norm model necessarily assigns a margin of ± 1, as dictated by the desired output. On the test cases, however, our analysis revealed an intriguing emergent behavior: the model’s response to item pair (*i, j*) invariably reflects a ranking system:

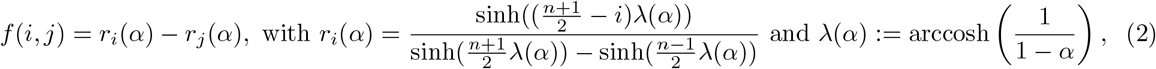

where *n* is the total number of items (see Appendix S1.4.3 for the derivation of this result). This is remarkable as the model architecture is not constrained or pre-configured to implement a ranking system. Rather, the behavior is a consequence of the principle of norm minimization, operating on a partly conjunctive representation. Importantly, as implied by its ranking system, the model generalizes transitively (as long as overlapping trials are represented as more similar than distinct ones, i.e. *α* < 1), and exhibits a symbolic distance effect.

For an intuition as to why a ranking system emerges, consider a particular class of representations having one-hot representations of the first and second item individually as well as their conjunction (Fig. 4c). This representation will have a conjunctivity factor of *α* if the item-wise units are weighted by 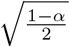 and the conjunctive units are weighted by 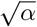. Changes in the item-wise unit weights correspond to changes in the model’s rank, as they generalize to overlapping trials. In contrast, changes in the conjunctive unit weights correspond to memorization, as they have no effect on overlapping trials (akin to a fully conjunctive representation). In principle, the model could learn the training set through pure memorization, i.e. changes in the conjunctive unit weights alone. However, because more distributed weights tend to have a smaller weight norm, norm minimization causes the model to learn the training premises by changing both the conjunctive unit weights (resulting in memorization) and the item-wise unit weights (resulting in transitive generalization). Thus, the partial conjunctivity is necessary for the existence of an item-wise population; the principle of norm minimization ascertains that this population is implicated in the learning process.

Beyond transitive generalization, the principle of norm minimization gives rise to several empirically observed behavioral patterns. The hyperbolic sine making up the rank expression compresses more intermediate items more strongly than more terminal items (Fig. 4d), thus giving rise to a terminal item effect (Fig. 4f). This effect becomes stronger for higher values of *α*. A higher *α* also compresses the ranking more strongly overall, leading to lower margins on the test set. As *α* approaches one (the fully conjunctive case), the ranking becomes entirely flat and therefore no longer supports transitive generalization.

The form of the ranking also depends on the total number of items. Specifically, intermediate items are compressed more strongly when there are more items in total, an effect that is moreover dependent on *α*. For *α* = 0, the ranking grows linearly with the number of items, whereas at higher values of *α*, the model ranking’s overall range (between the first and last item) is almost invariant to the number of items (Fig. 4e). Finally, when *α* > 0, the model assigns a larger margin to the training cases than specified by the ranking. Intuitively, this is because the conjunctive unit weights contribute to model behavior on the training cases, but do not transfer to the test cases. Since a higher *α* compresses the ranks further, it leads to a higher discrepancy between training and test behavior. At a sufficiently high *α*, the margin of the training cases is larger than that of the test cases with a small symbolic distance (Fig. 4f), giving rise to a memorization effect. While Ciranka *et al*. [29] explained this effect by fitting an explicit memorization parameter, our analysis reveals that it is an emergent consequence of having a nonlinear representation with sufficiently high conjunctivity factor. Notably, the fact that the memorization effect, according to our model, only arises in a subset of relational representations may explain why it is only occasionally observed in living subjects. In contrast, the symbolic distance and terminal item effect arise across the full spectrum of relational representations (except the fully conjunctive case) and indeed are also observed much more consistently in living subjects.

The above analytical solutions depend on the exchangeability assumption. In practice, the trials might be represented in a manner that violates this assumption. Indeed, we already saw that though randomly sampled features satisfy exchangeability in expectation, a model with insufficiently many of those features will have a non-exchangeable representation due to finite samples (Fig. 3d). In simulations (see Appendix S2.2), we found that a slight violation of exchangeability (e.g. resulting from a representation with many random features) does not change model behavior substantially. For larger violations (e.g. resulting from a representation with fewer random features), model behavior deviates from our theoretical account, but is still well approximated by it. Further, our account captured the average behavior across many models with a small number of random features almost exactly. This suggests that models with non-exchangeable representations behave differently from those with exchangeable representations but our analytical solutions can still be useful for understanding their behavior.

Our analysis demonstrates that norm minimization can explain not only transitive generalization, but also the symbolic distance effect, the terminal item effect (though only on test cases), and the memorization effect. We next characterize two popular statistical learning models implementing norm minimization: ridge regression and gradient flow, as applied to learning of network readout weights.

### 3.4 Learning through gradient flow or with weight regularization smoothly approaches the minimal norm solution

To see how the principle of norm minimization governs TI behavior across learning, we analyzed models with readout weights learned through either ridge regression [Fig. 5a; 17] or gradient flow, using mean squared error as the loss function *L*(*w*) (with the target response on the training cases being ±1). Ridge regression minimizes the sum of this loss and the squared *L*_2_-norm of the model weights, i.e.

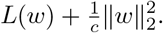

Here, the regularization coefficient *c* balances the weight penalty 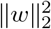 with the minimization of the loss function *L*(*w*). Biologically, such weight regularization could, for instance, be implemented by mechanisms of synaptic homeostasis [72]. Gradient flow, on the other hand, assumes that a model minimizes *L*(*w*) by following the pointwise gradient exactly. This approximates gradient descent with a small learning rate, and is more amenable to formal analysis [73]. In this case, the learning duration *t* determines how well the model has learned to minimize the loss function in the allotted time. At initialization, the model should be agnostic to all choices (i.e. output zero) and we therefore assumed that the weights are initialized at zero.

**Figure 5:**
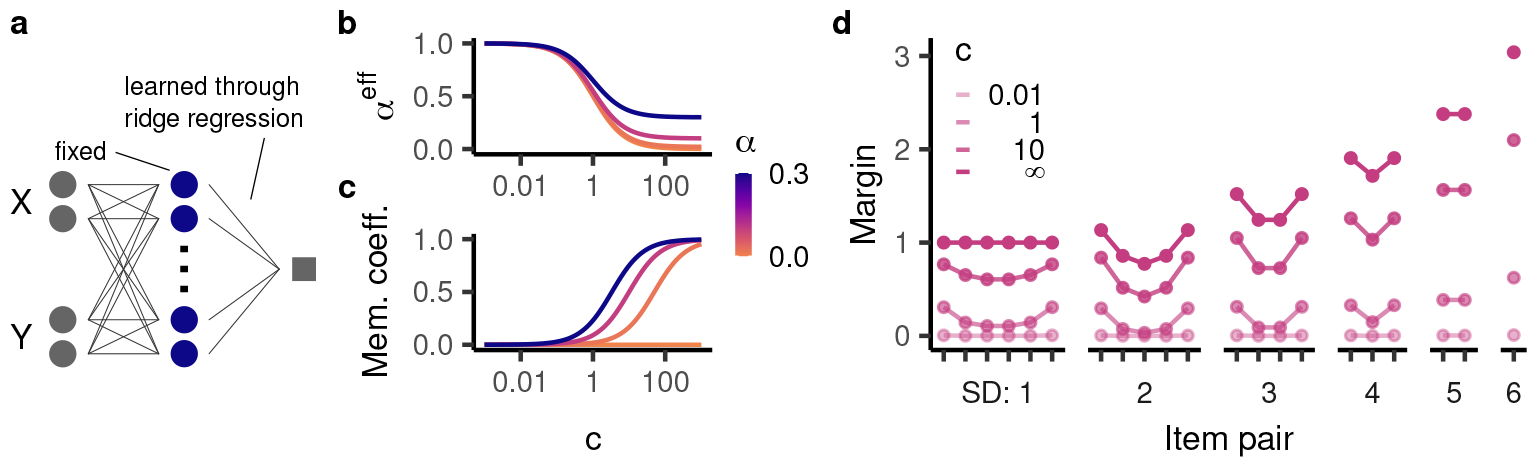
TI behavior for models with a fixed representation and readout weights trained using regularized regression. **a**, Illustration of the setup. **b**,**c**, The (b) effective conjunctivity factor and (c) memorization coefficient as a function of the inverse regularization coefficient *c*. **d**, Generalization behavior for *α* = 0.1 and different values of *c*. The margins overall become larger as *c* increases.

The regularization coefficient *c* and the learning duration *t* play a similar role in the two learning models. With *c* = 0, the weight penalization is infinitely more important than the task-based component of the loss function and therefore the model weights are all zero. Similarly, at *t* = 0, the weights are initialized to zero. On the other hand, in the limit of infinite training (*t* → ∞), the model converges to the minimal norm solution determined in the section above [74]. This is also the case for the limit of models trained with increasingly small weight penalization (*c* → ∞) [75]. Ridge regression and gradient flow are therefore two instances of common learning models which can perform TI by implementing the principle of norm minimization.

Going beyond these limits, we obtained exact solutions to model behavior for arbitrary *t* or *c*. For ridge regression (see Appendix S1.4.3), we found that the test behavior of a model with weight regularization and a given conjunctivity factor *α* is equivalent to that of a model without weight regularization but a different conjunctivity factor. This “effective conjunctivity factor” *α*^eff^ depends on both *α* and *c* (see Lemma S1.4):

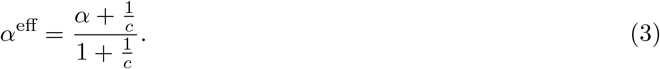

*α*^eff^ is generally higher (i.e. more conjunctive) than *α* and, as *c* becomes larger, gradually approaches *α* from above (Fig. 5b). Thus, a model with smaller *c* has a more compressed rank and a more pronounced terminal item effect.

On the training cases, the model’s performance is boosted compared to the rank difference, just like in the case of norm minimization. Specifically, its behavior is given by

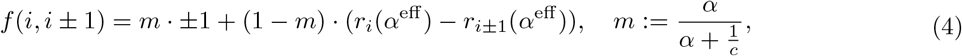

i.e. a mixture of memorization (assigning a constant margin of ± 1) and reliance on the ranking system. The balance between the two behaviors is specified by the “memorization coefficient” *m* ∈ [0, 1]. *m* starts out at zero, indicating no memorization and full reliance on the ranking system. As *c* grows larger, *m* increases as well (Fig. 5c). The smaller *m*, the more strongly model behavior relies on the ranking. In particular, this partial reliance leads to a terminal item effect on the training cases in addition to the test cases (Fig. 5d). For *c* → ∞, *m* converges to 1, indicating full memorization as observed for the minimal norm solution.

For gradient flow, we found qualitatively similar solutions for model behavior (Appendix S1.4.4). In particular, the model’s behavior on test cases can be described by a ranking system throughout learning. Further, the model ranks gradually approach the minimal norm solution and, at earlier stages of training, have a more pronounced terminal item effect. Finally, the learning dynamics give rise to a transient terminal item effect on the training cases that vanishes as *t* → ∞.

Humans and animals generally exhibit a terminal item effect on the training set. Our analysis suggests that this could be caused by either weight regularization or incomplete training (or both), arising from a mechanism that is related to but distinct from the mechanism giving rise to the terminal item effect on the test cases.

### 3.5 The conjunctivity factor exposes a tradeoff between learning transitive and non-transitive relational tasks

Our analysis thus far indicates that a higher conjunctivity factor generally yields worse performance on transitive inference. Indeed, if maximal generalization performance on TI were the sole aim, models with fully additive representations would be ideal. However, as noted above, humans and animals learn a broad range of relational tasks, not all of them transitive. Because models with fully additive representations are constrained to implementing a transitive relation, this makes partially conjunctive representations necessary (Fig. 6a).

**Figure 6:**
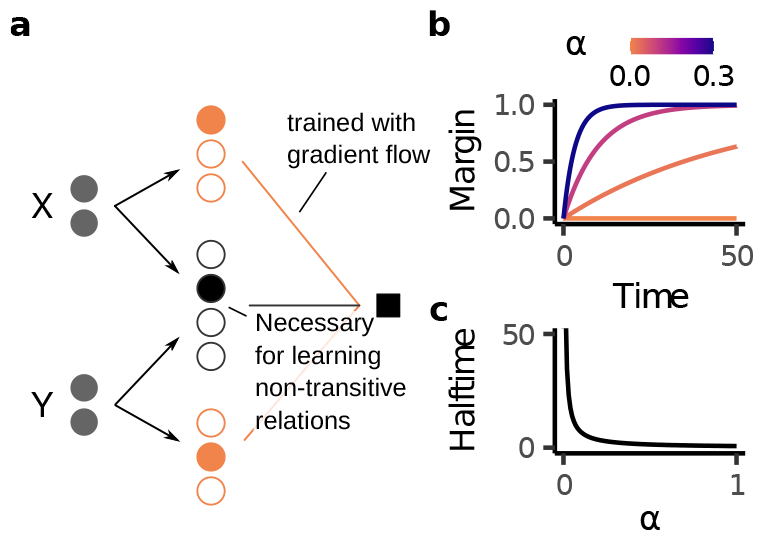
Behavior of models trained with readout weights trained through gradient flow on transverse patterning. **a**, Learning a non-transitive relation requires a conjunctive component. **b**, Margin over time for different values of *α*. **c**, Halftime, i.e. time until the model reaches a margin of 0.5, as a function of the conjunctivity factor *α*.

To investigate how different representational geometries change model behavior beyond transitive inference, we considered transverse patterning (“rock, paper, scissors” with more than three items, e.g.: *A* > *B, B* > *C*, …, *F* > *G, G* > *A*). Transverse patterning exemplifies a non-transitive relation and has been shown to be learned by both humans and various animals [34, 52, 76–79].

As with TI, we considered models with fixed, exchangeable representations with readout weights trained through gradient flow (Fig. 6a). We found that as long as *α* > 0, models were able to learn the task by relying on the conjunctive population. Further, by solving the learning dynamics of gradient descent analytically (see Appendix S1.6), we found that a higher *α* leads to faster learning of the training trials (Fig. 6b,c). In contrast, on TI, a higher *α* leads such models to have a *smaller* test margin. These behaviors highlights a potential tradeoff that *α* imposes across different kinds of relations. In particular, our analysis predicts that subjects who are better at transitive inference should be slower to learn transverse patterning and vice versa.

One strategy for avoiding the tradeoff described above is representation learning. If models were able to adapt their internal representations to a given task, they could in principle learn an additive representation for TI and a non-additive representation for non-transitive relational tasks such as transverse patterning. We now investigate this hypothesis.

### 3.6 Neural networks with adaptive representations deviate from living subjects’ TI behavior

Deep neural networks, which learn by updating their internal weights, have become increasingly relevant both as artificial learning systems [80] and models of cognitive processes [81, 82]. Importantly, recent work has revealed that their generalization behavior fundamentally depends on the magnitude of their initial weights [83, 84]. For large initial weights, learning dynamics can be approximated by gradient flow on a particular fixed-feature model called the neural tangent kernel [NTK; 85, 86]. Thus, even though the network updates its internal weights, it effectively still relies on a fixed representation. Accordingly, this regime is often called “lazy” [83]. In contrast, neural networks initialized from sufficiently small values learn truly task-specific representations (“rich regime”). Broadly considered, adapting a model’s internal representation to a given task could address the competing demands imposed by the wide range of tasks subjects need to perform.

Indeed, the rich regime is seen as essential to the remarkable generalization capabilities of deep neural networks [83, 87, 88]. There is also widespread evidence that subjects adapt their representation to a given task [89, 90] and may do so similarly to deep neural networks trained in the rich regime [91]. In relational tasks, in particular, neural representations change to reflect how different items are related to each other [30, 92–96] and this may be essential for successful generalization on certain tasks [30, 97]. In light of this lazy vs. rich distinction and its potential relevance to biological learning, we investigated through simulations whether deep neural networks are suitable as a model of relational representation learning (Fig. 7a). Specifically, we trained neural networks from different scales of initialization using gradient descent (for details on training, see Appendix S2.3). In light of the important role played by the initialization scale, we covered a broad range of potential values, focusing on three representative values: 1 (resulting in lazy behavior), 10^−3^ (resulting in rich behavior), 10^−16^ (to characterize model behavior in the limit of small initialization).

**Figure 7:**
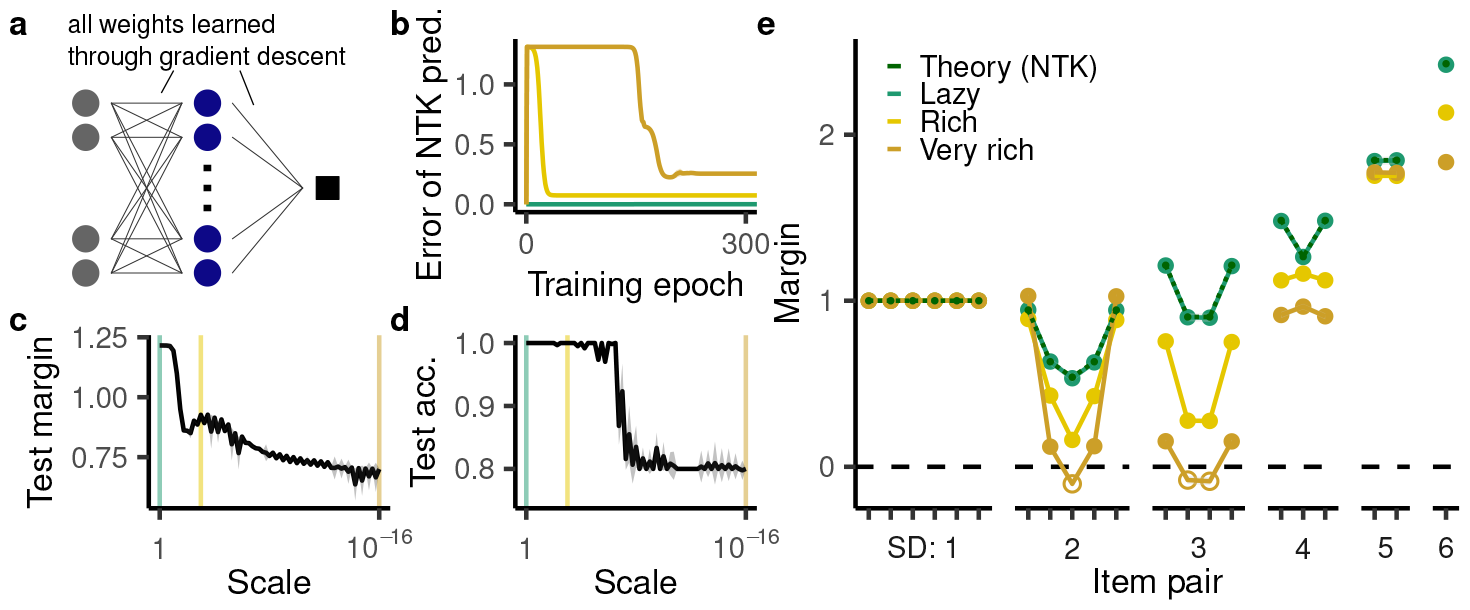
TI behavior in deep neural networks with modifiable representations. We considered twenty instances of a network with a ReLU nonlinearity and one hidden layer with 50,000 units, trained using mean squared error. Appendix S2.5 details other architectures, nonlinearities, and objective functions. Shaded regions indicate mean ± one standard deviation. (Note that some of the behaviors have sufficiently small variance that this shaded region may be invisible.) **a**, Illustration of network training. In contrast to the previous setups, the hidden layer weights were also trained using backpropagation. **b**, The mean squared error of the prediction made by the neural tangent kernel (NTK) at three different initialization scales (lazy: 1; rich: 10^−3^; very rich: 10^−16^). **c**,**d**, The (c) average test margin and (d) test accuracy as a function of initialization scale. The colored lines highlight the three representative values analyzed in more detail in panels b and e. **e**, TI performance according to our NTK-based prediction as well as of networks trained with backpropagation at the three representative scales.

In the lazy regime, we computed the NTK’s conjunctivity factor analytically (Appendix S1.5). For large initialization, the learning trajectory of neural networks trained with full weight updating is approximated by gradient flow on the fixed NTK representation. Hence we were able to use the gradient flow solutions determined in the previous sections to predict the network behavior over the course of training, finding a virtually perfect match with simulations in the case of wide neural networks (Fig. 7b,e, green line). Our account may therefore be able to explain why previous studies [53, 54] found empirically that feedforward neural networks generalize transitively.

Surprisingly, we found that networks trained in the rich regime performed worse at TI. They had a smaller test margin (Fig. 7c) and also made systematic errors on the cases CE, BE, and CF for a sufficiently small initialization scale (Fig. 7d,e, hollow points). Further, and in contrast to all learning models considered thus far, the behavior of neural networks in the rich regime was not consistent with a ranking system. This is apparent from the fact that the networks’ margin at a symbolic distance of three was smaller than at a symbolic distance of two (Fig. 7e). Importantly, in contrast to the previous limited violation of the symbolic distance effect, this cannot be explained by memorization as none of these cases are in the training set. Finally, the networks in the rich regime exhibited a violation of the terminal item effect at a symbolic distance of 4, a somewhat surprising pattern that no models examined thus far have exhibited. The networks’ unconvential behavior was not due to the specific setup considered here: we observed similar behavior for alternative activation functions (Fig. S5), alternative loss functions (Fig. S7a), and deeper networks (Fig. S7b). Further, the networks exhibited even more overtly idiosyncratic behavior for larger numbers of items, one striking behavioral pattern being a periodic (rather than monotonically increasing) symbolic distance effect (Fig. S6). These findings indicate that, for TI, representation learning in standard neural network does not necessarily confer the benefits of representation learning suggested in prior work.

### 3.7 The rich-regime networks implement a cooperative code that lacks a transitive inductive bias

Given that the rich regime has been found previously to improve generalization on other tasks, we sought to understand why it yields anomalous behavior on TI. To this end, we leveraged previous work indicating that lazy and rich regimes implement different forms of norm minimization: the lazy regime minimizes the 𝓁_2_-norm of the network’s readout weights, whereas the rich regime approximately minimizes the 𝓁_2_-norm of all weights in the network [98–100; but see 84, 101]. For the networks studied here, we found that the norm of all weights in a fully trained network is dramatically smaller for smaller scales of initialization (Fig. 8a). To clarify why norm minimization over all network weights is associated with an anomalous inductive bias (unlike norm minimization over readout weights, analyzed in the previous sections), we directly analyzed the computations performed by the rich-regime neural network.

**Figure 8:**
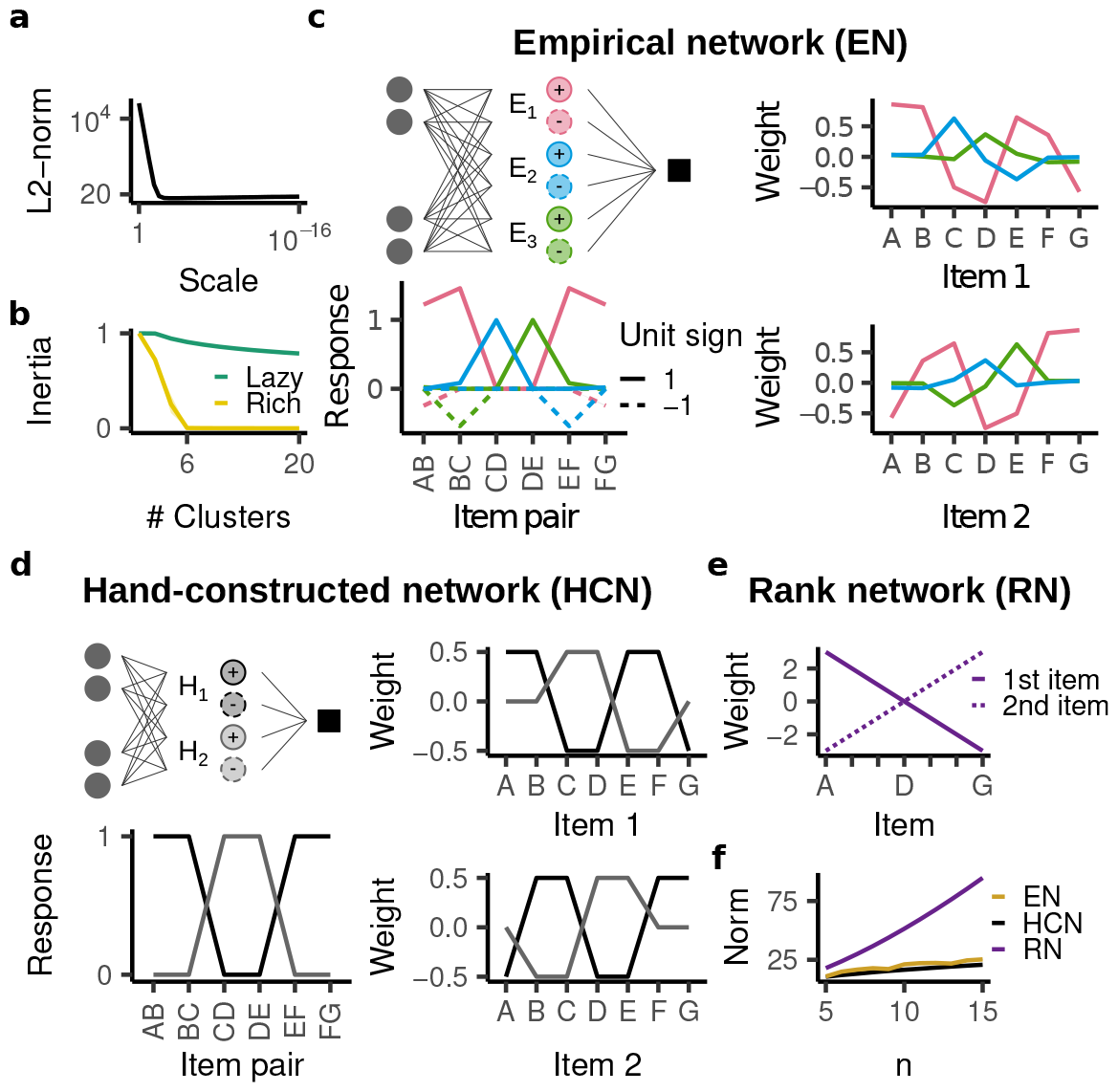
A mechanistic analysis of the rich regime’s inductive bias. All plots show the mean ± standard deviation across twenty random initializations. Standard deviation is too small to be visible. **a**, 𝓁_2_-norm of of all weights in the fully trained networks as a function of initialization scale. The rich regime yields much smaller norm. **b**, Inertia (i.e. proportion of explained variance) as a function of the number of clusters, for the lazy and rich network. Six clusters only leave 0.001% of the variance unexplained. **c**, The empirical network is therefore described by a network with six units, three with positive and three with negative responses. The right panels show the weights of the different units and the bottom panel shows how each units responds to the different training trials. Only units with positive readout weights are shown; the units with negative readout weights have the same structure but with Item 1 and 2 reversed. **d**, Analogous depiction of the hand-constructed network, which has four units. **e**, The rank network only has two units, but they span a much wider range. **f**, Weight norm of the empirical, hand-constructed, and rank network as a function of the number of items. The rank network has a much larger 𝓁_2_-norm, whereas the norm of the empirical network is similar to that of the hand-constructed network.

Prior studies [100, 102, 103] have found that minimization of the 𝓁_2_-norm over all network weights induces the networks’ hidden units to “specialize” into a low number of functionally distinct clusters with low 𝓁_2_-norm. To assess this possible structure, we performed k-means clustering with respect to the normalized weight vectors across all units, finding that in the rich regime, all 50,000 units of the network fall into just six clusters (Fig. 8b; details described in Appendix S3.3). In contrast, such a compact description was not apparent in the lazy regime. Remarkably, the six cluster centroids were extremely consistent between different random initializations (Fig. 8c).

We found that three of those centroids (“units”) were associated with positive readout weights, whereas the other three units were associated with negative readout weights. This is because the network has rectified activations in its hidden layer, which caused different units to specialize for trials with positive and negative labels. In examining hidden-layer weights, we focused on the positive units, which we denote by E_1+_, E_2+_, and E_3+_ (negative units are analogous, Fig. S10). Because we presented the network with concatenated onehot vectors, each weight entry corresponds to a different presented item and we identify each weight by its corresponding item. Note that items presented in the first and second position correspond to entirely distinct weights; we denote item *X* in the first or second position by *X*^(1)^ and *X*^(2)^, respectively.

We noticed two putative aspects of the underlying computation in the network: different units responded predominantly to a non-overlapping set of trials (“staggered response”; e.g. E_1+_ (pink) responded to AB, BC, EF, and FG, while E_2+_ (blue) responded to CD and E_3+_ (green) responded to DE; Fig. 8c, bottom left) and each unit encoded its corresponding trials by distributing positive weights across both items (“cooperative code”; e.g. to encode AB, E_1+_ assigned a positive weight to *A*^(1)^ and *B*^(2)^; Fig. 8c, right).

To see whether these two aspects could provide a sufficient explanation for the behavior in the rich-regime, we used them to hand-construct a simplified ReLU network (Fig. 8d). Specifically, the hand-constructed network has four hidden units, two with positive readout weights and two with negative readout weights. Again, we focus on those with positive readout weights, which we denote by H_1+_ and H_2+_ (see Fig. S9 for the analogous negative units). Note that we do not prove that the hand-constructed network actually learns the training trials with minimal 𝓁_2_-norm (though it has the lowest norm among all networks considered here). Rather, it serves as a useful construction to understand why the described computation yields both low norm and a non-transitive inductive bias.

Each unit in our construction implements a cooperative code, i.e. to encode its response to a trial, it distributes its weights equally between the two items. In particular, H_1+_ classifies trials AB and BC by assigning a weight of 0.5 to *A*^(1)^ and *B*^(2)^ as well as *B*^(1)^ and *C*^(2)^. Because of the positive weights associated with these items, H_1+_ would also respond positively to trials BA, CB, and DC. Preventing this positive response requires negative weights associated with items *A*^(2)^, *C*^(1)^, and *D*^(1)^. However, because a negative weight is already associated with *C*^(1)^ and *D*^(1)^, H_1+_ cannot classify trials CD and DE using a cooperative code. Thus, H_1+_ stays silent on these trials, which are instead encoded by H_2+_ (also using a cooperative code). This pattern explains the staggered responses observed in the empirical network. The interference from the negative weights associated with *E*^(1)^ and *F*^(1)^ prevents H_2+_ from classifying EF and FG, but these trials are no longer affected by interference with AB and BC and can therefore be encoded by H_1+_. For TI variants with more than seven items, the two units would continue alternating in the same way, giving rise to periodic network weights (Fig. S9).

Intuitively, the coding scheme amounts to a set of local rules that support learning of the training trials, but do not generalize to the test trials. For example, because the cooperative code implemented in unit H_1+_ encodes a positive response to trials AB and EF, it also results in an incorrectly positive response on EB (see Fig. S9).

Critically, the hand-constructed network has a lower 𝓁_2_-norm because the cooperative code keeps the network weights in a range from −0.5 to 0.5. In contrast, a network using a ranking system would require weights in a range from 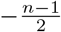 to 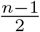 (Fig. 8e). As a result, its 𝓁_2_-norm is not only consistently larger, but also increases more quickly with increasing number of items (Fig. 8f).

Importantly, the staggered cooperative coding scheme is only norm-efficient because the hidden weights are followed by a rectifying nonlinearity. For example, H_1+_ takes on a value of −1 in response to CD. Without the rectification, H_2+_ would then have to compensate for the negative response to produce a positive label, which would require larger weights and thus would not be norm-efficient. However, because of the rectification following the hidden weights, H_1+_ remains silent on CD and requires no such compensation from H_2+_. This explains why a cooperative and staggered coding scheme has a low 𝓁_2_-norm when all weights of the network are trained, but not when only the readout weights are trained.

The hand-constructed network illustrates how a constraint that imposes a low 𝓁_2_-norm on all network weights can give rise to anomalous behavior on transitive inference. As noted above, the empirical network implemented the same computational principles as the hand-constructed network, albeit in a slightly different way: E_1+_ approximately implemented H_1+_, whereas E_2+_ and E_3+_ jointly approximated H_2+_. These differences resulted in a somewhat higher, but qualitatively similar norm in the empirical network (Fig. 8f; see Appendix S3.3.3). In particular, the norm of both the empirical and the hand-constructed networks is systematically lower than that of the rank-network (Fig. 8f). Remarkably, the mechanism implemented by the empirical network is consistent across not only different random initializations but also different numbers of items *n*: E_1+_ always approximates H_1+_, whereas E_2+_ and E_3+_ jointly implement H_2+_ (Fig. S9f).

Our findings in this section may appear contrary to those of Nelli *et al*. [30], who also studied TI performance of neural networks, but found that the rich regime did not impair transitive generalization and further yielded an explicit rank representation in the hidden layer. We found that this was caused by a weight symmetry imposed on the hidden layer that constrains their networks to an additive internal representation (see Appendix S2.6). Indeed, our clustering analysis revealed that these networks approximately implemented the rank network (Fig. 8e) and the 𝓁_2_-norm of their weights was much higher than that of the unconstrained neural network (Fig. S10a). This further illustrates the misalignment between the rich regime’s norm minimization and a transitive inductive bias: architectural constraints that improve transitive generalization (here, weight symmetry) lead to a higher weight norm.

## 4 Discussion

We found that standard statistical learning models can perform transitive inference (TI) and recapitulate three empirically observed behavioral patterns (symbolic distance effect, terminal item effect, and memorization effect). The behavior of a given model is sufficiently captured by a single scalar “conjunctivity factor” *α*, which characterizes the model’s internal representation (of task items) on a spectrum from fully compositional (*α* = 0) to fully conjunctive (*α* = 1). For *α* = 0, the model is constrained to encoding a transitive relation. For partly conjunctive representations (0 < *α* < 1), the model is not constrained in this way, but the principle of norm minimization nevertheless yields transitive generalization. For fully conjunctive representations (*α* = 1), the model cannot generalize transitively. Finally, we found that when representation learning is enabled in hidden layers, networks perform worse on TI and exhibit different behavioral patterns than living subjects. Through hand-constructed networks and a clustering-based analysis of empirical networks, we suggest that this anomalous behavior arises from the different form of norm minimization implemented by the rich regime.

Models of relational cognition often represent relations explicitly and are pre-configured to have a particular relational inductive bias [104–106]. In particular, alternative accounts of TI are either pre-configured to associate a rank (or value) with each item [29, 31, 49–51] or suggest that humans rely on abstract knowledge of transitivity [e.g. 31, 34, 107] (as implemented, for instance, in a cognitive map [108–110]). In contrast to these accounts, we took a “minimal principles” approach, representing nothing but the input itself (i.e. the two presented items). This perspective casts higher-level behavioral capacities (in this case transitive generalization) as emergent from minimally structured learning systems [111, 112]. Studies within this paradigm usually rely on simulations, which can leave the mechanisms for emergent behavior, such as generalization, unclear [cf. 113]. In contrast, our analytical account identifies the specific model components responsible for transitive generalization. This clarifies how the brain could implement relational generalizations without pre-configured representations or compositional constraints.

More generally, relational cognition likely relies on a wide range of learning mechanisms [114, 115]. The models considered here require repeated interleaved presentations of the training trials [116], suggesting learning mechanisms associated with prefrontal or higher-level association cortices. In contrast, other brain regions, such as the hippocampus, support rapid learning without the need for repeated trial presentations, presumably through the operation of memory re-activation (e.g. inferring that *B* > *D* by recalling that *B* > *C* and *C* > *D*) [117–120]. A recurrent neural network model with Hebbian plasticity (“*REMERGE*“) has been proposed to explain how a reactivation-based mechanism could support transitive generalization [121]. While this learning model does not recapitulate behavioral patterns such as the symbolic distance effect, it can explain how subjects may learn transitive inference from a minimal number of trials. Finally, whereas both the reactivation-based and statistical learning models described above rely on emergent inductive biases of the underlying learning mechanisms, learning systems can also develop relational inductive biases through structure learning or meta-learning [122, 123], which are associated with both hippocampus and prefrontal cortex [115, 124–126].

Which of these learning mechanisms is implicated in a particular TI task variant likely depends on factors such as the stimulus structure, how training trials are presented, and when test trials are presented. Subjects may also rely on a mixture of learning mechanisms on a single task. For example, trials could initially be encoded in the hippocampus but eventually be consolidated in prefrontal cortex [114, 127]. Creating more unified models of relational learning (e.g. fusing the *REMERGE* -model and our similarity-based mechanism, or incorporating structure learning) could shed further light on the interplay between different learning mechanisms on TI and their dependency on different task parameters such as stimulus presentation and training curriculum. Doing so may also allow closer investigation of proposed neural implementations of relational learning [117, 128].

With respect to similarity-based relational learning models, our account could be seen as an endpoint to a series of investigations of TI behavior: expanding upon previous studies [30, 53, 54], we show comprehensively that the principle of norm minimization enables any model with partly conjunctive representations to generalize transitively and further gives rise to naturalistic behavior on TI. Importantly, norm minimization is implicated not only in gradient flow and ridge regression (the examples we consider), but also a much broader range of learning models [129], including reinforcement learning [130]. Accordingly, the consistent behaviors many different animals exhibit on the task could be due to this shared, underlying learning principle. This is an alternative to the view that the ubiquity of TI stems from its ecological role in social cognition, in particular in animals which form hierarchies [131, 132], and the view that TI entails explicit reasoning.

Our results, while providing an alternative explanatory account of TI, should nevertheless be interpreted cautiously with regards to the basis of TI in living subjects, as our model is limited in a number of ways. In terms of behavioral predictions, it cannot account for the asymmetry effect, the observation that performance in living subjects is often better for items towards the start of the hierarchy (e.g. AB) than items towards of the end (e.g. FG) [29]. Further, our model does not generate predictions for what behavior we might expect from subjects that, after being trained on item pairs, are presented with three items at a time and expected to pick the largest item [33, 40, 133, 134]. Finally, our results do not speak to tasks testing for transitive inference in the form of a single question (e.g. “Alice is taller than Bob and Bob is taller than Chris. Who is taller, Alice or Chris?”) or after minimal training [26, 135]. Such a format likely requires a different mechanism from the one considered here and may be better understood as an instance of explicit reasoning.

These limitations, and extensions of TI, can inspire refining or expanding the statistical modeling approach. For example, Nelli *et al*. [30] found that conventional statistical learning models could not account for human behavior on a particular variant of TI [“list linking,” 136], but that a modification in which uncertainty is encoded could. More broadly, TI, with its rich and well-established set of empirical phenomena [e.g. 137–139], may serve as a useful model task to compare statistical learning models (and their failure modes) to human and animal behavior. For example, the lack of an asymmetry effect and the inability to perform the three-item task indicate what is lost by treating TI as a binary categorization task. A model that understands its decision as tied to one of the presented items (rather than as a choice between two arbitrary categories) may be able to better account for these behaviors [29]. Notably, better accounts of behavior within the TI framework may not only elucidate how subjects perform relational learning, but even inform more general models of statistical learning in living subjects.

Despite these potential limitations, our account still makes a set of falsifiable predictions that can produce evidence for or against the biological relevance of our insights. In particular, we identified the conjunctivity factor *α* as a broadly important parameter for TI task behavior. Our account would predict that changes in *α* should lead to a set of coordinated changes in behavior. For example, a stronger memorization effect should be associated with a weaker terminal item effect on the training cases (as both are caused by an increase in *α*). Observing that these different behavioral patterns indeed change in a coordinated manner (for example between different subjects performing the same task) may indicate that our theory can accurately describe behavior.

Changes in the conjunctivity factor may arise from inter-individual differences (different subjects may have representations that are best described by different values of *α*), but could also be introduced through experimental interventions. In particular, certain task variations may promote a representation of the input as either two separate items or a single conjunctive stimulus [140]; for example, presenting the two items in a common scene rather than as distinct stimuli may result in a higher *α*. More broadly, this suggests that the conjunctivity factor may be a useful parameter for comparing subjects and TI task variants.

Beyond behavioral predictions, the conjunctivity factor also suggests a new approach for clarifying the neural basis of TI. On the one hand, perturbations in relevant neural areas, for example through lesions [78], may affect the conjunctivity factor. On the other hand, using neural recordings to estimate the empirical representational similarity [141] between distinct, overlapping, and identical trials could enable the estimation of an effective conjunctivity factor in a particular neural area. If differences in this estimated conjunctivity factor (either due to inter-individual differences or as a result of lesions) are associated with the corresponding behavioral differences predicted by our model, this may suggest a role for statistical learning as described here, and, in addition, that the recorded neural area is indeed involved in TI.

Intriguingly, past work suggests that a conjunctivity tradeoff – manifesting across different tasks – can be linked to a specific brain region. Lesions of the hippocampus have been found to impair generalization (but not the learning of training trials) in transitive inference (142; but see 143) and, in separate work, accelerate learning of transverse patterning [144], two effects that would both arise from an increased conjunctivity factor in our model. Notably, other studies indicate that hippocampal lesions can also impair learning of transverse patterning [145–147]. These differences appear to depend on whether items are presented together (e.g. together on a screen), in which case the hippocampus was not required for transverse patterning, or apart (e.g. in separate containers), in which case hippocampus was required. These results broadly suggest a role for hippocampus both in building representations of items presented separately [140] and in limiting the conjunctivity of items presented together.

Moreover, our findings on rich-regime neural networks performing worse on transitive inference may have important implications for machine learning, as deep neural networks have been observed to struggle with tasks involving compositional generalization more broadly [148–154]. A common strategy in attempting to address these shortcomings consists in training ever larger models on ever larger datasets [155]. Such models have shown impressive results on tasks such as natural language production [156, 157], but the overall scale and complexity of the training data, model, and learning algorithm make it difficult to attribute successful or unsuccessful generalization behaviors to specific components of the model [though see 158–161]. In contrast, TI is a particularly simple relational task on which deep networks exhibit non-naturalistic behavior. Our findings explain why the rich regime, which has a useful inductive bias on many different tasks, can give rise to such anomalous behavior on transitive inference. This can help in designing principled modifications to prevent such anomalous behavior – for example by adding additional training trials, regularizing the hidden layer to prevent a local encoding mechanism, or changing the connectivity structure of the network – and in this way improve our understanding of how relational learning is implemented [162–165]. Notably, the tools we employ to analyze the rich-regime behavior may also prove useful for analyzing neural networks trained on other tasks. At the same time, our analysis here was primarily empirical, but future work could provide an analytical characterization of the rich-regime solution [99, 100, 166].

Finally, transitive inference is a simple instance of compositional generalization, i.e. the ability to conceptualize prior experience in terms of components that can be re-configured in a novel situation [167, 168]. More broadly, compositional generalization has long been understood to be a crucial component for human-like learning and generalization [163], in large part due to the diversity and breadth of its applications: compositional generalization often involves the composition of many rules, relations, or attributes [163, 169–171]. The comparative simplicity of TI enabled us to identify how minimally structured learning systems can implement the inductive biases needed for this task. Our analysis provides a case study for how standard statistical inductive biases determine behavior on compositional tasks, in particular clarifying how representational structure (conceptualized through the conjunctivity factor *α*) impacts compositional generalization [172]. At the same time, TI certainly does not capture the complexity of most compositional task paradigms and future work should extend our formal analysis to a broader scope of compositional tasks. This would clarify the conditions under which standard statistical learning principles can explain compositional generalization and clarify challenging compositional motifs that require additional learning mechanisms.

In summary, we here investigated transitive inference, a task that tests for a fundamental logical capacity and has fascinated researchers across neuroscience and psychology for many decades. We derived exact mathematical equations describing the TI behavior of a large class of statistical learning models. This allowed us to understand systematically how these models can generalize compositionally, using TI as an example. Our theory provides a basis for using TI to investigate the neural structures implementing relational cognition. At the same time, our findings also suggest that TI, in its standard form, may not be sufficient to clarify relational inference abilities that require more than statistical learning. For this purpose, other relational tasks should be considered.

## Acknowledgments

We are grateful to the members of the Center for Theoretical Neuroscience at Columbia University, the SueYeon Chung lab at Flatiron/NYU, and attendees at Cosyne 2023 and the Gatsby Tri-Centre Meeting 2023 for helpful comments. We thank Taiga Abe, Veronica Bossio, Tala Fakhoury, Ching Fang, Jeff Johnston, Erica Shook, Sharon Su, and Denis Turcu for detailed feedback. The work was supported by NSF 1707398 (Neuronex), Gatsby Charitable Foundation GAT3708, and NIH-R01MH111703.

## S1 Mathematical companion

In this section, we prove the mathematical statements made in the main text and provide additional details on the analysis of model behavior.

- Appendix S1.1 extends the analysis of additive models presented in Section “An additive representation yields transitive generalization” (p. 3).
- Appendix S1.2 defines the conjunctivity factor and determines the possible range of values it can take on.
- Appendix S1.3 determines the conjunctivity factor of a broad range of possible networks, clarifying the connection between network archiectures and TI behavior.
- Appendix S1.4 characterizes the TI behavior of models implementing norm minimization, gradient descent, and ridge regression.
- Appendix S1.5 uses this theory to predict the behavior of neural networks in the lazy regime.
- Appendix S1.6 analyzes the behavior of models trained through gradient flow on transverse patterning.

To leave the section relatively compact, we outsource the details for some of the more involved proofs to Appendix S4.

### S1.1 Models with additive representations and choice bias can make mistakes in TI but generalize correctly on average

Section 3.1 considered a model with no choice bias (i.e. *f* (*X, X*) = 0). Here we discuss the possible ramifications of a choice bias in additive representations. As a reminder, we consider a model

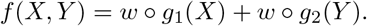

We can understand the model’s decision making in terms of two “implicit” ranks

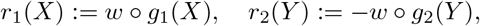

so that

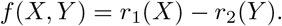

*f* will classify adjacent items *i* and *i* + 1, *i* ∈ {1, …, *n* − 1}, correctly if and only if those ranks satisfy

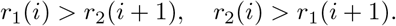

We can characterize this condition more easily by defining two sequences: one starting at *i* = 1 with *r*_1_ (we call this one *s*_1_) and one starting at *i* = 1 with *r*_2_ (we call this one *s*_2_). At each item those sequences flip whether they specify the rank of *r*_1_ or *r*_2_. That i s: *s*_1_ specifies *r*_1_ at odd items and *r*_2_ at even items and *s*_2_ behaves in the exact opposite way:

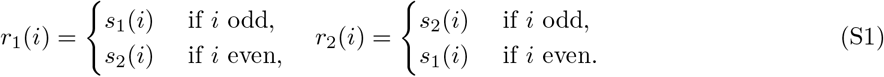

By construction, *f* will classify adjacent items if and only if both *s*_1_ and *s*_2_ are monotonically decreasing. For items *i, j* with an odd symbolic distance, this immediately implies that *f* must also correctly order those, as *r*_1_(*i*) and *r*_2_(*j*) as well as *r*_2_(*i*) and *r*_1_(*j*) will be assigned to the same sequence and therefore be monotonic. For items with an even symbolic distance, however, the two ranks will be mapped to different sequences. The models do not have to generalize transitively and can generalize pairs of items incorrectly for arbitrarily large symbolic distances.

We leave with two notes. First, the average margin

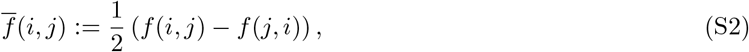

has no choice bias even when *f* itself does (i.e. 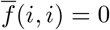). As a result, this average margin still generalizes transitively even when *f* itself does not in generalize. Second, if the model classifies pairs (*i, j*) correctly for some even symbolic distance, it must also do so for any higher symbolic distance.

### S1.2 The conjunctivity factor

We consider a general representation

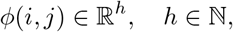

where *i* and *j* denotes the numerical rank of the two presented items *X, Y*. We then consider the representational similarity (measured by the dot product) between two trials (*i, j*) and (*i*^′^, *j*^′^):

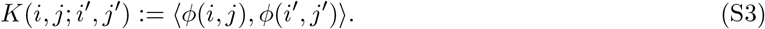

We assume that there are three values *K* can take, *κ*_*s*_ if *i* = *i*^′^ and *j* = *j*^′^, *κ*_*o*_ if either *i* = *i*^′^ or *j* = *j*^′^, and *κ*_*d*_ if *i* ≠ *i*_′_ and *j* ≠ *j*_′_. *α* is then defined as

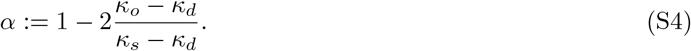

We assume *κ*_*s*_ ≠ *κ*_*d*_. If *κ*_*s*_ = *κ*_*d*_, then *κ*_*s*_ = *κ*_*o*_, meaning that *ϕ*(*i, j*) is constant and unable to learn anything.

#### Lemma S1.1

*α is restricted to the range* [0, 3]. *For arbitrary n, α can take on values in the range* [0, 1].

*Proof*. For compactness, we denote *κ*_*o*_ by *o, κ*_*d*_ by *d*, and *κ*_*s*_ by *s*. We then consider the Gram Matrix of all possible combinations of the first three items, i.e.

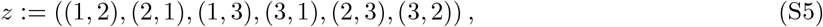

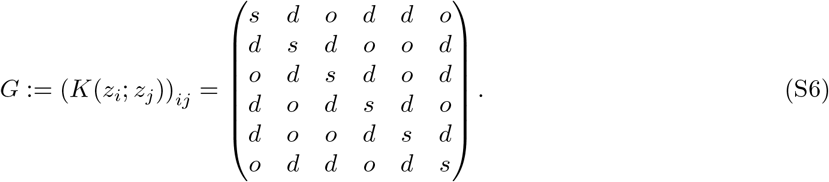

*G* is positive definite (as it is a Gram matrix), implying that all its eigenvalues are positive.

Using Mathematica, we determine two of these eigenvalues as

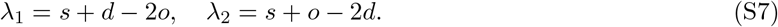

We can transform

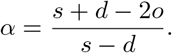

As *s* − *d* ≥ 0 (by basic properties of a kernel), *λ*_1_ ≥ 0 implies that *α* ≥ 0. Further, *λ*_2_ ≥ 0 implies that

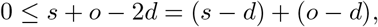

and therefore

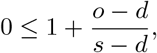

implying that *α* ≤ 3. Finally, *α* can take on arbitrary values in [0, 1] as *α* = 0 corresponds to an itemic encoding and *α* = 1 corresponds to a onehot encoding. By linear combinations of those encodings, we can achieve arbitrary values in-between. □

#### Remark S1.2

*The lemma leaves open which values in the range* (1, 3] *are attainable. In Appendix S4*.*2, we provide a construction that can attain α* > 1 *for all n. Notably, this indicates a feature basis that encodes overlapping item pairs as less similar than distinct item pairs. For n* = 3 *attains α* = 3. *As n increases, however, α decreases, gradually approaching* 1. *It remains an open question whether there are features that, at larger n attain higher values for α than our construction*.

In subsequent sections, we will constrain ourselves to *α* ∈ [0, 1].

### S1.3 Impact of nonlinearity, depth, and connectivity structure on a deep neural network’s conjunctivity factor

We here compute the conjunctivity factor for a broad range of hidden representations, to better understand how it is impacted by different choices in architecture.

#### S1.3.1 Densely connected neural networks

This corresponds to a standard densely connected neural network with layerwise weights *W*^(*l*)^:

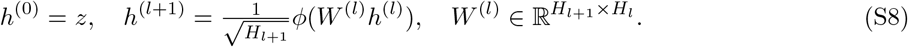

where *ϕ* is some nonlinearity. We assume that these weights are sampled from a rotationally invariant distribution with some variance 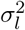 (e.g. a normal distribution). Further, we assume that *ϕ* is structured as

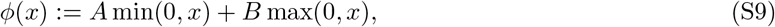

which has as special cases ReLU [*A* = 0, *B* = 1, 173], leaky ReLU [*A* < 1, *B* = 1, 174], and the absolute value (*A* = −1, *B* = 1). As shown below, the conjunctivity factor of *h*^(*l*)^ is invariant to *ϕ*’s “magnitude”

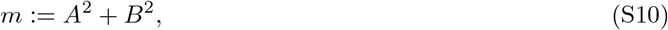

but depends on its “relative slope”

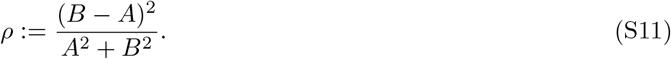

In particular, *ρ* = 1 for ReLU. (Note that similar analytical expressions could be derived for a broad range of other nonlinearities.)

We are now interested in the trial-by-trial similarity in each network layer. The expected similarity (which the empirical similarity will converge to in the large-width limit) is

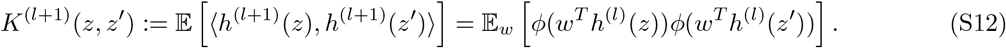

Our goal is to express this expression in terms of *K*^(*l*)^, the kernel of the previous layer in order to derive a recursive formula.

Notably, *ϕ* is homogeneous and therefore

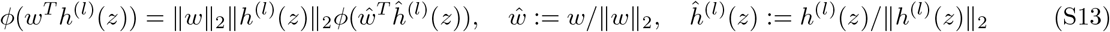

As a result,

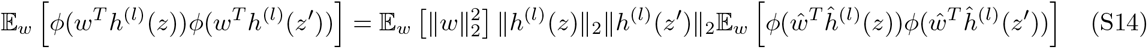

Notably, prior work [71, 175–177] has found that

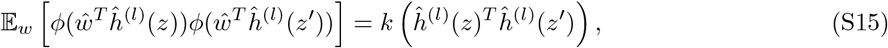

where

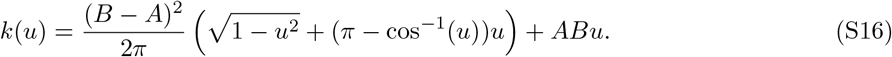

This means that

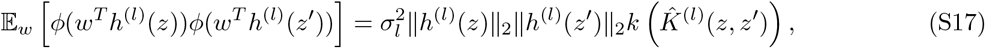

where

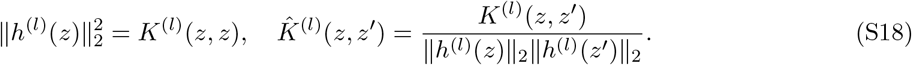

Defining

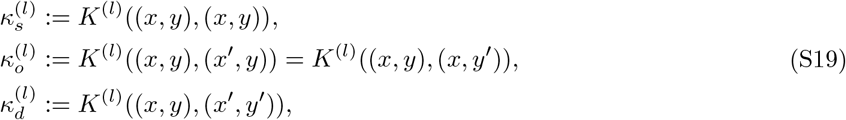

we infer

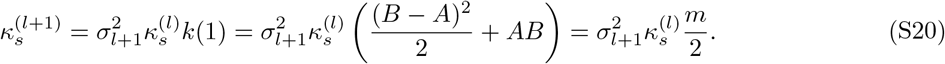

Further,

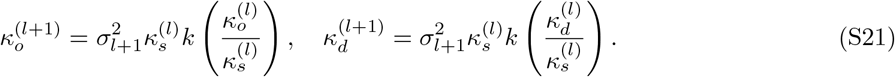

Defining

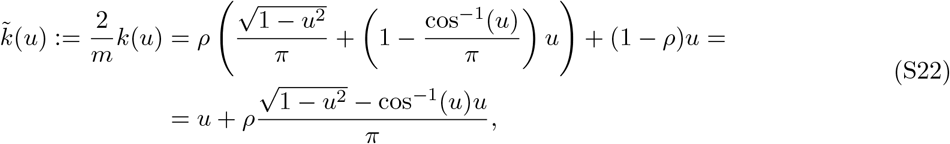

and

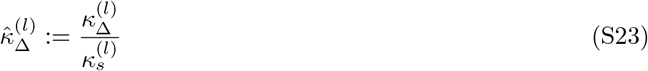

we infer

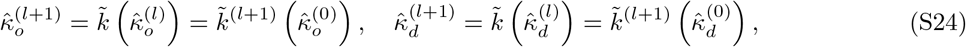

where 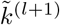 denotes *l* + 1 applications of 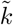. This allows us to recursively compute

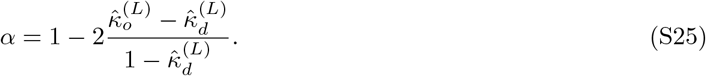

As a special case, we consider input items *x*, which are all orthogonal to each other, meaning that

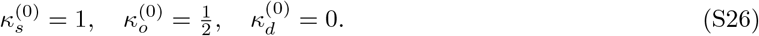

and compute the similarities of the first hidden layer:

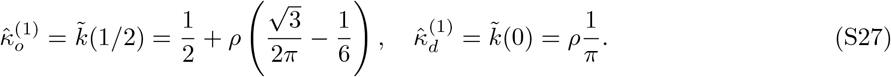

The conjunctivity factor *α* in this case is given by

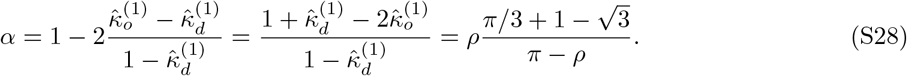

For ReLU (i.e. *ρ* = 1), *α* ≈ 0.15.

#### S1.3.2 Neural networks with additive nodes

We can now extend these formulas to networks with partly additive nodes. In particular, we assume that the fraction (1 − *λ*) is fully connected and the fraction *λ* is only connected to one of the two items, though followed by the same nonlinearity. This part of the representation has the similarities

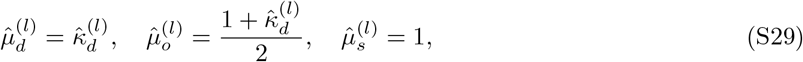

and so the new normalized dissimilarities are given by

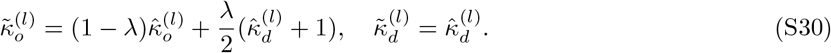

This, in turn, means that

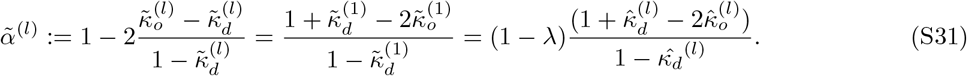

Predictably, for a single layer, the additive units therefore simply reduce the conjunctivity factor by a proportional factor. For multiple layers, this reduction has ramifications on the densely connected units as well, as these units draw upon both the additive and densely connected units. In fact, in the limit of infinite depth, any neural network with some proportion of additive units will approach a fully compositional representation:

##### Lemma S1.3

*For a nonlinearity ϕ with relative slope ρ and a proportion of additive units λ, the infinite-depth conjunctivity factor*

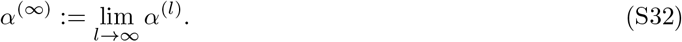

*is given by*

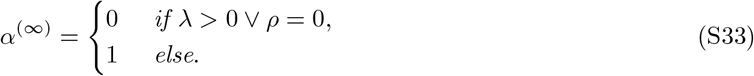

*Proof*. We prove the lemma in Appendix S4.5. □

### S1.4 Standard statistical learning models have a transitive inductive bias

In this section, we derive the TI behavior of different standard statistical learning models. Appendix S1.4.1 describes the general setup of transitive inference and our learning models. Appendix S1.4.2 then provides a general intuition for why these models implement an emergent ranking system and how we can compute these ranks. The subsequent sections go on to characterize model behavior for ridge regression (Appendix S1.4.3) and gradient flow (Appendix S1.4.4).

#### S1.4.1 Setup

We consider *n* ∈ ℕ items 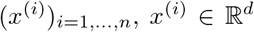. The task presents two of those items together and we denote this by

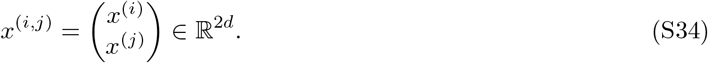

The corresponding label is given by

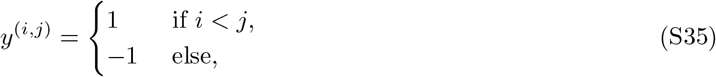

and the training dataset consists of all adjacent items, i.e.

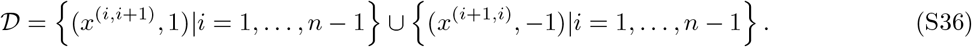

In matrix form, we denote this dataset as

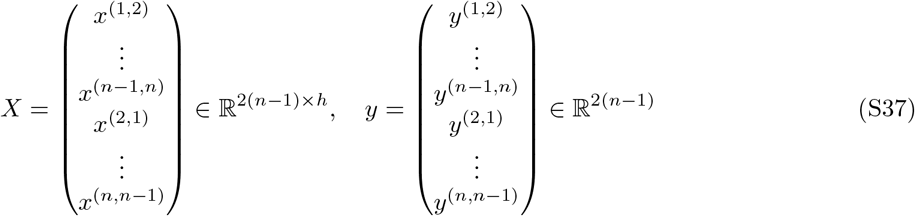

We consider a set of features

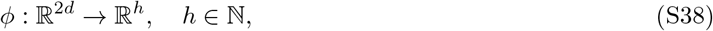

and a linear model of those latent features that predicts the output, i.e.

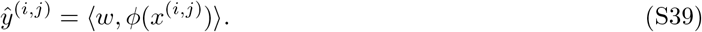

#### S1.4.2 General intuition

Many standard statistical learning models (in particular the ones we consider below) only depend on their input representation through its trial-by-trial similarity (or induced kernel, see also Appendix S1.2, 178):

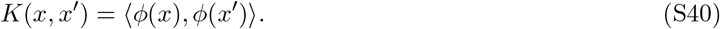

In particular, the minimal norm solution (described in Section 3.3), can be described in terms of the “dual coefficients”

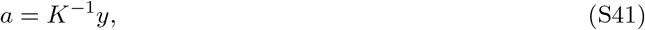

where *K* is the trial-by-trial similarity of the training dataset:

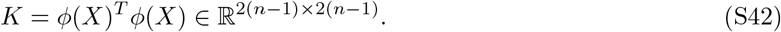

Each dual coefficient corresponds to a particular training data point (*i, j*) and we denote it by *a*_(*i,j*)_. The model behavior on a new data point *x*^(*i,j*)^ is then given by

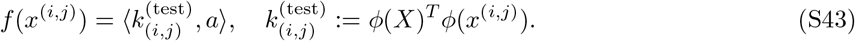

Importantly, the similarity between test data points and training data points can only take on two possible values: *κ*_*o*_ and *κ*_*d*_. Specifically, the dual coefficients *a*_(*i,i*+1)_, *a*_(*i,i*−1)_, *a*_(*j*−1,*j*)_, and *a*_(*j*+1,*j*)_ are overlapping with the test input (*i, j*) and therefore have similarity *κ*_*o*_. (Note that if *i, j* ∈ {1, *n*}, some of these dual coefficients will not exist. To handle this special cases, we simply set the corresponding dual coefficients to zero. For example, if *i* = 0, *a*_(*i,i*−1)_ = *a*_(1,0)_ corresponds to a non-existent data point and is set to zero.) All other data points have similarity *κ*_*d*_. Taken together, we can express model behavior as

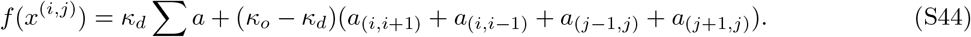

Crucially, this behavior is a sum of values dependent on *i* and values dependent on *j* — just like we observed for a purely additive representation. This immediately implies that the model implements a ranking system. Specifically we define

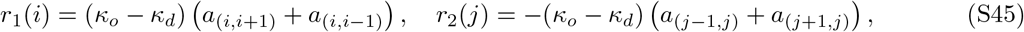

yielding

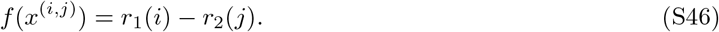

To see whether the model generalizes, we need to determine whether the ranks are ordered correctly. To do so, we analytically solve the inverse problem Eq. (S41). This is described in detail in the next section, which considers a generalization of the problem considered here. As a brief summary, we rely on previous results on inverses for banded diagonal matrices [179] to compute the inverse matrix. This inverse is expressed in terms of hyperbolic functions. We then use hyperbolic identities in a few different ways to compute *a* and, finally *r*_1_ and *r*_2_ from this. This reveals that *r*_1_(*i*) = *r*_2_(*i*) and further gives rise to the analytical expression provided in the main text.

#### S1.4.3 Ridge regression

Here we assume that the weights are learned through ridge regression. Specifically, our loss function is given by the mean squared error over the training set:

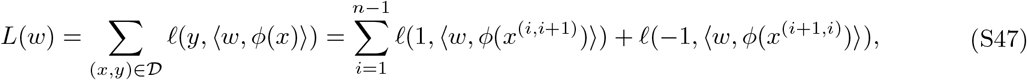

where 𝓁(*y, ŷ*) = (*y* − *ŷ*)^2^. The weights *w* are then determined by minimizing the optimization problem

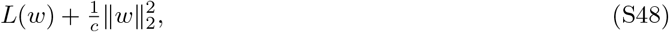

where *c* is the inverse strength of the regularization. The corresponding dual problem is given by

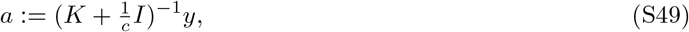

where *K* is the cross-sample similarity,

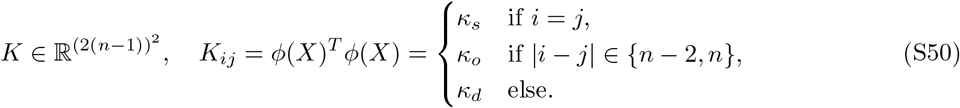

Note that for 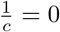, we obtain the special case from the previous section (Eq. (S41)). Writing *a* as

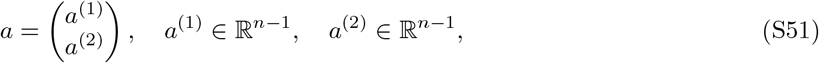

we can write 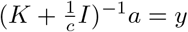 as

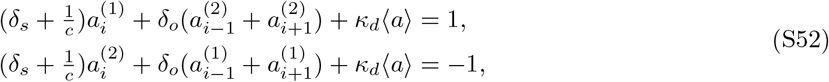

where we define *δ*_*s*_ = *κ*_*s*_ – *κ*_*d*_, *δ*_*o*_ = *κ*_*o*_ – *κ*_*d*_, 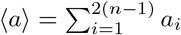 and, for notational simplicity, 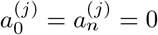. This means that if *a*^(1)^, *a*^(2)^ are a solution to the equation, then so are

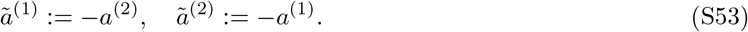

Since (S49) is well-defined, the solution *a* must be unique and we can infer

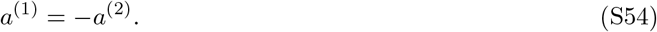

We thus define

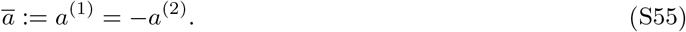

Setting

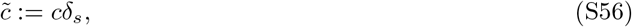

(S52) yields

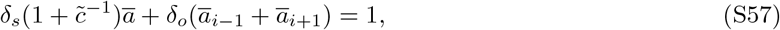

as ⟨*a*⟩ = ⟨*a*^(1)^⟩ + ⟨*a*^(2)^⟩ = 0 and therefore the last summand vanishes. We then divide by *δ*_*s*_ on both sides; as 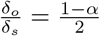, this results in

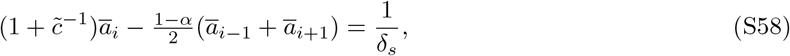

In matrix form, this results in

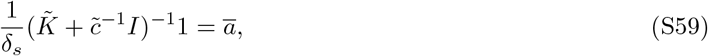

where

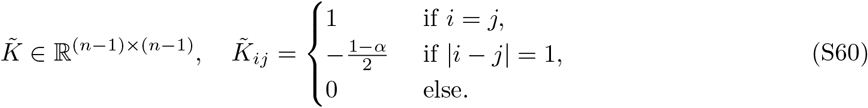

The predictions of the fitted model can be expressed in terms of the dual coefficients by

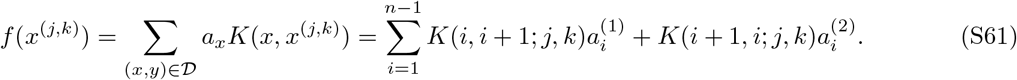

We can express this in terms of *ā* as

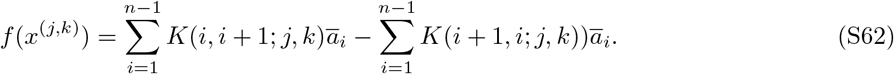

There are now two different cases: either *x*^(*j,k*)^ is in the training data or not.

**Case 1: Test data**. If *x*^(*j,k*)^ is not in the training data, all data points are either overlapping or distinct. In the first data point, a data point is overlapping if *i* = *j* or *i* = *k* − 1 and therefore

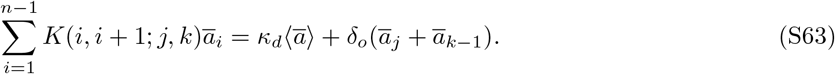

(Note that one of these dual coefficients, as in the previous section, may not exist, if *i* ∈ {1, *n*}. To leave the expression simple, we define *ā*_0_ = *āa*_*n*_ = 0, covering these cases as well. Analogously, the second sum amounts to

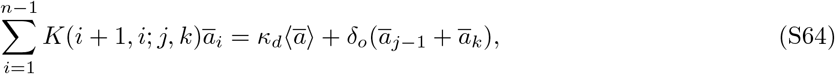

and overall, the model behavior is expressed as

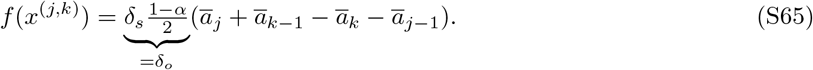

**Case 2: Training data**. If *x*^(*j,k*)^ is in the training data, then either *k* = *j* + 1 or *k* = *j* − 1. In the first case, the first sum, has as one of its data points (*j, k*) = (*j, j* + 1) itself. As it has no overlapping data points, this sum is expressed as

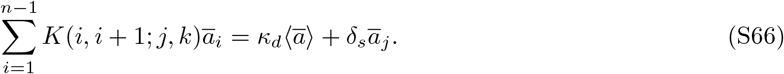

The second sum still has two overlapping data points as in the previous case and as a result, model behavior is given by

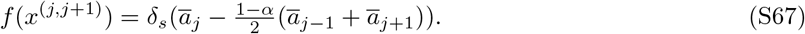

The second case (*k* = *j* − 1) works analogously.

Taken together, the prediction made by *ā* are expressed as

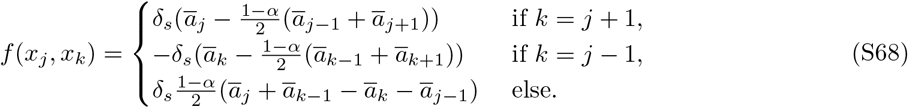

In particular, this makes apparent the emergent rank representation

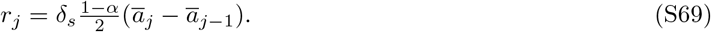

To compute the specific rank representation, we need to invert 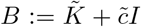. *B* is a tridiagonal matrix with diagonal 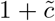 and off-diagonal 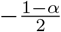. The ratio between diagonal and off-diagonal is therefore given by

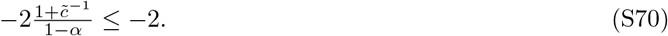

As this ratio is smaller than −2, we can apply Theorem S4.1 to infer

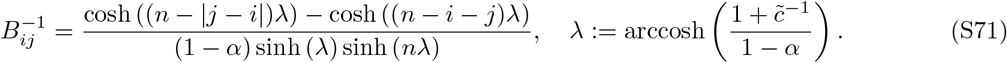

We can then use this to compute *δ*_*s*_*ā* = *B*^−1^1. Using hyperbolic trigonometric identities, we can simplify these expression further, resulting in the following lemma:

##### Lemma S1.4

*For non-adjacent items*,

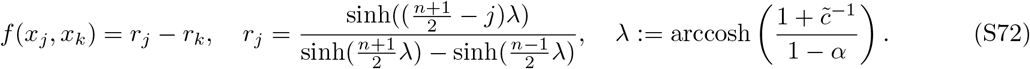

*For adjacent items*,

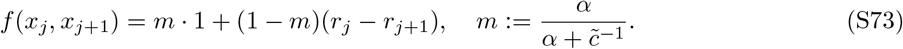

*Proof*. See Appendix S4.3.

#### S1.4.4 Gradient flow dynamics

We now consider the case where the weights *w* are trained using gradient flow rather than ridge regression. Specifically, the loss function consists in

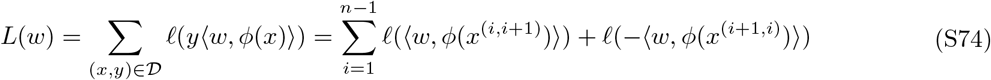

and weights are updated using gradient flow:

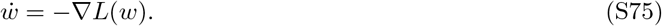

Note that in contrast to the previous expression (Eq. (S47)), here we express the loss function as a function of one variable 𝓁(*yŷ*). This is possible because *y* is a binary label, so the mean squared error, for example, turns into

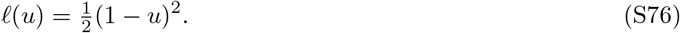

More generally we consider any convex, montonically decreasing loss function 𝓁(*y ŷ*). Other examples include the crossentropy

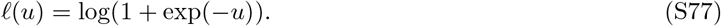

To simplify our notation, we will denote

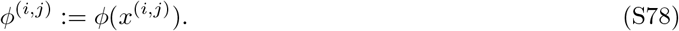

Our loss function is then given by

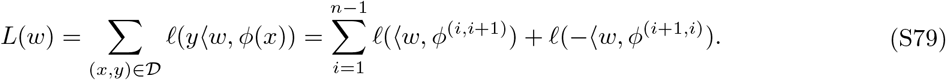

We now first consider the general case explaining how gradient descent gives rise to a terminal item effect and, in turn, how a terminal item effect on the training trials is connected to successful generalization on test trials. We then make more precise statements regarding the case of mean squared error.

##### General loss functions

Setting

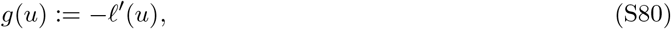

we write

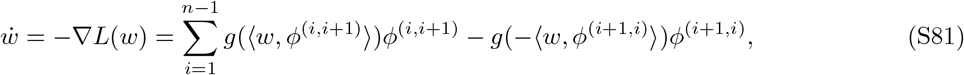

and so writing

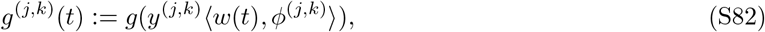

we can express

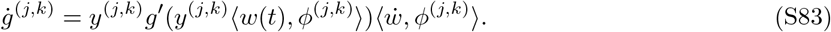

We now aim to express this differential equation purely in terms of *g* (rather than *ϕ*). To do so, we consider the intercept

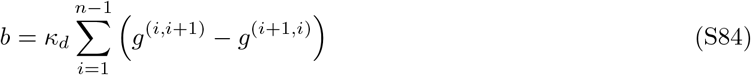

Moreover, we define

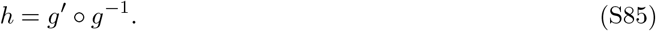

Using Eq. (S81), we express

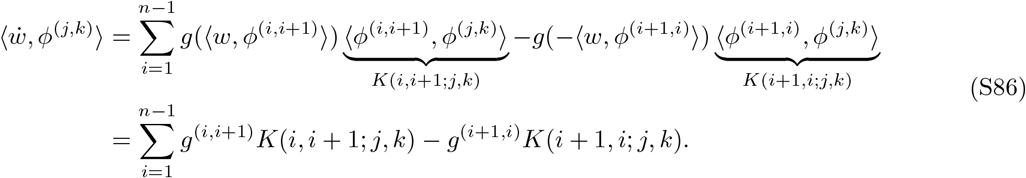

This allows us to express *ġ* as dependent on *g* and the kernel *K*:

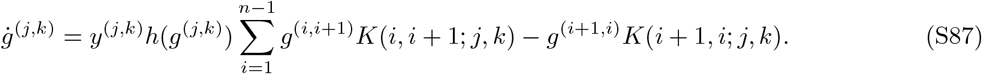

First, consider *k* = *j* + 1. By the same logic as in the previous sections, the first sum has one identical data point and all other data points are distinct, whereas the second sum has two overlapping data points:

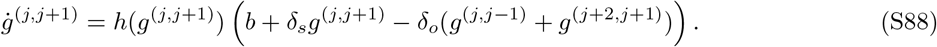

As 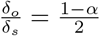, this implies

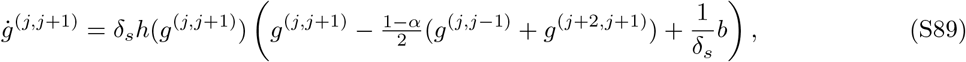

where we define as boundary conditions *g*^(0,1)^ ≡ *g*^(1,0)^ ≡ 0 and *g*^(*n,n*+1)^ ≡ *g*^(*n*+1,*n*)^ ≡ 0. Analogously, for *k* = *j* − 1,

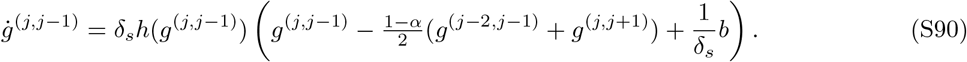

At initialization, *w* = 0 and therefore,

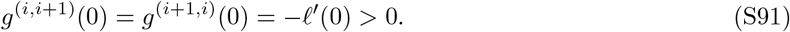

The differential equations above imply that the equality *g*^(*i,i*+1)^(*t*) = *g*^(*i*+1,*i*)^(*t*) keeps holding and we define

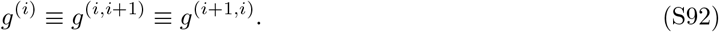

This implies *b* = 0 and we can express

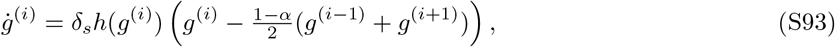

with the boundary conditions *g*^(0)^ ≡ *g*^(*n*)^ ≡ 0. At initialization, *g*^(*i*)^(0) = −𝓁^′^(0) > 0, as 𝓁 is monotonically decreasing. As 𝓁 is convex, *h* is strictly negative. If *α* = 0, at initialization, for *i* = 2, …, *n* − 2,

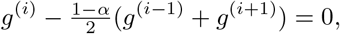

As a result, the gradient of all intermediate items initially remains unchanged. The boundary conditions break this symmetry for the terminal items *g*^(1)^ and *g*^(*n*−1)^, which immediately decrease. Once they decrease, the gradient of their neighboring items, *g*^(2)^ and *g*^(*n*−2)^ will also become negative and in this way, the changes in *g* percolate from outside in. This causes a terminal item effect.

For *α* > 0,

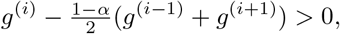

all *g*^(*i*)^ decrease from the start. However, the terminal items *g*^(1)^ and *g*^(*n*−1)^ decrease faster, and the faster decreases gradually percolate in as well. As a result, for higher *α*, the terminal item effect is more attenuated, but nevertheless still obtains.

On the other hand, for | *j* − *k* | > 1, all sums in Eq. (S86) consist only of overlapping and distinct data points. The distinct data points (consolidated in *b* = 0) cancel out, leaving us with

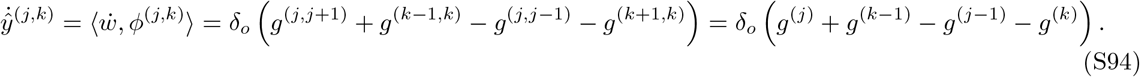

As 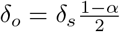, this implies

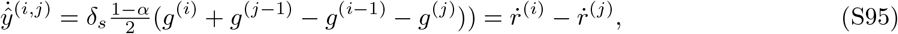

where we define the rank representation as

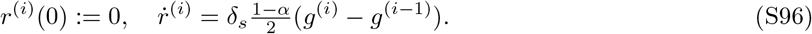

The terminal item effect implies that this rank representation will be increasing for the 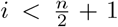 and decreasing for 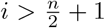. Further, the rank representation will be monotonic in *i* if *g*^(*i*−1)^ − *g*^(*i*)^ has higher magnitude for more terminal *i*. Remarkably, this means that a sufficient condition for the emergence of a rank representation from gradient descent is given by the observation of a convex terminal item effect throughout training.

##### Mean squared error

For the mean squared error,

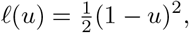

the dynamics are simplified as

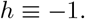

As a result,

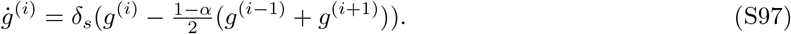

Defining the vector

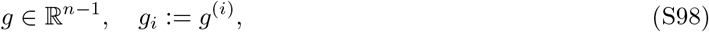

these dynamics can be written as

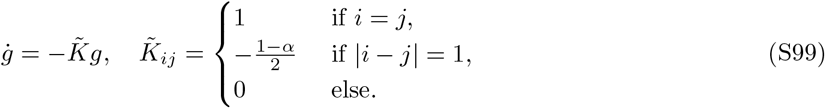

Further, as *g*(*u*) = 1 − *u*,

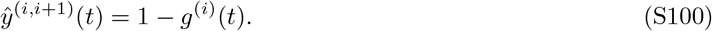

We determine *g*(*t*) by using the eigenvalue decomposition of *K*, as provided by Theorem S4.2. After some algebraic manipulation, this yields the following formulas for *g*_*j*_ and *r*_*j*_:

###### Lemma S1.5

*Given the mean squared error as a loss function, g*_*j*_ *and r*_*j*_ *amount to*

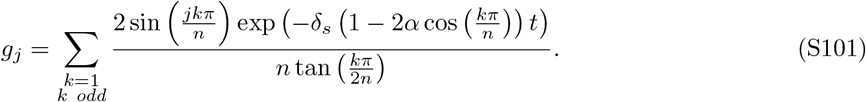

*and*

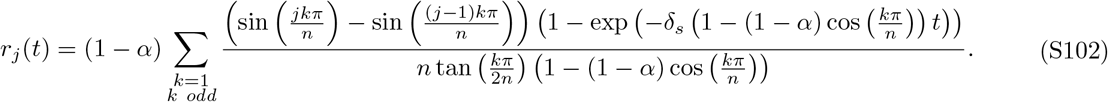

*Proof*. See Appendix S4.4.

We can further simplify this solution for *t* → ∞:

###### Lemma S1.6

*Given the mean squared error as a loss function*,

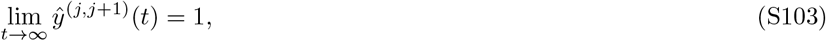

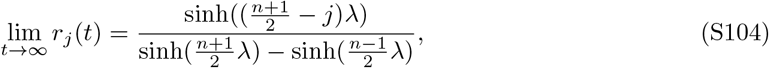

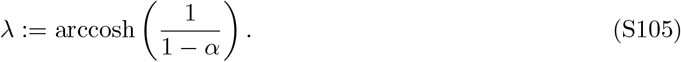

*Proof*. The solutions found by gradient descent and ridge regression (see below) converge to the same limit as *t, c* → ∞. The lemma thus follows directly from Lemma S1.4 below.

###### Remark S1.7

*Models are often trained on crossentropy for classification problems. While a closed-form solution to the learning dynamics cannot be determined, we can characterize its infinite-time solution. First, we note that crossentropy cannot be minimized by any finite value, but rather attains its minimum as u* → ∞. *As a consequence, the weights w diverge:* 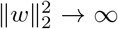. *However, we can normalize by the minimal margin. This normalized model converges to the support vector machine solution* [*19*]. *As all* 2(*n* −1) *training cases are support vectors for both the regression and classification problem, support vector regression and classification are equivalent and the normalized model is characterized by the lemma above*.

### S1.5 Exact learning dynamics of neural networks trained with backpropagation in the lazy regime

#### S1.5.1 Neural tangent kernel

We consider the deep network

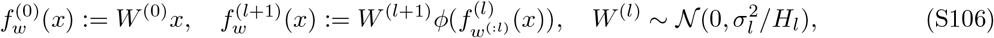

where

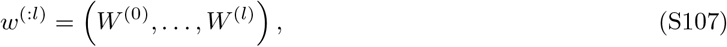

is the sequence of weights up to the *l*-th layer. We denote

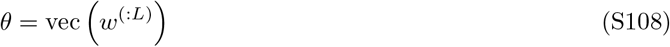

as the vectorized total parameters of the network, writing 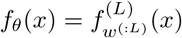. The *neural tangent features* are given by the linearization of the network around its initial parameters *θ*_0_,

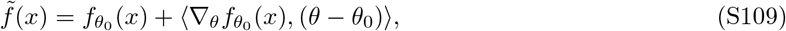

and the *neural tangent kernel* is given by the similarity according to those features:

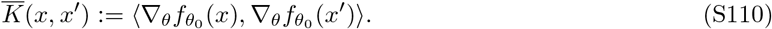

If the distance 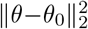 remains infinitesimally small throughout training, the linear approximation 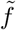 remains valid, and backpropagation can be approximated by gradient descent on the linear readout model given above. By computing the conjunctivity factor associated with 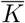, we can therefore predict the behavior of neural networks in the regime where the approximation is valid (the “lazy” regime). We make this computation in the next section, but previously remark on our choice of initialization.

##### Remark S1.8

*We here scale the weight variance by the input dimensionality, H*_*l*_, *rather than the output dimensionality, H*_*l*+1_. *This initialization is standard in practice as it conserves the variance of the network output throughout the layers with adequately chosen* 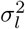 [*180*]. *In investigations of the neural tangent kernel, weights are frequently scaled by the output dimensionality instead. Part of the reason is that, as we will see below, standard scaling leads to less interesting behavior in wide neural networks. We here use the neural tangent kernel to show that our theory captures the behavior of neural networks in the lazy regime. For alternative initialization scales, this would still be the case, though the conjunctivity factor may be different*.

#### S1.5.2 The NTK’s conjunctivity factor

We consider a neural network with one hidden layer. Note that

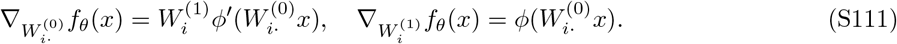

As a result,

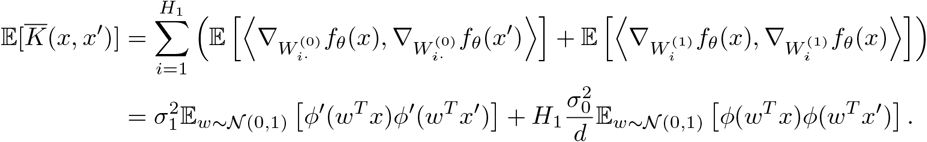

This means that as *H*_1_ grows large, this similarity diverges. Normalizing the similarity by *H*_1_, in turn, highlights that it is dominated by the similarity of the hidden layer. As a result, under standard initialization, the conjunctivity factor of the neural network with one hidden layer trained with backpropagation is equal to the conjunctivity factor of the hidden layer of the neural network.

In a sense, backpropagation is “trivial” in this regime as it is equivalent to training a linear readout. This is the origin of the different initialization paradigm highlighted in Remark S1.8. If the variance of *W*^(0)^ is also scaled with *H*_1_, the first term and the second term have comparable magnitude. While we refrain from computing the conjunctivity factor in that case, we note that similar analytical solutions can be derived and the resulting conjunctivity factor obeys the same qualitative trends observed in the previous section, though it is generally slightly higher.

### S1.6 Exact learning dynamics of models with fixed representations on transverse patterning

To illustrate the impact of different possible representations on tasks other than transitive inference, we consider the transverse patterning task. In this case, the training dataset is given by

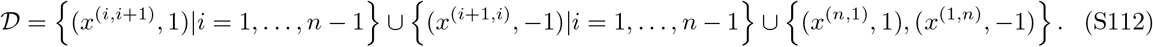

Thus, the loss function is given by

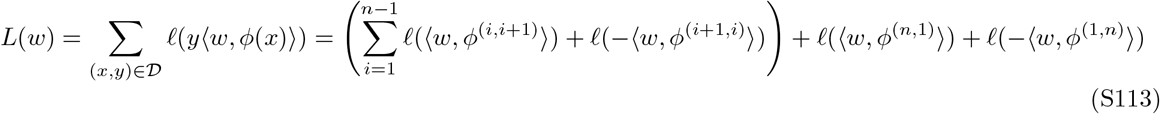

To write this more compactly, we define *ϕ*^(*n,n*+1)^ := *ϕ*^(*n*,1)^ and *ϕ*^(*n*+1,*n*)^ := *ϕ*^(1,*n*)^ (i.e. consider modular addition). We consider the mean squared error and set

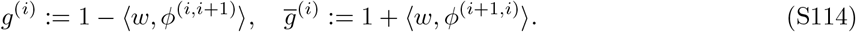

Gradient flow is then given by

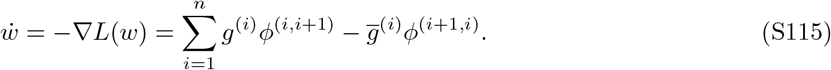

Thus,

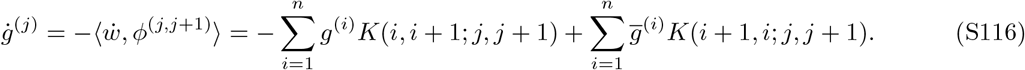

For the first sum, if *i* = *j*, the similarity will be *κ*_*s*_ and otherwise *κ*_*d*_. For the second sum, if *i* ∈ {*j* + 1, *j* − 1}, the similarity is *κ*_*o*_ and otherwise *κ*_*d*_. Thus, we can define

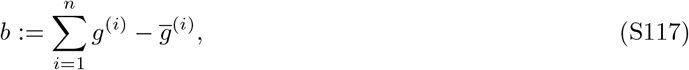

and express

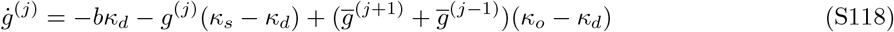

Analogously,

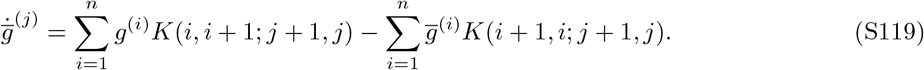

Here, the first sum has similarity *κ*_*o*_ if *i* ∈ {*j* + 1, *j* − 1} and *κ*_*d*_ otherwise. The second sum has similarity *κ*_*s*_ if *i* = *j* and *κ*_*d*_ otherwise. Thus,

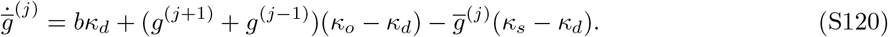

Initially 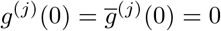. Because the differential equations are symmetric, this equality keeps holding: 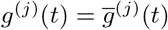 and *b* ≡ 0. Thus, we can write

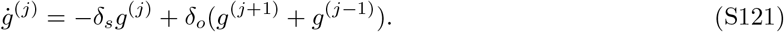

This equation, in turn is symmetric across all *j*. Because *g*^(*j*)^(0) = − 𝓁^′^(0) is equal for all *j*, this means that *g*^(*j*)^(*t*) remain equal. We thus express them as a single scalar *g*(*t*):

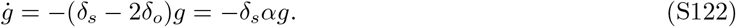

The margin is given by *m* = 1 − *g*. Its speed of change therefore depends on both the overall magnitude of the representation and the conjunctivity factor *α*. In particular, when *δ*_*s*_ = 1,

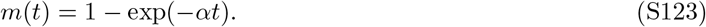

In particular, setting *m*(*t*) = 0.5 reveals the margin’s halftime:

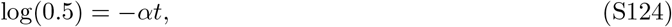

and therefore

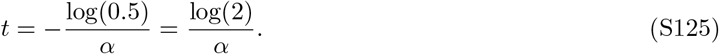

**Figure S1:**
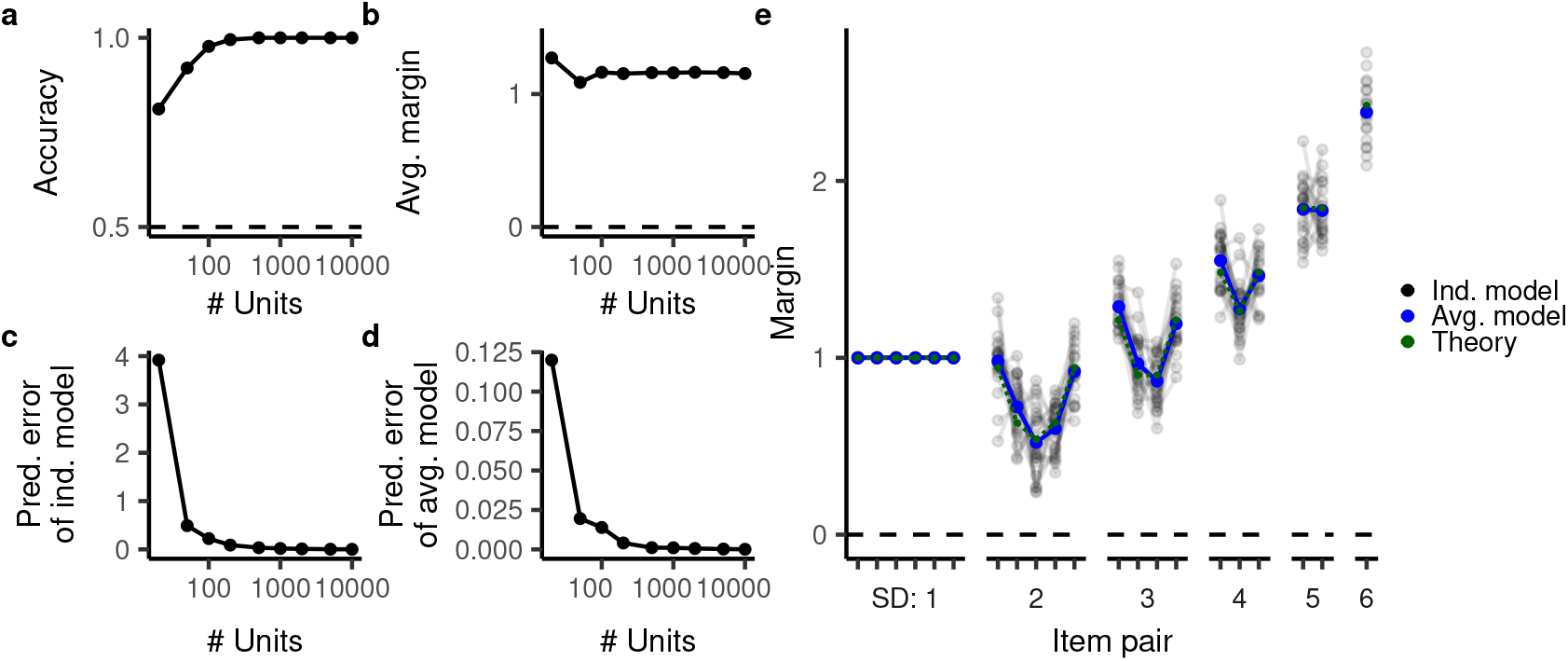
Behavior of models with fixed, randomly sampled ReLU representations with different numbers of units. **a-b**, Accuracy (a) and average margin (b). **c**, Error of the theoretical prediction compared to the individual model prediction. **d**, Error of the theoretical prediction compared to the average model prediction. **e** Margins for individual ReLU networks with 1000 units (black translucent lines), their average (blue line), and the theoretical prediction (green line).

## S2 Experimental details

All experiments and figures can be reproduced from the public Github repository: https://github.com/sflippl/relational-generalization-in-ti. This repository is also attached as Software S1.

### S2.1 Data analysis and visualization

We analysed the data in R [181] and provide an R package containing all datasets listed below. In addition, we provide infrastructure to load these datasets into Python. The visualizations were created using `ggplot2` [182] and `patchwork` [183].

### S2.2 TI behavior in models with non-exchangeable representations

To explore whether our theory can capture the behavior of a non-exchangeable representation, we explored the behavior of models with fixed, randomly sampled ReLU features. For a sufficiently small number of units, this stochasticity results in a drop in accuracy (Fig. S1a), though the margin remains the same (Fig. S1b). Further, because the representations violate exchangeability, the individual models’ behavior is not fully captured by the theoretical account. For 1000 units, for example, the models exhibited considerable spread around the theoretical prediction, though they were still well approximated (Fig. S1c,e). Notably, our account captured the average behavior across many models much better, even for a small number of units (Fig. S1d,e).

### S2.3 Deep neural network training

We trained deep neural networks with standard He initialization [180], transitioning between the lazy and rich regime by scaling the output of the networks. An output scaling of *σ* is equivalent to scaling the initialization by *σ*^1*/*(*l*+1)^, where *l* is the depth of the network. (Indeed, the main text describes the transition between the lazy and rich regime in terms of scaling the initialization.) In the lazy regime, the network adheres to a linear readout model with an additional random intercept given by its initial values 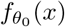 (see Appendix S1.5.1). As this complicates analysis, but does not qualitatively change behavior, we subtract this initial intercept. The complete network was therefore given by

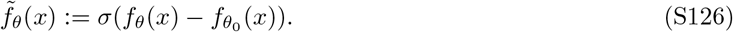

**Figure S2:**
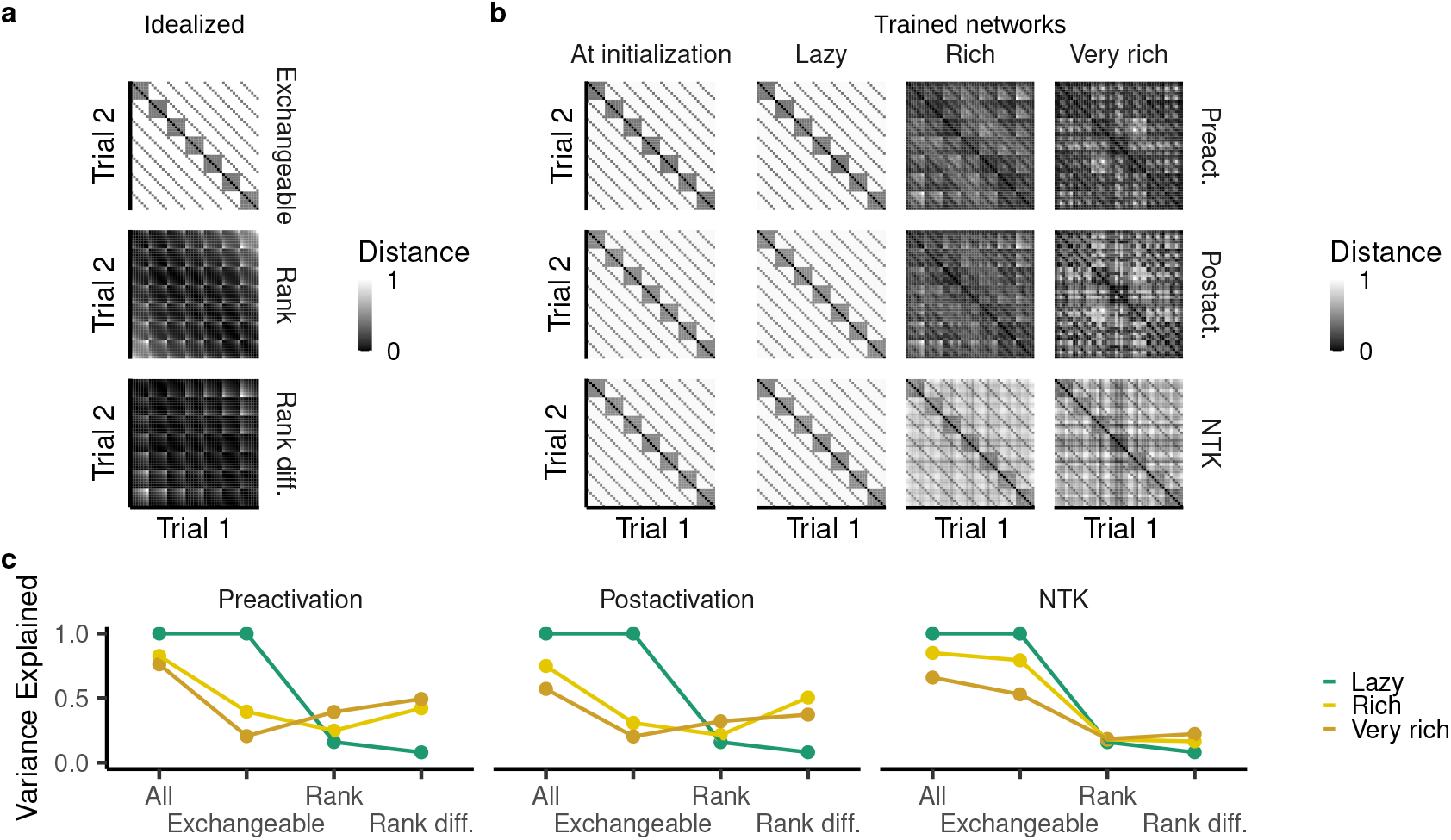
Representational dissimilarity of the hidden-layer representation, computed as the squared Euclidean distance. **a**, Trial-by-trial representational dissimilarity matrix (RDM) of an idealized hidden layer with either an exchangeable representation, a representation of both items’ rank or a representation of the rank difference. The trials are arranged in lexical ordering, i.e. AA, AB, …, AG, BA, …, BG, …, GG. **b**, RDM of the networks’ preactivations, postactivations, and neural tangent features at initialization and after lazy or rich training. **c**, Variance explained by a model which predicts the RDM in terms of a exchangeable representation, a representation of rank, a representation of rank difference, or a mixture of all three (“All”).

We implemented network training in Pytorch [184], using stochastic gradient descent without momentum. The learning rate was chosen automatically by the following process. We started at an extremely high learning rate of 10^37^. This is because we considered extremely big ranges of scalings, which accordingly tampered down the network output by factors of up to 10^−32^ and had to compensate the learning rate accordingly.) If network training at some point yielded a loss of over 1, we divided the learning rate by 10 and restarted training. This is not an efficient way to determine the optimal learning rate, but suffices for our purposes. Further, we trained the networks until they 1) had a loss under 10^−5^ and 2) did not improve their loss in the last 50 epochs.

### S2.4 Analysis of the hidden-layer representation

Fig. 6 presented the representational dissimilarity matrix (RDM) for a subset of item pairs. Fig. S2 depicts the full trial-by-trial RDM. Candidates for a useful hidden-layer representation are an exchangeable representation (Fig. 6a, top row) or a rank representation; such a rank representation may represent both items’ ranks individually (Fig. 6a, middle row) or it may represent the difference between the items’ ranks (Fig. 6a, bottom row). We compare the hypothetical representations to those empirically found in the network (Fig. S2b). While Fig. 6 presented those of the postactivation, *ϕ*(*W*^(0)^*x*), we here additionally analyse the preactivation, *W*^(0)^*x* and the neural tangent kernel (NTK; Appendix S1.5). All of them have an exchangeable representation in the lazy regime. In the rich regime, the representation exhibits some similarity to an exchangeable representation (in particular for the NTK) and perhaps a rank representation. There is also some structure, however, that appear not to be captured by any of the hypothetical RDMs.

For quantitative comparison, we try to predict the RDM by fitting an exchangeable representation, a representation of each items’ rank, a representation of rank difference, and a mixture of all three representations. The free parameters of the exchangeable representation are the three distinct similarities. The free parameters of the rank representation are distinct ranks for each item, where we allow for different ranks depending on whether an item was presented first or second. We fit these parameters using gradient descent and measure the proportion of variance explained (Fig. S2c). This confirms our previous interpretation: the lazy regime is perfectly captured by an exchangeable representation, whereas the rich regime is partly captured by an exchangeable representation. In addition, the rich regime is partly captured by a representation of rank difference. However, even a mixture of all three hypothetical representations cannot fully explain the representational structure.

**Figure S3:**
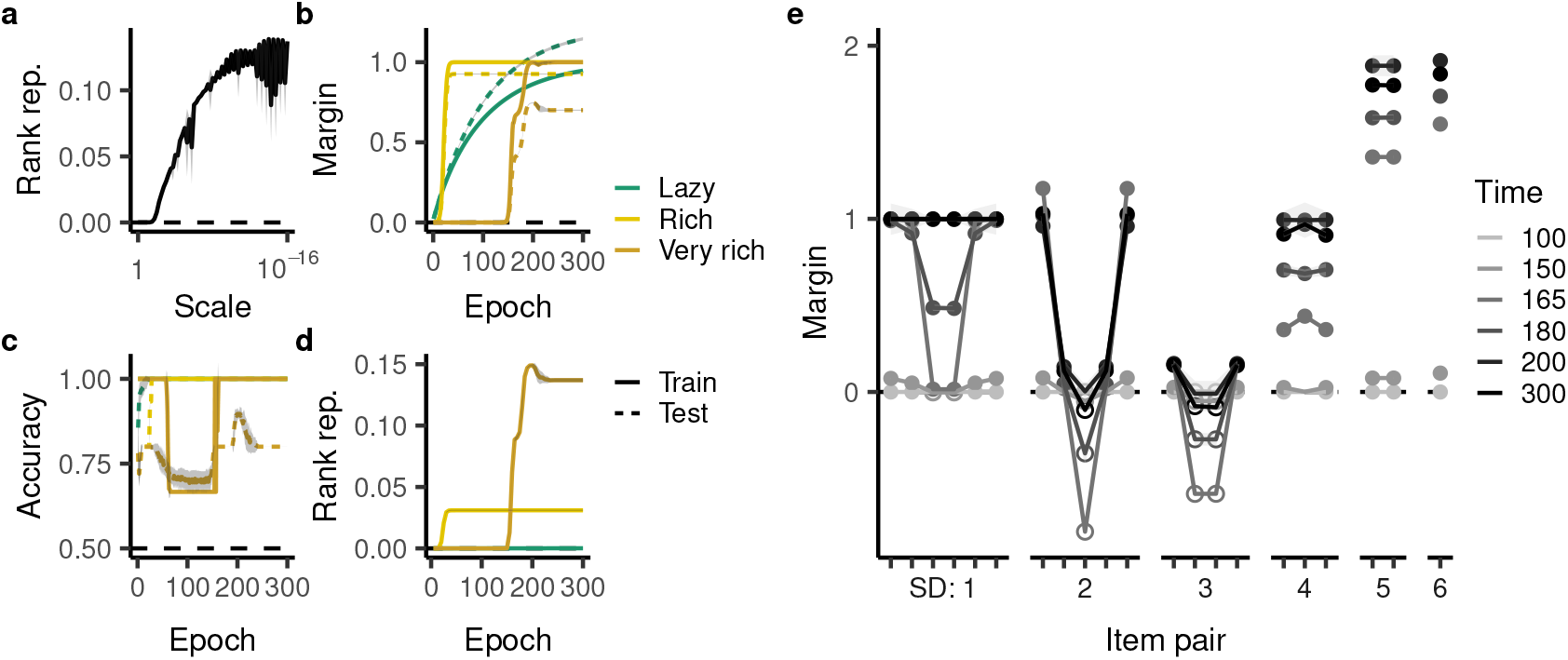
TI behavior of deep networks as a function of initialization scale and training duration. **a**, Rank representability at the end of training as a function of initialization scale. **b**-**d**, Margin, accuracy, and rank representability over the first 300 epochs. Margins (i.e. *ŷ y*, where *ŷ* is the model prediction and *y* ∈ {− 1, 1 } is the trial label) are averaged over all trials within the training and test set. **e**, Margin at the different item pairs as a function of training duration (very rich regime).

### S2.5 Detailed analysis of deep neural network behavior on transitive inference

This section presents a more detailed analysis of the deep network behavior. The results below demonstrate, in particular, that changing the activation function or considering deeper networks does not straightforwardly improve TI behavior. Indeed, these extended results further demonstrate the fundamental strangeness of the deep networks’ behavior on TI.

#### S2.5.1 Rank representability

The “rank representability” measures how well TI behavior can be captured by a rank representation, a value of 0 indicating that it can be perfectly captured. Specifically, a linear model is fit to approximate

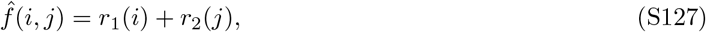

where *r*_1_ and *r*_2_ are individually estimated for each *i* and *j*. The rank representability is then given by the mean squared error of this approximation. As expected, in the lazy regime, the rank representability is zero. As the scaling decreases (and training enters the rich regime), the error increases, indicating that network behavior cannot be described by a rank representation anymore (Fig. S3a). This confirms the picture observed in Fig. 6e.

#### S2.5.2 Time trajectory

Fig. S3b-e depict the behavior of the neural network with one hidden layer over training. Notably, learning in the rich regime occurs much more suddenly (Fig. S3b) than in the lazy regime. Further, the TI behavior over training exhibits several curious intermediate stages (Fig. S3e). Specifically, the terminal items are learned essentially completely before the intermediate item pairs even improve slightly. In fact, the margin of the item pairs CE, BE, and CF decreases considerable before increasing again.

**Figure S4:**
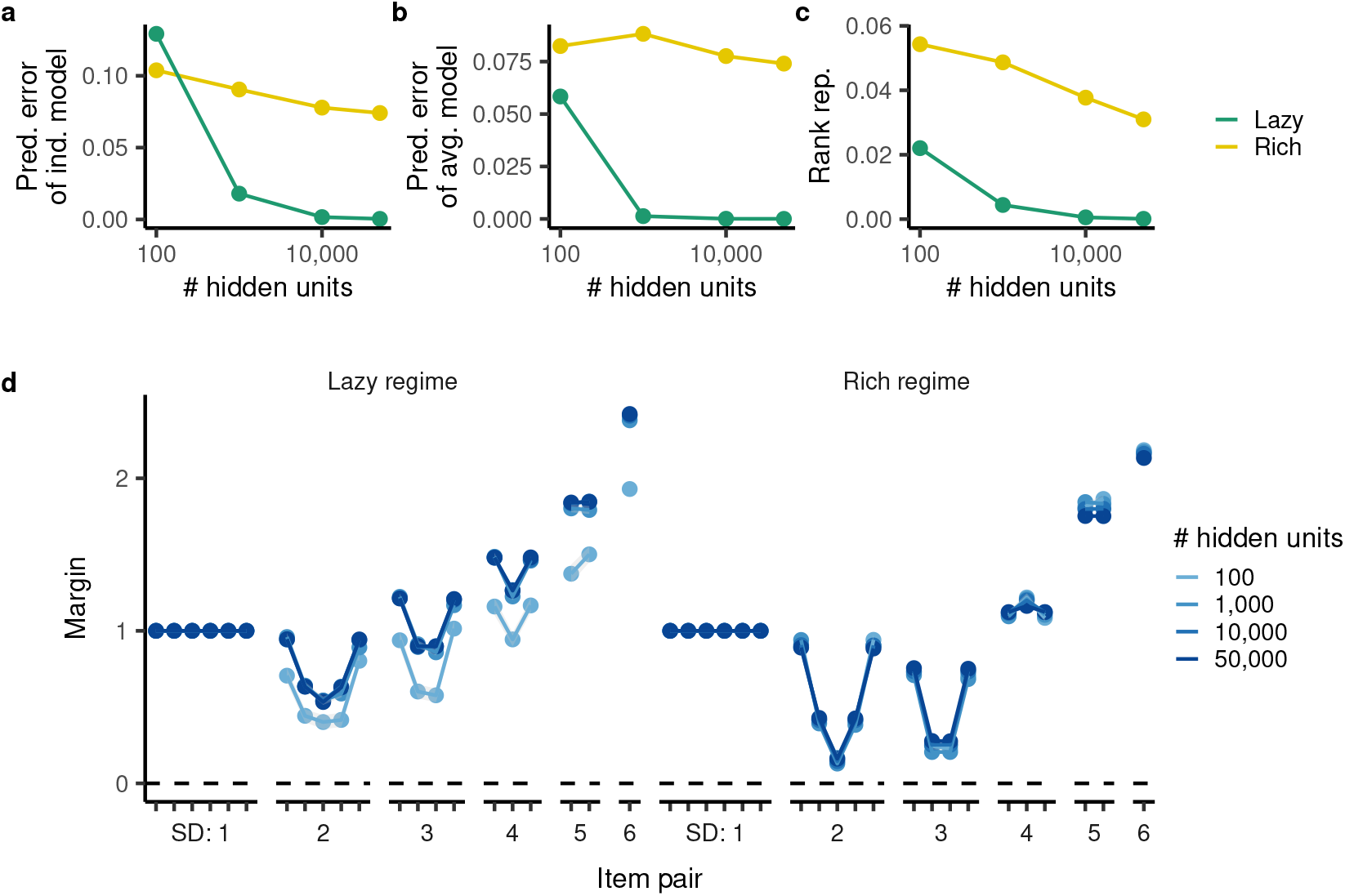
TI behavior for different numbers of hidden units and standard initialization (“lazy”) as well as smaller initialization (scale 10^−3^, “rich”). **a**, Error of the theoretical prediction compared to the individual model prediction. **b**, Error of the theoretical prediction compared to the average model prediction. **c**, Rank representability as a function of the number of hidden units. **d**, Predictions of the networks in different regimes and for different numbers of hidden units.

#### S2.5.3 Effect of the number of hidden units

At a fixed initialization, neural networks approach the lazy regime as they become wide [85, 86]. Neural networks with few hidden units therefore tend to be in the rich regime at standard initialization. Further, the random initialization leads to deviations from the NTK predictions (which are generated from the expected neural tangent kernel) even in the lazy regime; these deviations are larger for fewer hidden units. We therefore decided to explore how the number of hidden units affects the behavior of the network at different initialization scales. We find that at 100 and 1000 hidden units, the network behavior deviates substantially from our theoretical prediction even at standard initialization (Fig. S4a). Part of this is due to the random noise of individual initializations. As a result, the prediction of twenty models averaged together are much more accurate and, in fact, close to zero for 1,000 hidden units (Fig. S4b). At 100 units, in contrast, we do observe a systematic bias (Fig. S4d). Further, at 100 units, model behavior cannot be entirely described by a rank representation, whereas even individual models with 1,000 units do implement a rank representation (Fig. S4c). Taken together, this suggests that at 100 hidden units, the neural network is in the rich regime even at standard initialization, but the effect of the rich regime can be exacerbated by further decreasing the scale of initialization.

#### S2.5.4 Effect of the nonlinearity

Here we explore the effect of different nonlinearities: both piecewise linear functions and a tanh function as an instance of a saturating nonlinearity. Generally, as the difference in slopes *ρ* of the piecewise linear function increases (see Appendix S1.3 for a definition of *ρ*), test margin and test accuracy decrease (Fig. S5a,b) and their rank representation error increases (Fig. S5c). In fact, for *ρ* ≥ 1.2, the network performs *below chance-level*. While TI behavior is better at *ρ* = 0.5, it is still non-naturalistic, as an inverse terminal item effect is consistently observed (Fig. S5g). For the tanh nonlinearity, it is not as clear how the rich regime can be approached, as a small output scaling can only be overcome by making the readout large due to the saturating nonlinearity. Here we explore two different scalings: 10^−6^ and 10^−22^. We find that the tanh nonlinearity has better performance, though it is still characterized by an extremely sharp terminal item effect at a symbolic distance of two (Fig. S5g).

**Figure S5:**
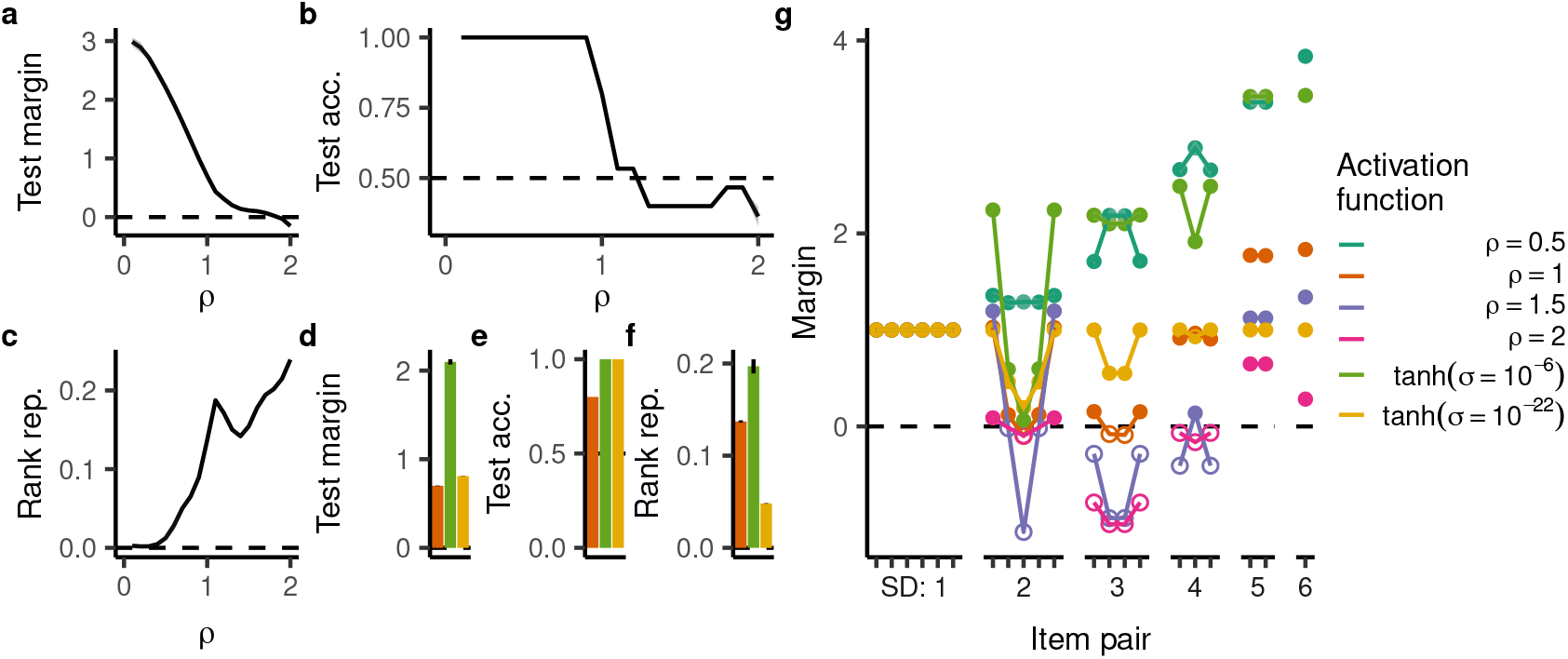
TI behavior for different activation functions. **a**-**c**, Margin (a), accuracy (b), and rank representability (c) as a function of *ρ* (see (S11)). **d**-**f**, Margin (d), accuracy (e) and rank representability (f) for the tanh activation function at two different output scalings *σ* (as well as, for comparison, the ReLU activation function (*ρ* = 1)). **g**, Margin of the different item pairs for selected activation functions.

#### S2.5.5 Effect of the number of items

We next determined the behavior of ReLU networks with one hidden layer in the very rich regime for different numbers of items. Generally, both test margin and accuracy decreased as the number of items increased (Fig. S6a,b). The rank representability exhibited a more non-monotonic pattern. In particular, none of the networks except for those trained on 5-item TI exhibited a symbolic distance effect (Fig. S6d). Instead their margin oscillated at intermediate symbolic distances before eventually increasing at large symbolic distances.

#### S2.5.6 Networks trained on crossentropy

Neural networks are often trained on crossentropy rather than mean squared error. We therefore study the inductive bias of neural networks trained on TI with crossentropy. Notably, the learning trajectory of neural networks trained on crossentropy can generally not be approximated by their neural tangent kernel as the weights must diverge to minimize the loss function. This is because crossentropy is minimized as the margins diverge towards infinity (see Remark S1.7). Nevertheless, we consider networks with standard (scaling 1) or small (scaling 10^−3^) initialization and find that they behave differently, presumably because we train for a finite number of epochs. Further, we consider both learning by backpropagation and only adjusting the linear readout.

While the margins grow over the duration of learning (Fig. S7a), we find that the normalized margins of the linear readout model are well approximated by our theory. This confirms our theoretical results (Remark S1.7). More surprisingly, the neural network trained by backpropagation with standard initialization is also well approximated by our theory (Fig. S7b). Only those networks trained with backpropagation and initialized with small weights exhibit behavior similar to that of neural network trained on MSE in the rich regime (Fig. S7a-c). There are two potential reasons for this: it is possible that after longer amounts of training, the neural network with standard initialization may exhibit similar behavior to that with smaller initialization [100]. It is also possible that different initializations genuinely affect the inductive bias of neural networks trained with crossentropy. This is an interesting possibility especially in light of the fact that known theoretical results are largely silent on such an effect [99, 100].

**Figure S6:**
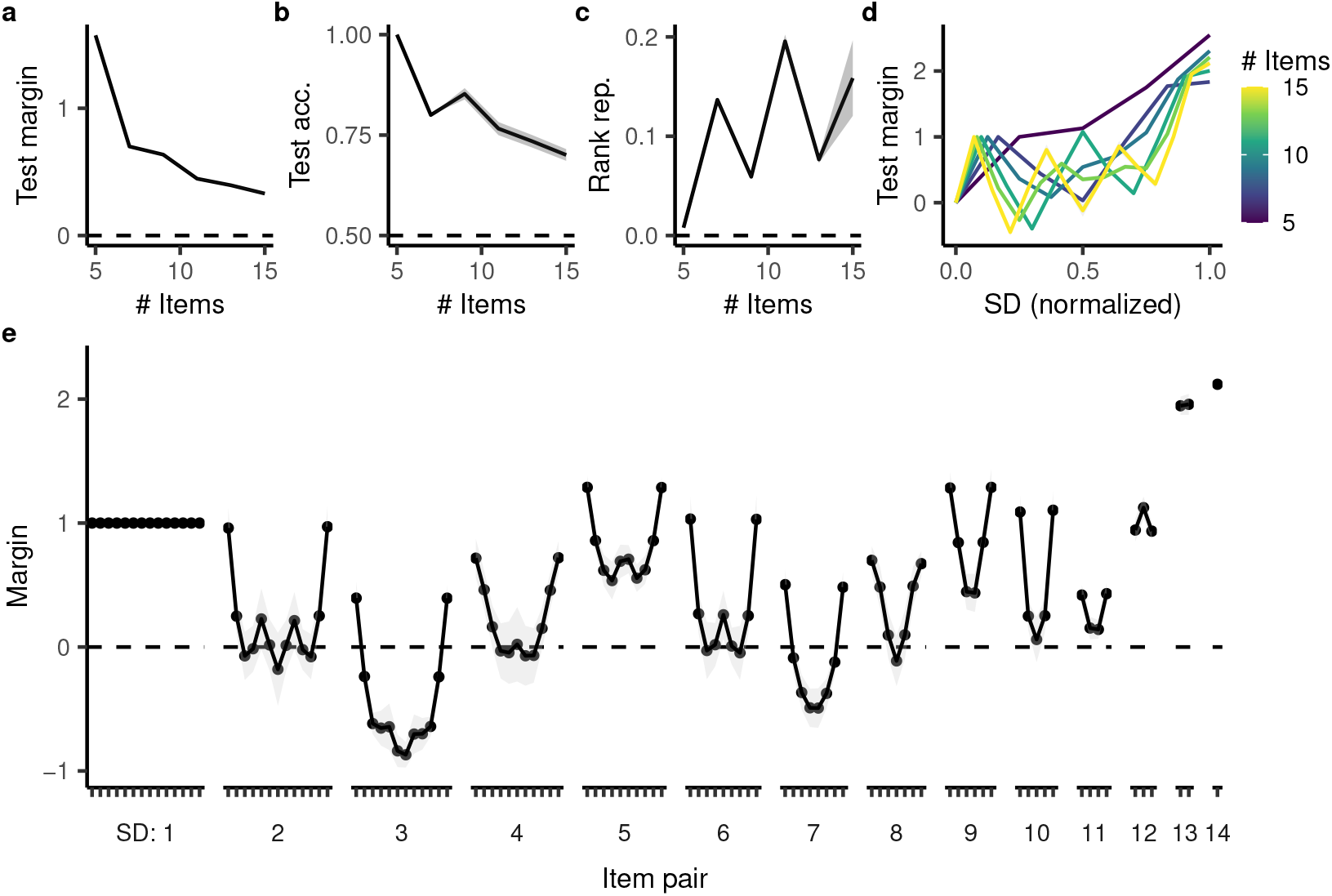
TI behavior as a function of the number of items. **a**-**c**, Margin, accuracy, and rank representability as a function of the number of items. **d**, The average test margin for a given symbolic distance (normalized by *n* − 1). **e**, Margin at the different item pairs for 15-item TI.

#### S2.5.7 Deeper networks

While neural networks with one hidden layer already exhibit representation learning, the superior generalization of deep neural networks is often tied to their depth. We therefore explore how increasing depth changes the networks’ inductive bias on TI. We use an initialization scale of 10^−3^ and train networks with one to three hidden layers. (Deeper networks do not converge to a minimal loss within six hours and we leave their exploration to future work.) We find little difference between the behavior of networks with different depth (Fig. S7d), suggesting that simply increasing the depth of the networks is not sufficient to improve their TI performance.

### S2.6 Symmetric weights in the input layer

In related work, Nelli *et al*. [30] also compare the TI behavior of deep neural networks in the lazy and rich regime, finding that neural networks trained in the rich regime have an emergent rank representation in their hidden layer. Notably, their networks implement a weight symmetry between the two input items. Specifically, given two items *x*_1_, *x*_2_ ∈ ℝ^*d*^, their network computes the output as

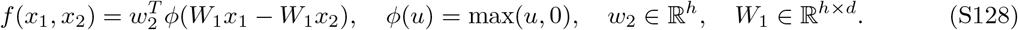

At first glance, this network is not additive – after all, *ϕ* mixes the two items nonlinearly. However, only the directional bias can be nonlinear. The average choice of the network between either order of presentation,

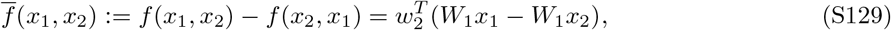

is additive, as *ϕ*(*u*) − *ϕ*(− *u*) = *u*.

Indeed, weight symmetry strongly improves test performance in the rich and lazy regime (Fig. S8a). In line with the parameters used in [30], the rich regime corresponds to an initialization scaling of 0.025 in the hidden layer and a scaling of 1 in the readout layer, whereas the lazy regime corresponds to a scaling of 1 in both layers. Note that training performance is suboptimal because the network is only trained for 40 epochs. The RDM of the postactivation (Fig. S8b) corresponds to a rank difference representation (Fig. S8c), whereas such a representation leaves a substantial proportion of variance unexplained in the lazy regime as well as without weight symmetry. Notably, neither the preactivation nor the NTK can be explained as a rank difference representation.

**Figure S7:**
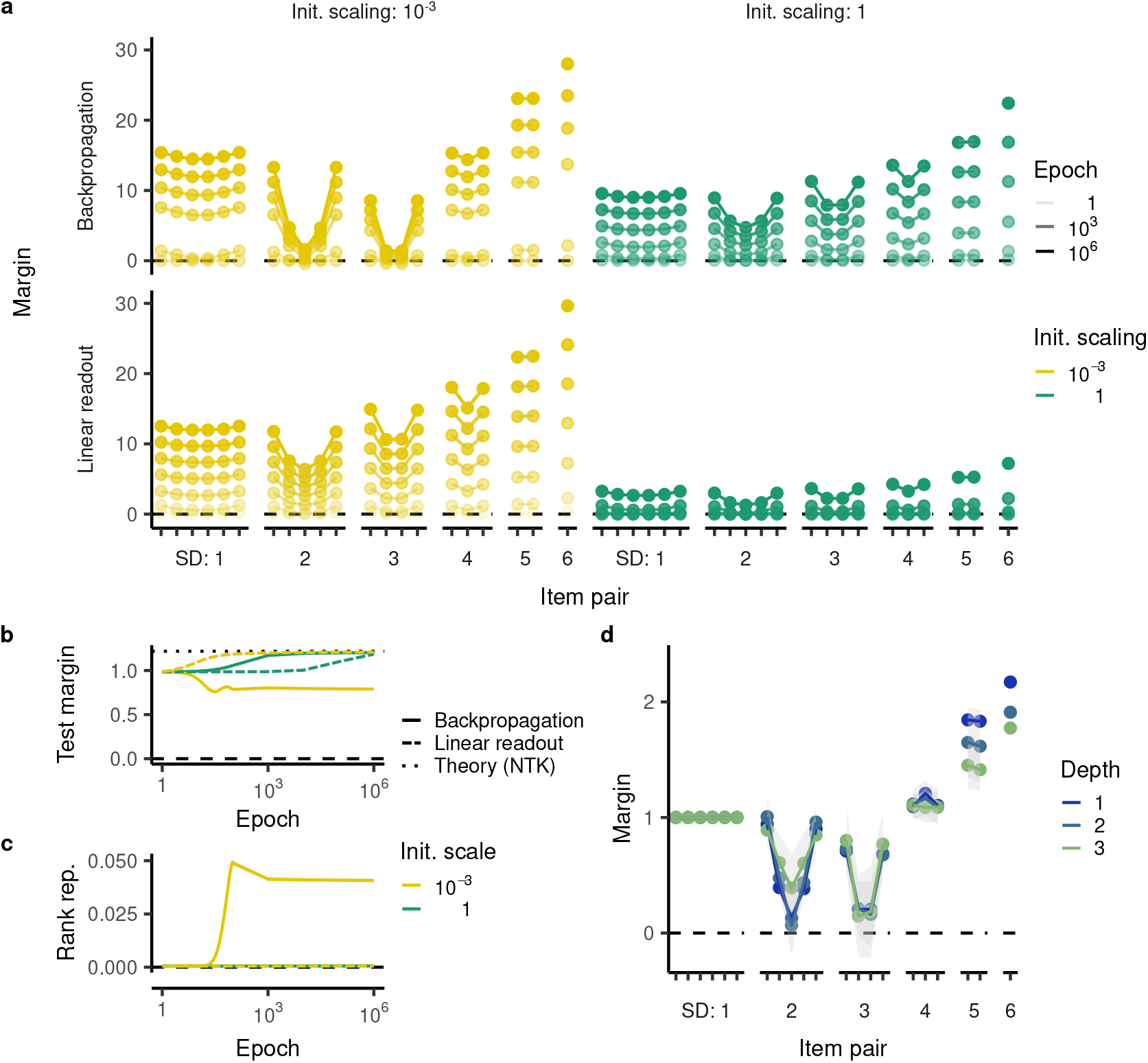
**a**, TI behavior of networks trained on crossentropy. The top row depicts a network that has been trained by backpropagation; the bottom row a network whose linear readout has been trained. Further, the left column is initialized with smaller weights than the right column. **b**, Test margin of the networks over training. **c**, Rank representability of the networks over training. **d**, TI behavior of networks with different depths at an initialization scale of 10^−3^.

In addition to weight symmetry, Nelli *et al*. implement training with some differences to our implementation: they run stochastic gradient descent with a batch size of one, use networks with only twenty units, run training for 40 epochs rather than until convergence, and initialize the hidden-layer weights with a smaller variance than the output weights (specifically multiplying them by a factor of 0.025). To investigate whether these differences are responsible for the emergence of a rank difference representation, we changed each of these hyperparameters independently. We found that weight symmetry is necessary and sufficient for the emergence of a rank difference representation. Note that in the wide network, the rank difference representation is weaker because the network is leaving the rich regime due to its increasing width. To re-enter the rich regime, we would have to initialize the weights by a smaller factor.

**Figure S8:**
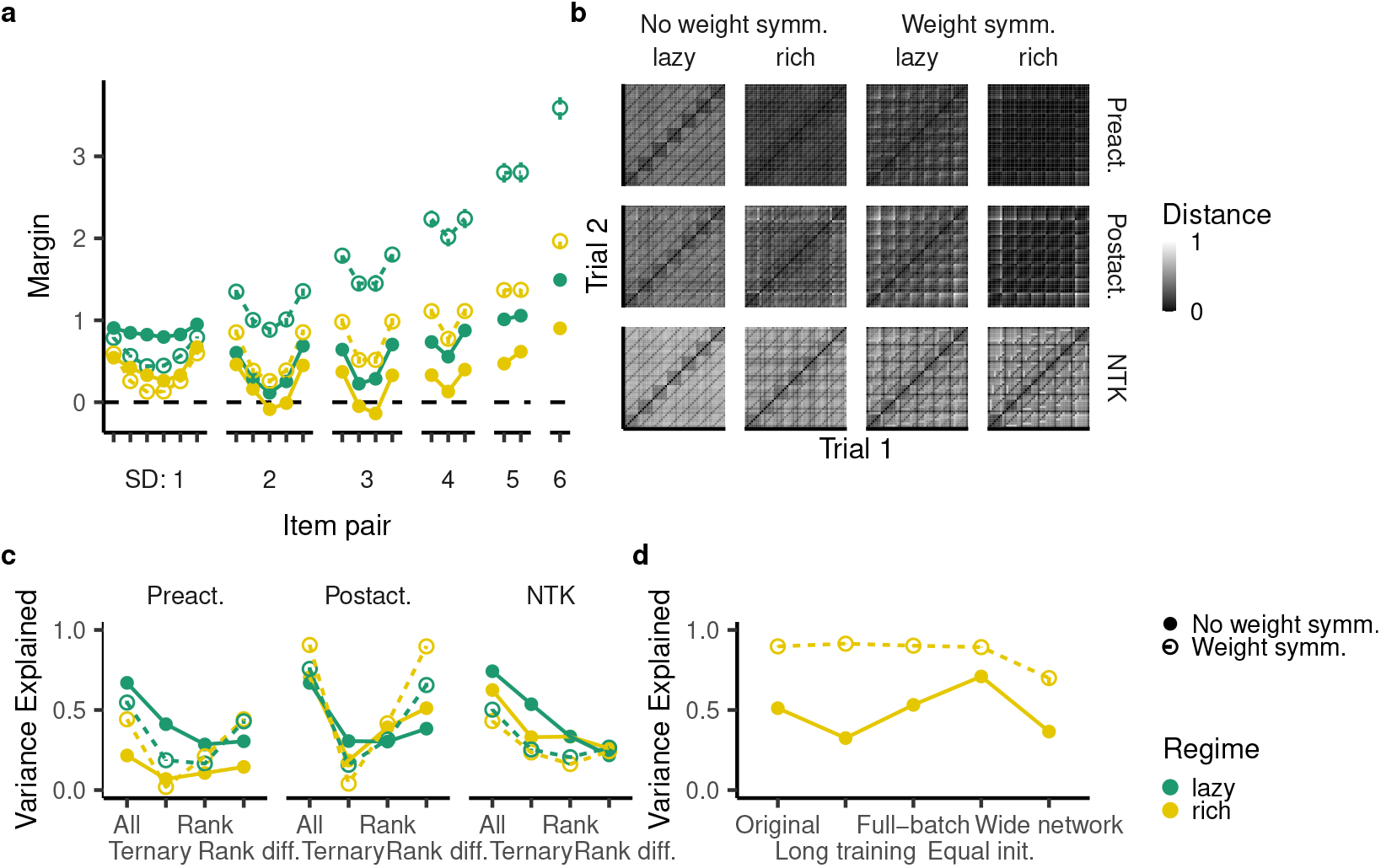
Weight symmetry restores generalization in the rich regime [30]. Specifically, weight symmetry is implemented as *W*^(1)^ = − *W*^(2)^, where *W*^(1)^ are the hidden weights corresponding to the first item and *W*^(2)^ are the hidden weights corresponding to the second item. **a**, Generalization behavior in the lazy and rich regime with and without weight symmetry. **b**, Trial-by-trial RDM of the preactivation, postactivation and NTK in the lazy and rich regime with and without weight symmetry. **c**, Variance of those RDMs explained by a exchangeable representation, a rank representation, a representation of rank difference, and a mixture of all representations. **d**, Variance explained of the RDMs that introduce one of the changes introduced by our training protocol. Long training indicates training until convergence and equal init. indicates that both the hidden layer and the readout layer were initialized with a scaling of 0.025. Finally, the wide network indicates a network with 1000 units.

## S3 Analysis of the rich-regime behavior

For hidden weights *W* ∈ ℝ^*H×d*^, 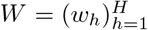 and readout weights *v* ∈ ℝ^*H*^, the ReLU network is expressed by

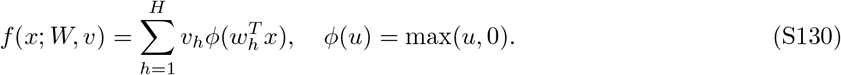

Because the ReLU nonlinearity *ϕ* is positively homogeneous (*ϕ*(*su*) = *sϕ*(*u*) for all *s* > 0), we can rewrite this network using the hidden weight directions *ŵ*_*h*_ = *w*_*h*_*/* ∥ *w*_*h*_ ∥_2_ ∈ ℝ^*d*^, the magnitude *m*_*h*_ = ∥ *w*_*h*_ ∥_2_ | *v*_*h*_ | ∈ ℝ, and the sign *σ*_*h*_ = sign(*v*_*h*_) ∈ {−1, 1}:

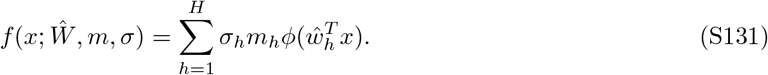

Prior work has found that minimizing the 𝓁_2_-norm of all weights in the network is equivalent to minimizing 2∥ *m* ∥_1_ (where the weight directions can be freely chosen) [100, 102, 103]. Because the 𝓁_1_-norm induces sparsity, this norm incentivizes a sparse set of hidden units, in addition to keeping the magnitude *m*_*h*_ of those hidden units low.

Below we describe the periodic coding schema for TI with arbitrary numbers of items *n* and compute the network’s weight norm. We then compute the weight norm of a rank-network, which instead implements a ranking system, finding that it is much larger. We then compare these hand-constructed networks to neural network simulations, finding that they capture important properties of empirically trained networks and describing specific differences.

### S3.1 Cooperative coding schema

The network we describe here uses four units (H_1+_, H_2+_, H_1−_, and H_2−_) to fit the training trials on TI. H_1+_ and H_2+_ have positive readout weights and H_1−_ and H_2−_ have negative readout weights:

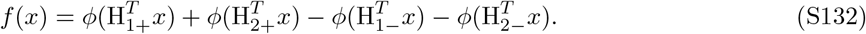

These units partition the dataset 𝒟 (see (S36)) into four disjoint subsets:

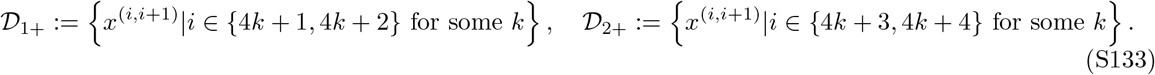

𝒟_1−_ and 𝒟_2−_ are defined analogously for *x*^(*i*+1,*i*)^. For example, for *n* = 7,

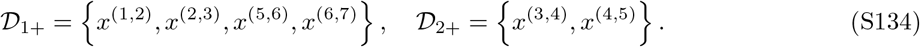

Each unit H_Δ_, Δ ∈ {1+, 2+, 1−, 2−} then outputs one on its own data points and zero on all others:

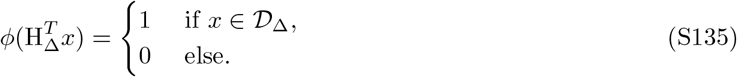

Below we describe how its weights allow it to do so. H_Δ_ ∈ ℝ^2*n*^ has one weight for each item and position. We write H_Δ_(*i, s*) for item *i* ∈ {1, …, *n*} in position *s* ∈ {1, 2}. Now consider a data point *x*^(*i,i*+1)^ ∈ 𝒟_Δ_ (i.e. Δ ∈ {1+, 2+}). The unit then distributes its weights equally across those two items:

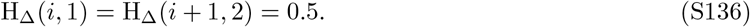

However, this also means that it would response positively to the data points *x*^(*i,i*−1)^, *x*^(*i*+2,*i*+1)^ ∉ 𝒟_Δ_. To avoid this, the unit requires a negative weight in response to the other item for those trials:

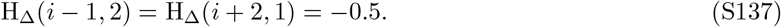

This same pattern can be repeated for two consecutive trials, e.g. *x*^(*i,i*+1)^, *x*^(*i*+1,*i*+2)^. However, encoding *x*^(*i*+2,*i*+3)^ into the same unit would conflict with (S137). For this reason, the specialization of units switches every two trials.

Note that for Δ ∈ {1−, 2−}, the exact roles of the first and second entry are reversed, i.e.

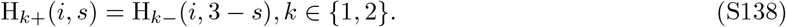

This logic allows us to define a formula that constructs this network for arbitrary numbers of trials. We show the example coding scheme for *n* = 7 in Fig. S9a,b. The above intuition also helps us get a rough estimate of the magnitude of these weights. The total number of training trials is 2(*n* − 1). Each unit H_Δ_ is responsible for roughly a quarter of those. For each trial, these units require four weights at a magnitude of 0.5. This means that each unit’s 𝓁_2_-norm (i.e. its magnitude) is roughly:

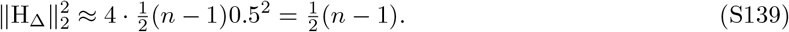

Thus, the networks overall weight norm is approximately

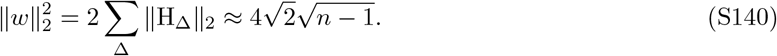

To determine the exact weight norm, we need to treat different numbers of trials slightly differently. This gives rise to a slightly more complicated formula that is still on the order of 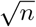 (Fig. S9d, details in Appendix S4.6).

**Figure S9:**
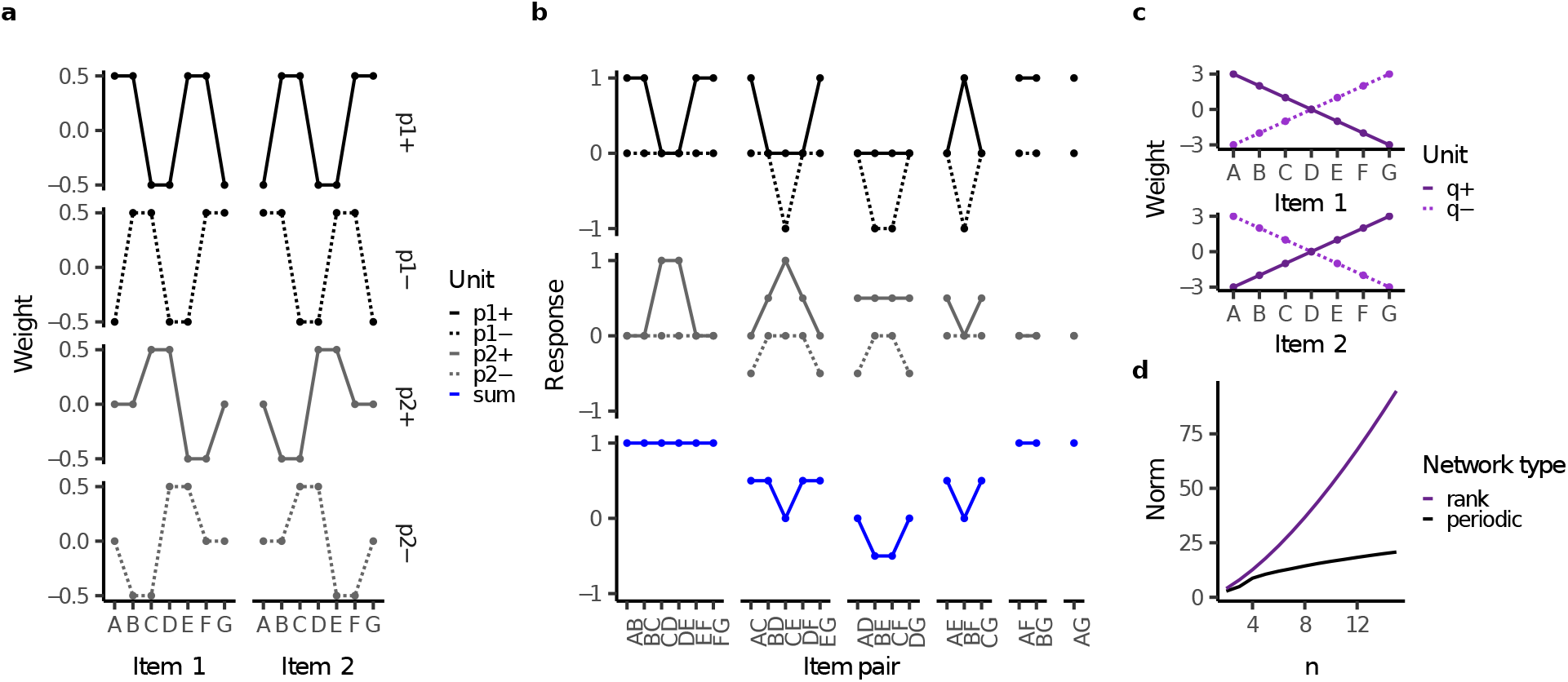
Hand-constructed networks. **a**, Weights of the network implementing the cooperative coding schema for *n* = 7. **b**, Responses to all trials with positive outputs. First row depicts the units H_1+_, H_1−_, second row depicts H_2+_ and H_2−_ and the third row depicts their sum. **c**, Weights of the rank network for *n* = 7. **d**, Norm of the two networks as a function of the number of items *n*.

### S3.2 Rank network

We contrast this cooperative coding schema with a simple ranking system in the hidden layer (Fig. S9c). Specifically, such a network would consist of two units *q*_+_, *q*_−_ and compute its response as

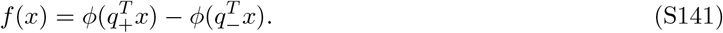

Furthermore, *q*_+_ = −*q*_−_ and therefore

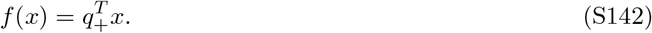

*q*_+_ then simply encodes the rank of each item, centered around zero:

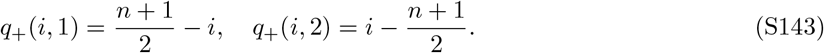

We now compute the magnitude of these units:

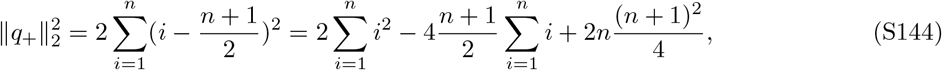

where we used Bernoulli’s formula. Note that [see e.g. 185]

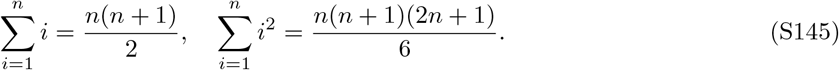

Thus,

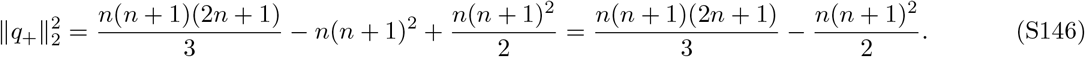

Simplifying this implies

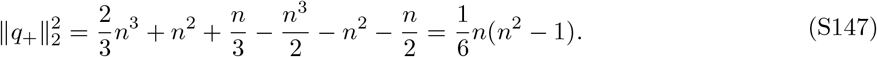

Thus, the network has a total weight norm of

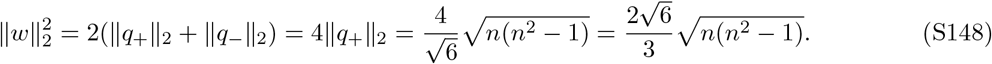

This weight norm is considerably bigger than that of the network implementing a cooperative coding schema. In particular, the norm is *O*(*n*^1.5^), while the latter network has a norm of 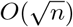 (Fig. S9d).

### S3.3 Comparison with empirical networks

Here we provide a detailed comparison between the hand-constructed networks and the empirically computed deep networks.

#### S3.3.1 Norm comparison

First, we compute the empirical networks’ norm. The hand-constructed networks have a lower norm than their empirical counterpart, but the values are generally close together (Fig. S10a). Further, the norms follow the same trends with respect to *n*. In particular, our theory would predict that 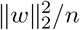 should increase for the symmetric network (which should be of order *O*(*n*^1.5^)) but decrease for the dense network (of order *O*(*n*^0.5^)). Indeed, this is the case (Fig. S10b). Note that the symmetric network’s norm appears to increase a bit faster than that of the rank network.

The suboptimalities in the empirical networks may have arisen because they found a local stationary point rather than a global minimum, because the initialization scale of 10^−16^ was still not small enough, or because training on mean squared error may not exactly minimize the 𝓁_2_-norm [101] (though we note that networks trained on crossentropy, which provably converge to a stationary point, implement the same mechanism).

#### S3.3.2 Clustering analysis

We assessed the network’s sparsity in hidden units by performing *k*-means clustering over the normalized weights 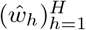, weighting each sample by its respective magnitude *m*_*h*_. We perform this clustering analysis separately for units with positive and negative readout weights, for the sake of easier interpretability. After identifying *k* clusters, we replace each hidden weight by its respective cluster centroid, giving rise to a network with *k* hidden units.

We analyzed networks trained on TI with *n* = 5, …, 15. We consistently found that these networks were well explained by six clusters in total (Fig. S10c). To compare the units extracted from networks with different initializations, we identified the unit-to-unit mapping that minimized the deviation between the different networks’ units. Fig. S10d,e depict the three units with positive and negative signs. Notably, their weights are symmetric in that each positive unit responds to item 1 in the same way that the corresponding negative unit responds to item 2 and vice versa. Finally, Fig. S10f depicts the six units extracted from the networks trained on 15 items. Again, these units are remarkably consistent.

#### S3.3.3 Differences between empirical and hand-constructed networks

We observe a few differences between the empirical and hand-constructed networks, as highlighted in Fig. 6h in the main text. We hand-construct two networks that interpolate between the solution described in the main text (HCN) and the empirical networks (for *n* = 7) (Table 1). In modification 1, the network does not silence the second unit on the trials BC/CB and EF/FE. Instead, the weights of the first unit are increased to compensate for this change. This, in turn, causes other weights to change as well to prevent this change from having ramifications. As a result, this network has a lower weight norm on unit 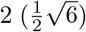 but a strongly increased weight norm on unit 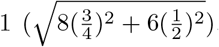. Keeping in mind that the network has corresponding negative units (and that, to compute its norm, the total 𝓁_2_-norm of its units needs to be multiplied by 2), its norm is

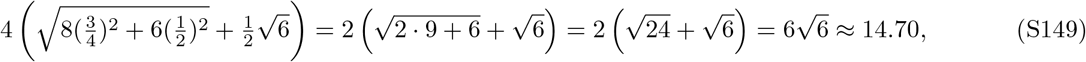

which is higher than the periodic network’s norm of 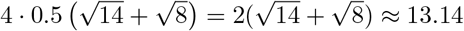.

**Table 1:**
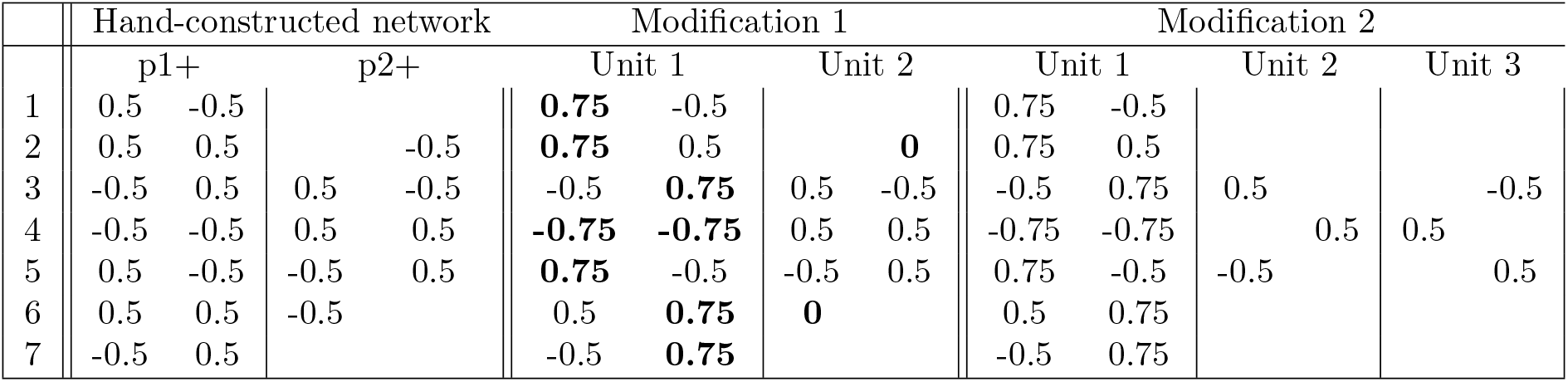
Weights associated with the hand-constructed network described in the main text on *n* = 7 as well as two modifications that account for the differences empirically observed. Bolded weights emphasize changes from the periodic network to modification 1.

**Figure S10:**
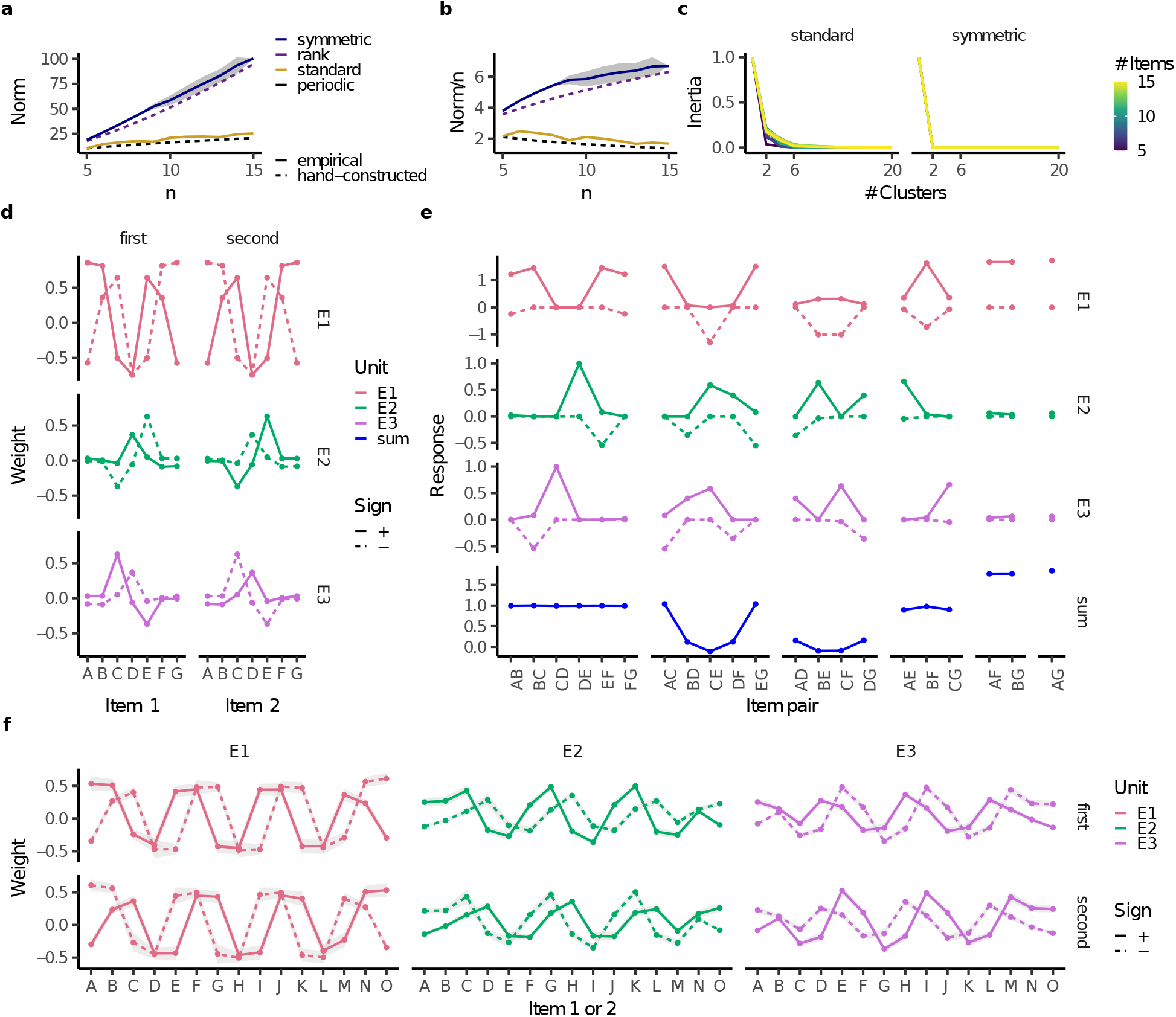
Extended analysis of empirical rich-regime networks (initialization scale 10^−16^). All figures depicted are averaged across twenty random initializations; shaded areas show ± one standard deviation. **a**,**b**, (a) Norm and (b) Norm divided by the number of items of standard and symmetric networks trained on different numbers of items (solid lines), compared to their corresponding hand-constructed network. **c** Inertia (i.e. proportion of unexplained variance) as a function the number of clusters, the network structure, and the number of items the networks were trained on. **d**, Weights of the three positive and negative centroid units. **e**, Response of the different units to all item pairs. (Negative item pairs look analogously, though with the role of positive and negative units switched. This is apparent from the symmetry in panel d.) **f**, Weights of the three positive and negative units for networks trained on 15 items. They indeed have a periodic structure approximately described by the periodic network and E2 and E3 continue to share the role of H2.

The second modification then turns unit 2 into to units with non-overlapping units. Even though the total number of weights doesn’t increase, the decrease in sparsity yields a higher weight norm. Specifically, each of those units has a norm of 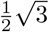 and their sum, 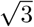 is higher than the previous unit’s norm 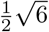. In total, this network has a norm of

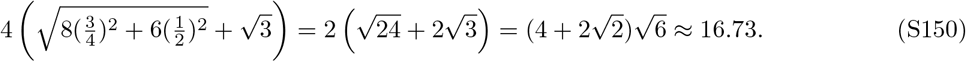

This is very close to the networks average weight norm, 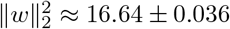.

## S4 Detailed mathematical analysis

### S4.1 Prerequisites

#### S4.1.1 Results on Tridiagonal Matrices

##### Theorem S4.1

(Hu & O’Connell [179]). *Let a, b* ∈ ℝ *such that* 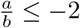 *and*

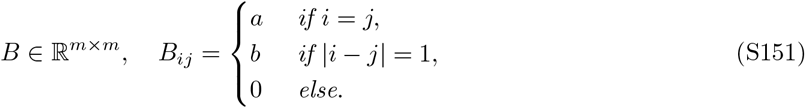

*Then*

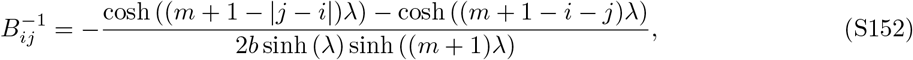

*where*

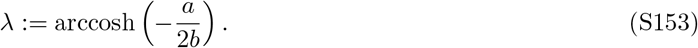

*Proof*. We define 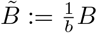. Using [179],

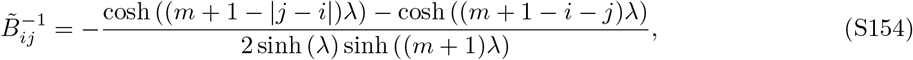

and thus

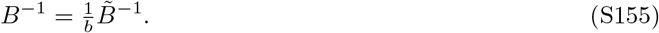

##### Theorem S4.2

(Böttcher & Grudsky [186]). *A tridiagonal matrix*

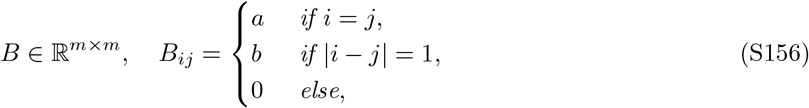

*has the eigenvalue decomposition*

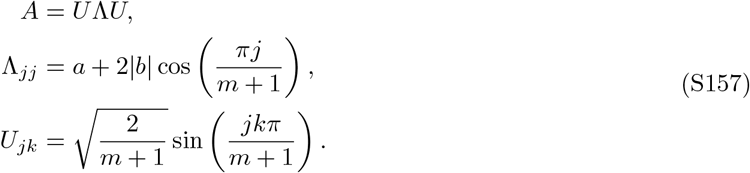

*Proof*. This is a special case of [186, Theorem 2.4]. All that remains is proving the scaling factor. We use

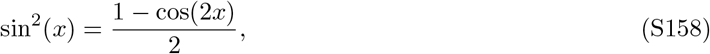

to infer

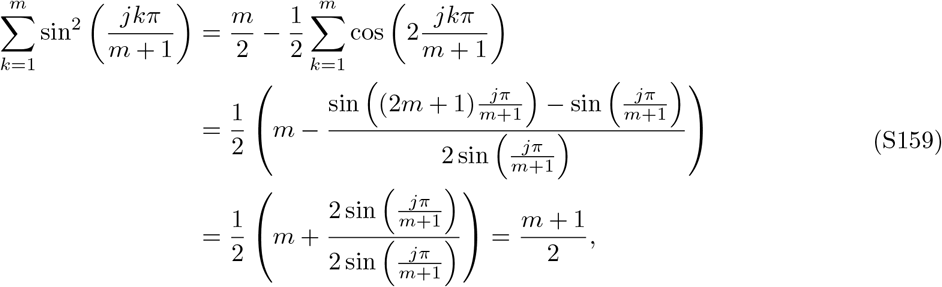

where the last equality holds because

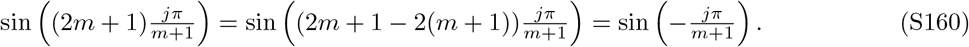

Note that we know that the eigenvectors are orthogonal because the matrix is symmetric and thus normal.

#### S4.1.2 Hyperbolic identities

##### Lemma S4.3

*For any x, and m, n* ∈ ℕ,

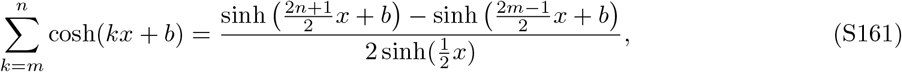

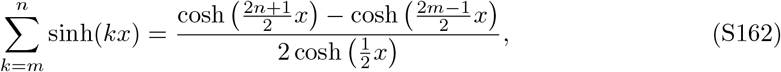

*Proof*. We prove (S161) (as in this StackExchange response) by

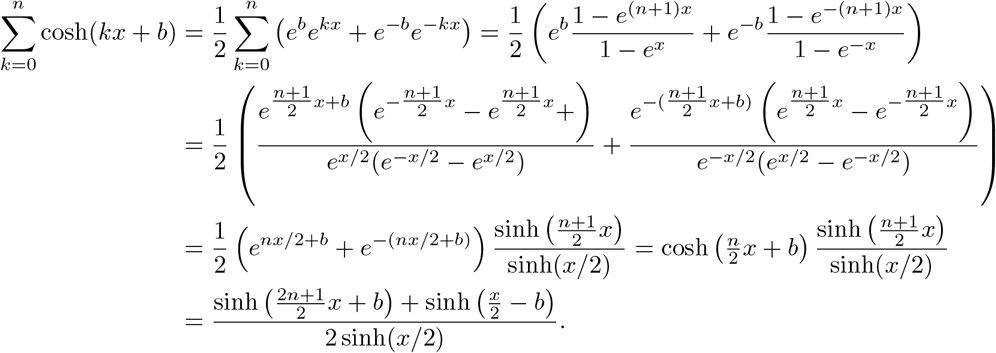

Analogously, we prove (S162) by

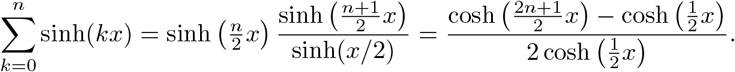

#### S4.1.3 Fixed-point theorem

In Appendix S4.5, we use a fixed-point theorem that works with a sequence of functions:

##### Theorem S4.4

(Gill [187]). *Let* (*f*_*n*_)_*n*_ *be a sequence of functions that are analytic on a simply connected region S and continuous on the closure S*^′^ *of S. Define the outer compositional sequence as*

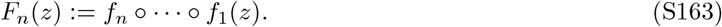

*Suppose there exists a compact set* Ω *contained in S such that* Ω ⊃ *f*_*n*_(*S*) *for all n. Furthermore, suppose the sequence of fixed points* (*α*_*n*_)_*n*_ *of* (*f*_*n*_)_*n*_ (*in* Ω) *converges to a number α. Then F*_*n*_(*z*) → *α uniformly on S*^′^.

### S4.2 Upper bounds on *α*

We here construct a set of features that encodes overlapping item pairs as less similar than distinct item pairs. To do so, we modify a onehot encoding to contain negative loadings at a trial’s overlapping trial loadings. Specifically, our features

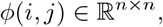

are defined as

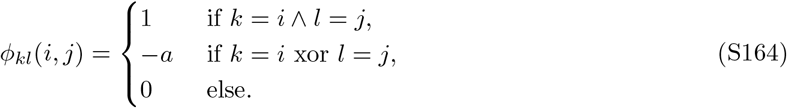

*a* is a hyperparameter that will depend on *n*.

We compute the similarities as

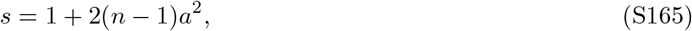

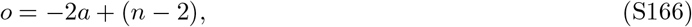

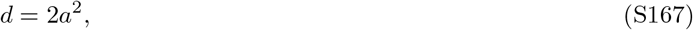

and therefore

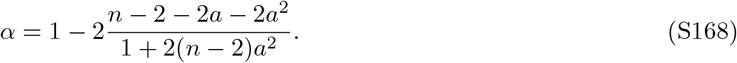

As the determinant of the polynomial in the numerator, 4 + 8(*n* − 2), is positive for all *n*, this polynomial can take on negative values for all *n*. As a result max_*a*_ *α* > 1 for all *n*, demonstrating that this construction encodes overlapping item pairs as less similar than distinct item pairs. Determining its value numerically, we find that it exponentially approaches 1 as *n* grows large. While we provide a lower bound on the upper bound on *α*, it remains unclear whether this is a tight lower bound or whether there is a construction that can attain higher values for *α*.

### S4.3 Ridge regression

We recall Lemma S1.4:

#### Lemma S1.4

*For non-adjacent items*,

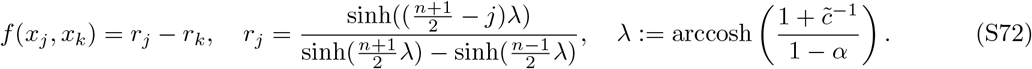

*For adjacent items*,

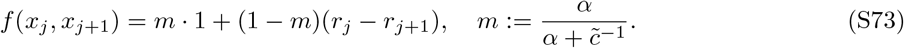

*Proof*. We had established that

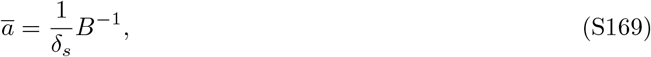

Where

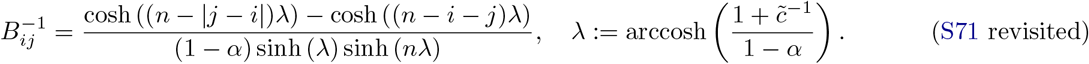

Thus

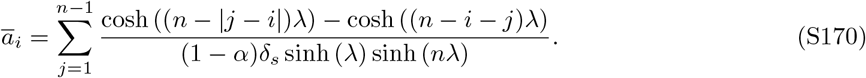

Setting

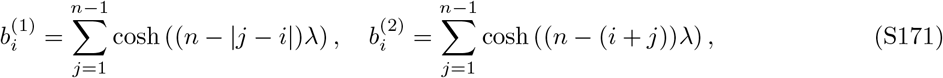

we can express this as

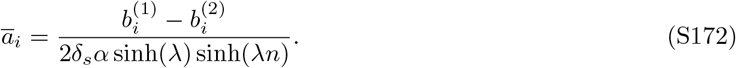

We simplify

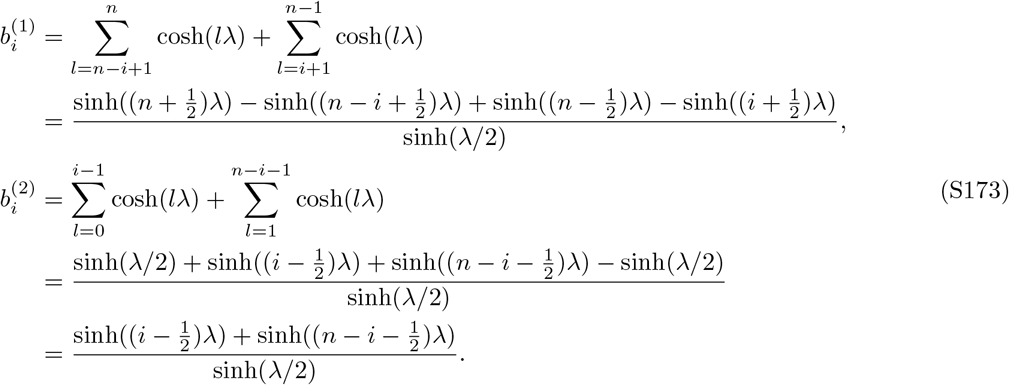

Moreover, note that

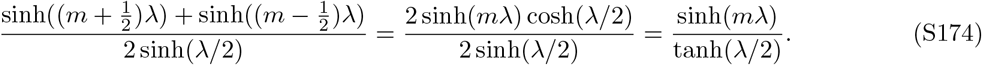

Applying this to *m* ∈ {*n, n* − *i, i*} results in

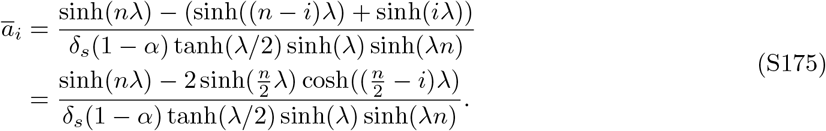

Note that the resulting boundary conditions are *ā*_0_ = *ā*_*n*_ = 0, consistent with our definition.

Recalling that 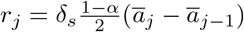, (S69), we find that

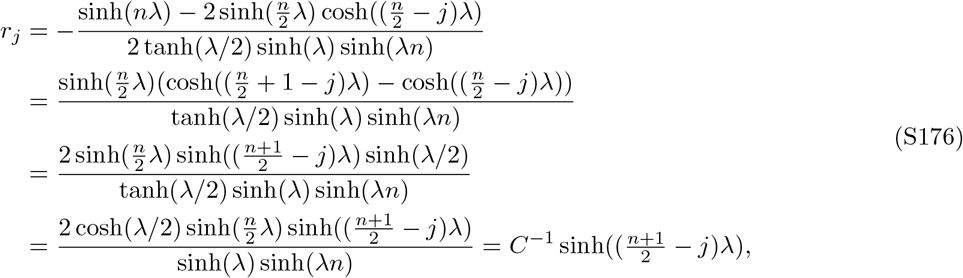

where

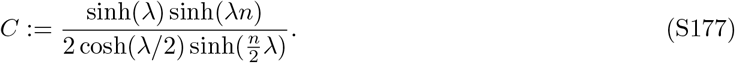

Noting that

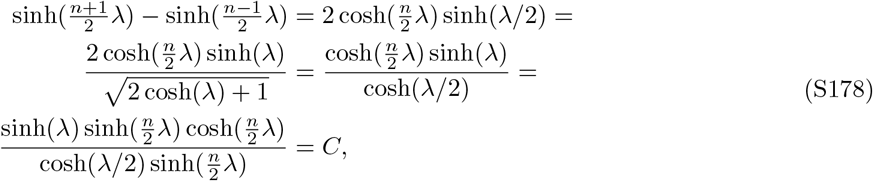

proves (S72).

Analysing the adjacent items, we observe that by definition,

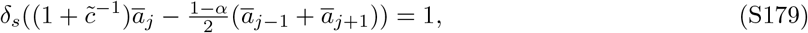

and thus

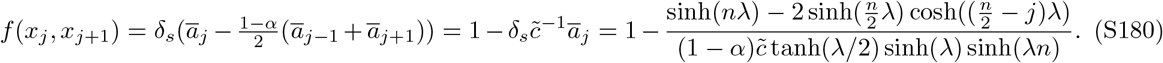

We split this up into two components: the memorization coefficient

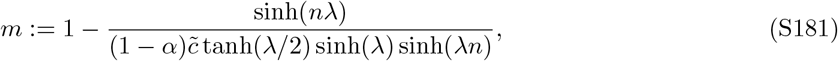

and the generalization coefficient *g*, which we pick such that

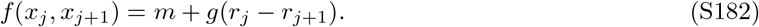

We first simplify

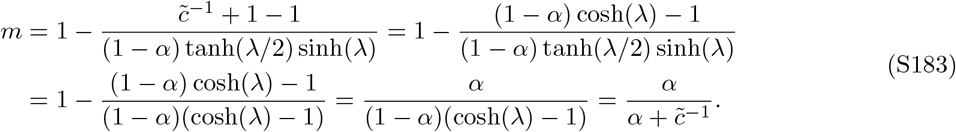

To determine *g*, we solve the equation

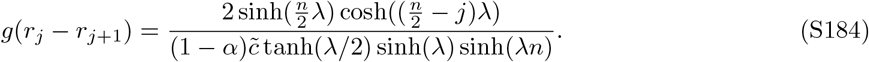

We observe

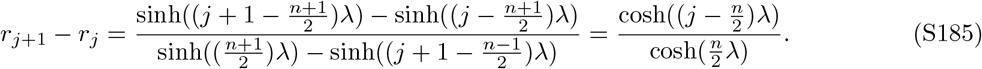

Thus,

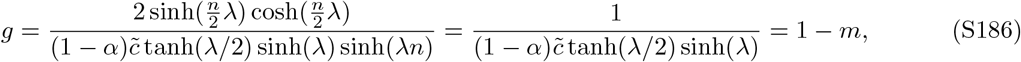

and we can write

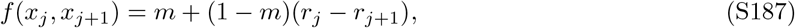

proving (S73).

### S4.4 Gradient descent dynamics for mean squared error

We recall Lemma S1.5:

#### Lemma S1.5

*Given the mean squared error as a loss function, g*_*j*_ *and r*_*j*_ *amount to*

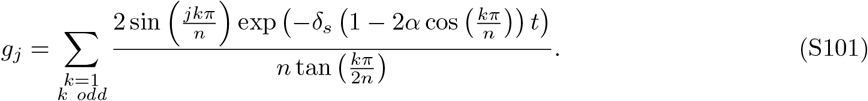

*and*

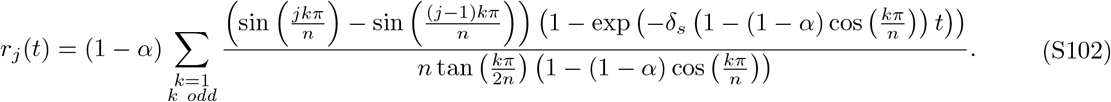

*Proof*. We here solve the differential equation

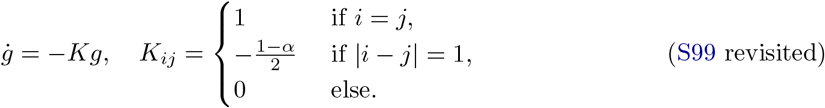

Setting *U* as in Theorem S4.2 (meaning that *K* = *U* Λ*U*), we define

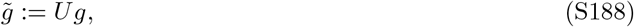

which results in the diagonalized equation

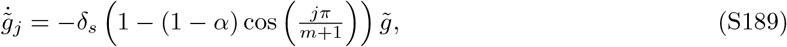

We know that *g*(0) = 1 and so

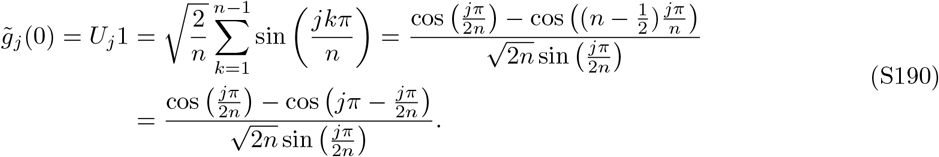

For odd *j*,

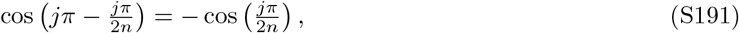

and therefore

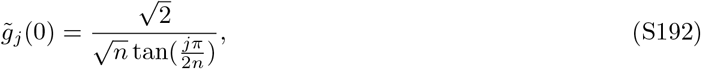

meaning that

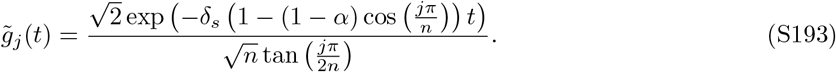

For even *j*,

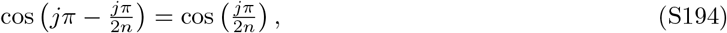

and therefore

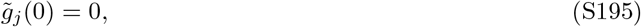

meaning that

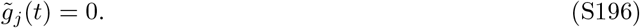

Therefore,

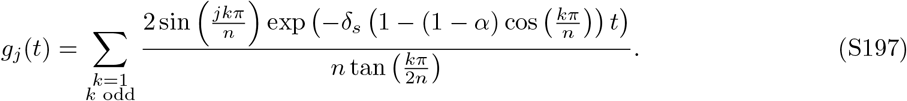

This proves (S101).

As a reminder, the rank representation was defined as

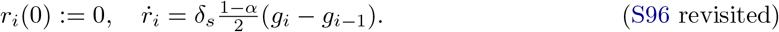

As a result,

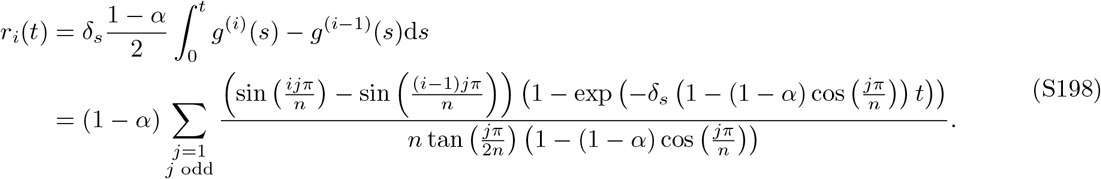

This proves (S102).

### S4.5 Deep neural networks’ conjunctivity factor in the limit of infinite depth

We recall Lemma S1.3:

#### Lemma S1.3

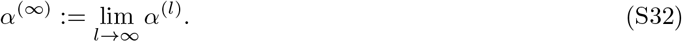

*is given by*

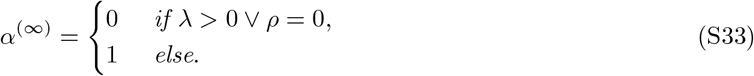

*Proof*. Note that for *ρ* = 0, *ϕ* is the identity function and as a results *α*^(*l*)^ = 0. We thus assume *ρ* ≠ 0. As a reminder,

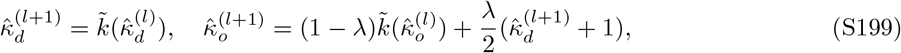

where

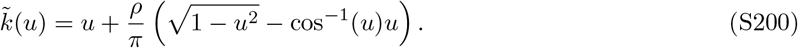

First, 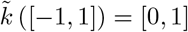. Further, 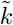 has precisely one fixed point at *u* = 1. As a result,

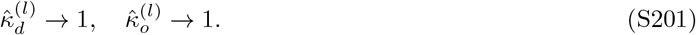

Note that

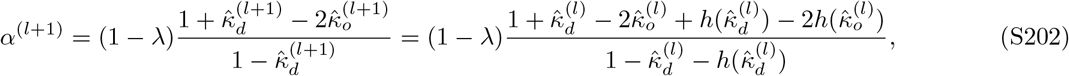

where

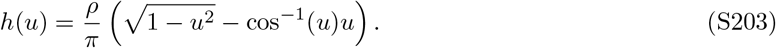

We further simplify

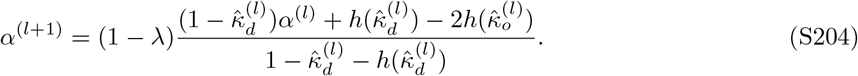

Noting that

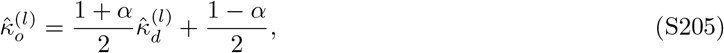

we can define updates to *α*^(*l*)^ by the iterated map

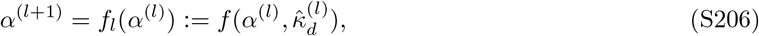

where

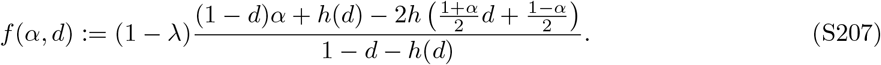

**Case** *λ* = 0. For *λ* = 0, the fixed point equation amounts to

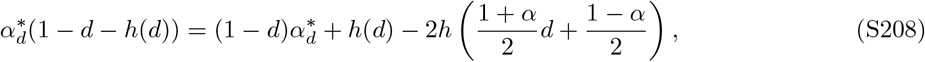

which is equivalent to

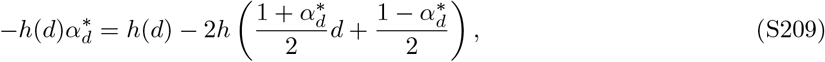

and thus

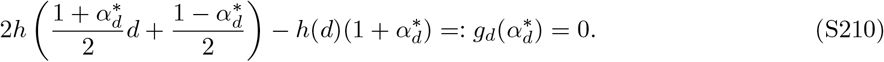

Clearly, *g*_*d*_(1) = 0. Note that

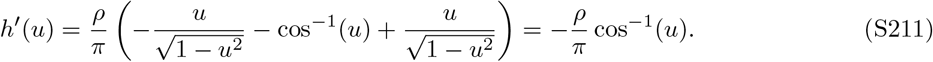

Therefore,

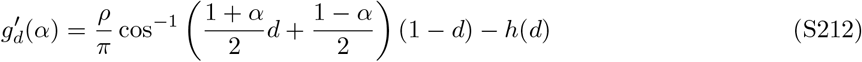

As cos^−1^ is monotonically decreasing,

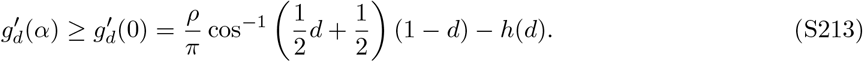

Visual inspection of this plot reveals 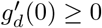 for all *d*. As a result, *g*_*d*_ is monotonically increasing on *α* and can therefore only have one zero. Put differently, the fixed point 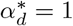 is unique. This means that each *f*_*l*_ has a unique fixed point at 1, and, by Theorem S4.4, *α*^(*l*)^ → 1.

**Case** *λ* > 0. Because *f* (·, *d*) maps [0, 1] onto [0, 1], it has a fixed point. If

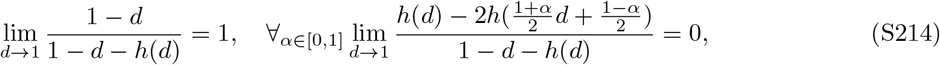

we can write

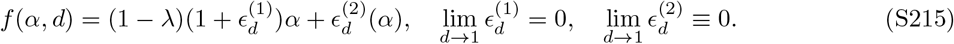

For any fixed point 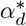, we thus observe that

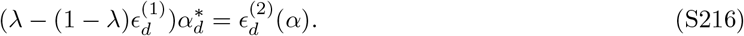

As 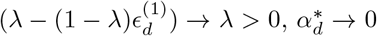. This means that any sequence of fixed points given by *f*_*l*_ as *l* → ∞ must converge to zero, and, as a result, *α*^(*l*)^ → 0.

All that remains to show is (S214). We first note that 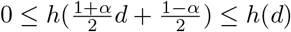 and thus

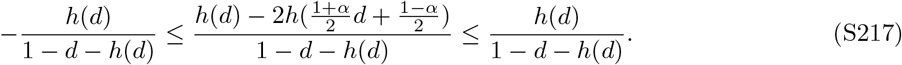

As the left limit in (S214) implies

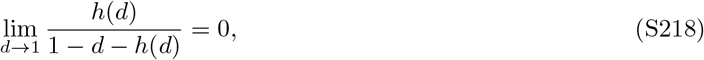

it immediately implies the right half of the equation. All that remains to show is the left half. Note that it is equivalent to

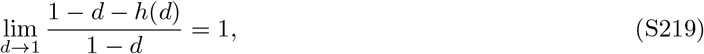

and thus

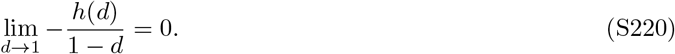

We observe that

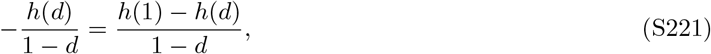

and thus

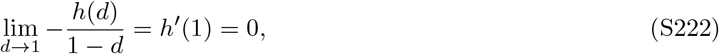

which concludes the proof.

### S4.6 Details on the cooperative coding scheme

We describe the construction of the units with positive readout weights. We make sure that these units are silent on data points with negative labels. The units with negative readout weights are constructed analogously and therefore silent on data points with positive labels. Because the induction step is a bit complicated, we describe the first few cases (*n* = 2, …, 6) concretely. All unit weights will either be zero or 0.5. We give an example of the specified code for *n* = 2, …, 7 in Table 2. Overall, there are five cases to be considered. Our induction step for each case will be that all trials involving the first *n* − 1 items are correctly classified. Note that from now on we will say that some unit *u* needs to encode *x* if we require that ⟨*u, x*⟩. Further, we say that *u* needs to be silent on *x* if we require that ⟨*u, x*⟩ ≤ 0.

**Table 2:**
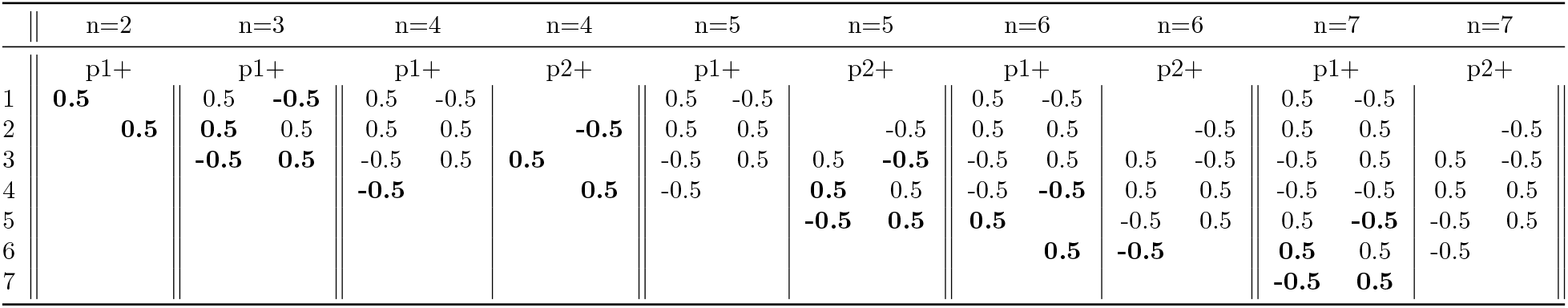
Weights of the hand-constructed network for *n* = 2, …, 7. Rows reference responses to the different units. First column for each unit is response to the item when in the first position, second column is response to the item when in the second position. Entries that are new between consecutive numbers of items are highlighted in bold.

**Case 1:** *n* = 2. H_1+_ needs to encode (1, 2) and be silent on (2, 1). It can do so simply by setting H_1+_(1, 1) = H_1+_(2, 2) = 0.5. Thus, the first unit has two weights added and the second unit has no weights added (as it is not needed at all).

**Case 2:** *n* = 4*k*+3 **(e.g**. *n* = 3, 7). This step adds the trial (4*k*+2, 4*k*+3), which is encoded by H_1+_, (as well as (4*k*+3, 4*k*+2)). Because H_1+_ also encodes (4*k*+1, 4*k*+2) and therefore H_1+_(4*k*+2, 2) = 0.5, it needs to add a negative weight H_1+_(4*k* +3, 1) = −0.5. To encode (4*k* +2, 4*k* +3), we add H_1+_(4*k* +2, 1) = H_1+_(4*k* +3, 2) = 0.5. Finally, to remain silent on (4*k* + 2, 4*k* + 1), we add a negative weight H_1+_(4*k* + 1, 2) = −0.5. In total there are **four weights added to H**_1+_. On the other hand, **H**_2+_ **needs no units added** as it is already silent on both added data points (we can check this by checking the two preceding cases 1 and 5).

**Case 3:** *n* = 4*k* + 4 **(e.g**. *n* = 4, 8). This steps adds the trial (4*k* + 3, 4*k* + 4), which is encoded by H_2+_ (as well as (4*k* + 4, 4*k* + 3)). We therefore need to add H_2+_(4*k* + 3, 1) = *p*_2+_(4*k* + 4, 2) = 0.5. To maintain silence on (4*k* + 3, 4*k* + 2), we also need to add H_2+_(4*k* + 2, 2) = −0.5 (which was not needed so far, as can be checked from case 2). In total, we have added **three weights to H**_2+_. For H_1+_, on the other hand, the positive weight at H_1+_(4*k* + 3, 2) (per case 2) needs to be counter-balanced by setting H_1+_(4*k* + 4, 1) = −0.5. We therefore add **one weight to H**_1+_.

**Case 4:** *n* = 4*k* + 5 **(e.g**. *n* = 5). This step adds the trial (4*k* + 4, 4*k* + 5), which is encoded by H_2+_ (as well as (4*k* + 5, 4*k* + 4)). Note that it is somewhat analogous to case 2 except with the roles of H_1+_ and H_2+_ being exchanged. Specifically, because H_2+_ also encodes (4*k* + 3, 4*k* + 4), we need to add a negative weight H_2+_(4*k* + 5, 1) = −0.5 to maintain silence on (4*k* + 5, 4*k* + 4). Additionally we need to add positive weights H_2+_(4*k* + 4, 1) = H_2+_(4*k* + 5, 2) = 0.5. To maintain silence on (4*k* + 4, 4*k* + 3), we finally need to add H_2+_(4*k* + 3, 2) = −0.5 (which had no weight assigned as per cases 2 and 3). In total we added **four weights to H**_2+_ and **zero weights to H**_1+_.

**Case 5:** *n* = 4*k* + 6 **(e.g**. *n* = 6). This step adds the trial (4*k* + 5, 4*k* + 6), which is encoded by H_1+_ (as well as (4*k* + 6, 4*k* + 5)). Note that it is analogous to case 3 with the roles of H_1+_ and H_2+_ being exchanged and we need to add **three weights to H**_1+_ and **one weight to H**_2+_.

In total, this means that we can define two recursive functions *c*_1_(*n*), *c*_2_(*n*) that count the weights of H_1*±*_ and H_2*±*_, respectively. The baseline conditions are

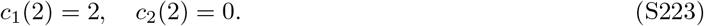

The increment is defined as

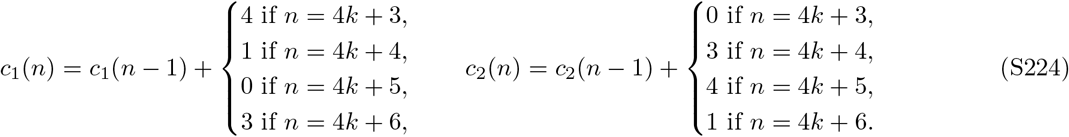

Note that *c*_1_(*n*), *c*_2_(*n*) ≈ 2(*n* − 1).

We can then compute the norm as

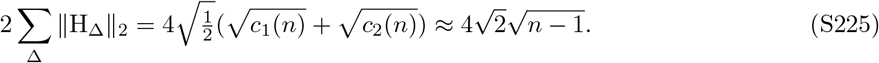

## Footnotes

1 A related paper [55] has come to our attention, in which the authors are concerned with a linear item representation. This is a special case of the additive representations here. Further, the authors derive an analytic solution for a specific learning algorithm, whereas we demonstrate a transitive constraint for a broader class of models.

## References

1. Halford, G. S., Wilson, W. H. & Phillips, S. Relational knowledge: the foundation of higher cognition. en. Trends in Cognitive Sciences 14, 497–505. issn: 1364-6613. https://www.sciencedirect.com /science/article/pii/S1364661310002020(2023) (Nov. 2010).

2. Cheney, D. L., Seyfarth, R. M. & Silk, J. B. The responses of female baboons (Papio cynocephalus ursinus) to anomalous social interactions: Evidence for causal reasoning? Journal of Comparative Psychology 109, 134– 141. issn: 1939-2087(Electronic),0735-7036(Print) (1995).

3. Peake, T. M., Terry, A. M. R., McGregor, P. K. & Dabelsteen, T. Do great tits assess rivals by combining direct experience with information gathered by eavesdropping? Proceedings of the Royal Society of London. Series B: Biological Sciences 269. Publisher: Royal Society, 1925–1929. https://royalsocietypublishing.org/doi/abs/10.1098/rspb.2002.2112(2022) (Sept. 2002).

4. Paz-y-Miño C, G., Bond, A. B., Kamil, A. C. & Balda, R. P. Pinyon jays use transitive inference to predict social dominance. en. Nature 430. Number: 7001 Publisher: Nature Publishing Group, 778–781. issn: 1476-4687. https://www.nature.com/articles/nature02723 (2022) (Aug. 2004).

5. Etienne, A. S. & Jeffery, K. J. Path integration in mammals. en. Hippocampus 14, 180–192. issn: 1364-6613. https://onlinelibrary.wiley.com/doi/abs/10.1002/hipo.10173 (2022) (2004).

6. Seed, A. & Byrne, R. Animal Tool-Use. en. Current Biology 20, R1032–R1039. issn: 1364-6613. https://www.sciencedirect.com/science/article/pii/S0960982210011607(2022) (Dec. 2010).

7. Baldwin, M. W. Relational schemas and the processing of social information. Psychological bulletin 112, 461 (1992).

8. Tolman, E. C. Cognitive maps in rats and men. Psychological Review 55, 189–208. issn: 1939-1471 (1948).

9. Penn, D. C. & Povinelli, D. J. Causal cognition in human and nonhuman animals: A comparative, critical review. Annu. Rev. Psychol. 58. Publisher: Annual Reviews, 97–118 (2007).

10. Mitchell, T. M. The need for biases in learning generalizations (1980).

11. Krogh, A. & Hertz, J. A. Generalization in a linear perceptron in the presence of noise. en. Journal of Physics A: Mathematical and General 25, 1135. issn: 1364-6613. 10.1088/0305-4470/25/5/020(2023) (Mar. 1992).

12. Watkin, T. L. H., Rau, A. & Biehl, M. The statistical mechanics of learning a rule. Reviews of Modern Physics 65. Publisher: American Physical Society, 499–556. doi/10.1103/RevModPhys.65.499(2023) (Apr. 1993).

13. Sollich, P. Learning Curves for Gaussian Processes in Advances in Neural Information Processing Systems 11 (MIT Press, 1998). https://proceedings.neurips.cc/paper_files/paper/1998/hash/5cbdfd0dfa22a3fca7266376887f549b-Abstract.html(2023).

14. Liang, T. & Rakhlin, A. Just interpolate: Kernel “Ridgeless” regression can generalize. The Annals of Statistics 48. Publisher: Institute of Mathematical Statistics, 1329–1347. issn: 1364-6613. https://projecteuclid.org/journals/annals-of-statistics/volume-48/issue-3/Just-interpolate-Kernel-Ridgeless-regression-can-generalize/10.1214/19-AOS1849.full(2023) (June 2020).

15. Jacot, A., Simsek, B., Spadaro, F., Hongler, C. & Gabriel, F. Kernel Alignment Risk Estimator: Risk Prediction from Training Data in Advances in Neural Information Processing Systems 33 (Curran Associates, Inc., 2020), 15568–15578. https://proceedings.neurips.cc/paper/2020/hash/b367e525a7e574817c19ad24b7b35607-Abstract.html(2023).

16. Canatar, A., Bordelon, B. & Pehlevan, C. Spectral bias and task-model alignment explain generalization in kernel regression and infinitely wide neural networks. en. Nature Communications 12, 2914. issn: 1364-6613. https://www.nature.com/articles/s41467-021-23103-1 (2023) (May 2021).

17. Hoerl, A. E. & Kennard, R. W. Ridge Regression: Biased Estimation for Nonorthogonal Problems. Technometrics 12, 55–67. issn: 1364-6613. https://www.tandfonline.com/doi/abs/10.1080/00401706.1970.10488634 (2023) (Feb. 1970).

18. Cortes, C. & Vapnik, V. Support-vector networks. en. Machine Learning 20, 273–297. issn: 1364-6613. 10.1007/BF00994018(2022) (Sept. 1995).

19. Soudry, D., Hoffer, E., Nacson, M. S., Gunasekar, S. & Srebro, N. The implicit bias of gradient descent on separable data. Journal of Machine Learning Research 19, 1–57 (2018).

20. Krogh, A. & Hertz, J. A Simple Weight Decay Can Improve Generalization in Advances in Neural Information Processing Systems 4 (1991). https://proceedings.neurips.cc/paper/1991/hash/8eefcfdf5990e441f0fb6f3fad709e21-Abstract.html(2023).

21. Moody, J. The Effective Number of Parameters: An Analysis of Generalization and Regularization in Nonlinear Learning Systems in Advances in Neural Information Processing Systems 4 (1991). https://proceedings.neurips.cc/paper/1991/hash/d64a340bcb633f536d56e51874281454-Abstract.html(2023).

22. Barnett, S. M. & Ceci, S. J. When and where do we apply what we learn?: A taxonomy for far transfer. Psychological Bulletin 128. Place: US Publisher: American Psychological Association, 612–637. issn: 1939-1455 (2002).

23. Ye, H. et al. Towards a Theoretical Framework of Out-of-Distribution Generalization in Advances in Neural Information Processing Systems 34 (Curran Associates, Inc., 2021), 23519–23531. https://proceedings.neurips.cc/paper/2021/hash/c5c1cb0bebd56ae38817b251ad72bedb-Abstract.html(2023).

24. Abbe, E., Bengio, S., Lotfi, A. & Rizk, K. Generalization on the Unseen, Logic Reasoning and Degree Curriculum June 2023. http://arxiv.org/abs/2301.13105 (2023).

25. Battaglia, P. W. et al. Relational inductive biases, deep learning, and graph networks. arXiv preprint arXiv:1806.01261 (2018).

26. Piaget, J. Judgment and reasoning in the child. Publisher: Harcourt, Brace (1928).

27. Vasconcelos, M. Transitive inference in non-human animals: An empirical and theoretical analysis. en. Behavioural Processes 78, 313–334. issn: 1364-6613. https://www.sciencedirect.com/science/article/pii/S0376635708000818(2022) (July 2008).

28. Jensen, G. in APA handbook of comparative psychology: Perception, learning, and cognition, Vol. 2 385–409 (American Psychological Association, Washington, DC, US, 2017). isbn: 978-1-4338-2352-7978-1-4338-2353-4.

29. Ciranka, S. et al. Asymmetric reinforcement learning facilitates human inference of transitive relations. en. Nature Human Behaviour 6, 555–564. issn: 1364-6613. https://www.nature.com/articles/s41562-021-01263-w(2022) (Apr. 2022).

30. Nelli, S., Braun, L., Dumbalska, T., Saxe, A. & Summerfield, C. Neural knowledge assembly in humans and neural networks. Neuron (2023).

31. Jensen, G., Muñoz, F., Alkan, Y., Ferrera, V. P. & Terrace, H. S. Implicit Value Updating Explains Transitive Inference Performance: The Betasort Model. en. PLOS Computational Biology 11, e1004523. issn: 1364-6613. https://journals.plos.org/ploscompbiol/article?id=10.1371/journal.pcbi.1004523 (2022) (Sept. 2015).

32. Bryant, P. E. & Trabasso, T. Transitive Inferences and Memory in Young Children. en. Nature 232, 456–458. issn: 1476-4687. https://www.nature.com/articles/232456a0 (2022) (Aug. 1971).

33. McGonigle, B. O. & Chalmers, M. Are monkeys logical? Nature 267, 694–696. issn: 1476-4687 (1977).

34. Gillan, D. J. Reasoning in the chimpanzee: II. Transitive inference. Journal of Experimental Psychology: Animal Behavior Processes 7, 150–164. issn: 1939-2184 (1981).

35. Davis, H. Transitive inference in rats (Rattus norvegicus). Journal of Comparative Psychology 106, 342–349. issn: 1939-2087 (1992).

36. Tibbetts, E. A., Agudelo, J., Pandit, S. & Riojas, J. Transitive inference in Polistes paper wasps. Biology Letters 15, 20190015. https://royalsocietypublishing.org/doi/10.1098/rsbl.2019.0015 (2022) (May 2019).

37. Grosenick, L., Clement, T. S. & Fernald, R. D. Fish can infer social rank by observation alone. en. Nature 445, 429–432. issn: 1364-6613. https://www.nature.com/articles/nature05511 (2022) (Jan. 2007).

38. Potts, G. R. Storing and retrieving information about ordered relationships. Journal of Experimental Psychology 103, 431–439. issn: 0022-1015 (1974).

39. D’Amato, M. R. & Colombo, M. The symbolic distance effect in monkeys (Cebus apella). en. Animal Learning & Behavior 18, 133–140. issn: 1364-6613. 10.3758/BF03205250(2022) (June 1990).

40. McGonigle, B. & Chalmers, M. Monkeys are rational! The Quarterly Journal of Experimental Psychology Section B 45, 189–228. issn: 1364-6613. https://www.tandfonline.com/doi/abs/10.1080/14640749208401017 (2022) (Oct. 1992).

41. Merritt, D., MacLean, E. L., Jaffe, S. & Brannon, E. M. A comparative analysis of serial ordering in ring-tailed lemurs (Lemur catta). Journal of Comparative Psychology 121. Publisher: American Psychological Association, 363 (2007).

42. Scarf, D. & Colombo, M. Representation of serial order: A comparative analysis of humans, monkeys, and pigeons. en. Brain Research Bulletin. Special Issue:Brain Mechanisms, Cognition and Behaviour in Birds 76, 307–312. issn: 1364-6613. https://www.sciencedirect.com/science/article/pii/S0361923008000373 (2022) (June 2008).

43. Von Fersen, L., Wynne, C. D., Delius, J. D. & Staddon, J. E. Transitive inference formation in pigeons. Journal of Experimental Psychology: Animal Behavior Processes 17. Publisher: American Psychological Association, 334 (1991).

44. Wynne, C. D. L. Pigeon transitive inference: Tests of simple accounts of a complex performance. en. Behavioural Processes 39, 95–112. issn: 1364-6613. https://www.sciencedirect.com/science/article/pii/S0376635796000484(2022) (Jan. 1997).

45. Merritt, D. J. & Terrace, H. S. Mechanisms of inferential order judgments in humans (Homo sapiens) and rhesus monkeys (Macaca mulatta). Journal of Comparative Psychology 125. Publisher: American Psychological Association, 227 (2011).

46. Acuna, B. D., Sanes, J. N. & Donoghue, J. P. Cognitive mechanisms of transitive inference. en. Experimental Brain Research 146, 1–10. issn: 1364-6613. 10.1007/s00221-002-1092-y(2022) (Sept. 2002).

47. Van Elzakker, M., O’Reilly, R. C. & Rudy, J. W. Transitivity, flexibility, conjunctive representations, and the hippocampus. I. An empirical analysis. en. Hippocampus 13, 334–340. issn: 1364-6613. https://onlinelibrary.wiley.com/doi/abs/10.1002/hipo.10083(2022) (2003).

48. Wynne, C. D. L. in Models of Action Num Pages: 40 (Psychology Press, 1998). isbn: 978-0-203-77386-4.

49. Wynne, C. D. L. Reinforcement accounts for transitive inference performance. en. Animal Learning & Behavior 23, 207–217. issn: 1364-6613. 10.3758/BF03199936(2022) (June 1995).

50. Couvillon, P. A. & Bitterman, M. E. A conventional conditioning analysis of “transitive inference” in pigeons. Journal of Experimental Psychology: Animal Behavior Processes 18. Place: US Publisher: American Psychological Association, 308–310. issn: 1939-2184 (1992).

51. Jensen, G., Terrace, H. S. & Ferrera, V. P. Discovering Implied Serial Order Through Model-Free and Model-Based Learning. Frontiers in Neuroscience 13. issn: 1364-6613. https://www.frontiersin.org/articles/10.3389/fnins.2019.00878(2022) (2019).

52. Wynne, C. D. L. Transverse patterning in pigeons. en. Behavioural Processes 38, 119–130. issn: 1364-6613. https://www.sciencedirect.com/science/article/pii/S0376635796000320 (2022) (Nov. 1996).

53. De Lillo, C., Floreano, D. & Antinucci, F. Transitive choices by a simple, fully connected, backpropagation neural network: implications for the comparative study of transitive inference. en. Animal Cognition 4, 61–68. issn: 1364-6613. 10.1007/s100710100092 (2022) (July 2001).

54. Kay, K. et al. Neural dynamics and geometry for transitive inference en. Pages: 2022.10.10.511448 Section: New Results. Oct. 2022. https://www.biorxiv.org/content/10.1101/2022.10.10.511448v1 (2022).

55. Antonio, G. D., Raglio, S. & Mattia, M. Ranking and serial thinking: A geometric solution en. Pages: 2023.08.03.551859 Section: New Results. Aug. 2023. https://www.biorxiv.org/content/10.1101/2023.08.03.551859v1 (2023).

56. Carmesin, H. .-. & Schwegler, H. Parallel versus sequential processing of relational stimulus structures. en. Biological Cybernetics 71, 523–529. issn: 1364-6613. 10.1007/BF00198470(2022) (Oct. 1994).

57. Bogacz, R., Brown, E., Moehlis, J., Holmes, P. & Cohen, J. D. The physics of optimal decision making: a formal analysis of models of performance in two-alternative forced-choice tasks. eng. Psychological Review 113, 700– 765. issn: 0033-295X (Oct. 2006).

58. Gold, J. I. & Shadlen, M. N. The Neural Basis of Decision Making. Annual Review of Neuroscience 30. eprint: 10.1146/annurev.neuro.29.051605.113038, 535–574. 10.1146/annurev.neuro.29.051605.113038(2023) (2007).

59. Churchland, A. K. et al. Variance as a Signature of Neural Computations during Decision Making. English. Neuron 69. Publisher: Elsevier, 818–831. issn: 1364-6613. https://www.cell.com/neuron/abstract/S0896-6273(10)01087-1(2023) (Feb. 2011).

60. Ratcliff, R. A theory of memory retrieval. Psychological Review 85, 59–108. issn: 1939-1471 (1978).

61. Ratcliff, R. & McKoon, G. The Diffusion Decision Model: Theory and Data for Two-Choice Decision Tasks. Neural Computation 20, 873–922. issn: 1364-6613. 10.1162/neco.2008.12-06-420(2023) (Apr. 2008).

62. Frankland, S. M. & Greene, J. D. Two Ways to Build a Thought: Distinct Forms of Compositional Semantic Representation across Brain Regions. Cerebral Cortex 30, 3838–3855. issn: 1364-6613. 10.1093/cercor/bhaa001(2023) (May 2020).

63. Rigotti, M. et al. The importance of mixed selectivity in complex cognitive tasks. en. Nature 497. Number: 7451 Publisher: Nature Publishing Group, 585–590. issn: 1364-6613. https://www.nature.com/articles/nature12160(2022) (May 2013).

64. Rudy, J. W. & O’Reilly, R. C. Conjunctive representations, the hippocampus, and contextual fear conditioning. en. Cognitive, Affective, & Behavioral Neuroscience 1, 66–82. issn: 1364-6613. 10.3758/CABN.1.1.66(2023) (Mar. 2001).

65. Rescorla, R. A. “Configural” conditioning in discrete-trial bar pressing. eng. Journal of Comparative and Physiological Psychology 79, 307–317. issn: 0021-9940 (May 1972).

66. Mante, V., Sussillo, D., Shenoy, K. V. & Newsome, W. T. Context-dependent computation by recurrent dynamics in prefrontal cortex. en. Nature 503, 78–84. issn: 1364-6613. https://www.nature.com/articles/nature12742(2023) (Nov. 2013).

67. Komorowski, R. W., Manns, J. R. & Eichenbaum, H. Robust Conjunctive Item–Place Coding by Hippocampal Neurons Parallels Learning What Happens Where. en. Journal of Neuroscience 29. Publisher: Society for Neuroscience Section: Articles, 9918–9929. issn: 0270-6474, 1529-2401. https://www.jneurosci.org/content/29/31/9918(2023) (Aug. 2009).

68. McKenzie, S. et al. Hippocampal Representation of Related and Opposing Memories Develop within Distinct, Hierarchically Organized Neural Schemas. English. Neuron 83. Publisher: Elsevier, 202–215. issn: 1364-6613. https://www.cell.com/neuron/abstract/S0896-6273(14)00405-X (2023) (July 2014).

69. Fusi, S., Miller, E. K. & Rigotti, M. Why neurons mix: high dimensionality for higher cognition. en. Current Opinion in Neurobiology. Neurobiology of cognitive behavior 37, 66–74. issn: 1364-6613. https://www.sciencedirect.com/science/article/pii/S0959438816000118(2023) (Apr. 2016).

70. Davis, H. in Cognitive Aspects of Stimulus Control (Psychology Press, 1992). isbn: 978-1-315-78910-1.

71. Cho, Y. & Saul, L. Kernel Methods for Deep Learning in Advances in Neural Information Processing Systems 22 (2009). https://papers.nips.cc/paper/2009/hash/5751ec3e9a4feab575962e78e006250d-Abstract.html (2022).

72. Turrigiano, G. G. The Self-Tuning Neuron: Synaptic Scaling of Excitatory Synapses. English. Cell 135. Publisher: Elsevier, 422–435. issn: 1364-6613. https://www.cell.com/cell/abstract/S0092-8674(08)01298-1(2023) (Oct. 2008).

73. Saxe, A., McClelland, J. & Ganguli, S. Exact solutions to the nonlinear dynamics of learning in deep linear neural networks in International Conference on Learning Representations (2014).

74. Gunasekar, S., Lee, J., Soudry, D. & Srebro, N. Characterizing implicit bias in terms of optimization geometry in International Conference on Machine Learning (PMLR, 2018), 1832–1841.

75. Ji, Z., Dudík, M., Schapire, R. E. & Telgarsky, M. Gradient descent follows the regularization path for general losses in Conference on Learning Theory (PMLR, 2020), 2109–2136.

76. Spence, K. W. The nature of the response in discrimination learning. Psychological Review 59, 89–93. issn: 1939-1471 (1952).

77. Alvarado, M. C. & Rudy, J. W. Some properties of configural learning: An investigation of the transverse-patterning problem. Journal of Experimental Psychology: Animal Behavior Processes 18. Place: US Publisher: American Psychological Association, 145–153. issn: 1939-2184 (1992).

78. Dusek, J. A. & Eichenbaum, H. The hippocampus and transverse patterning guided by olfactory cues. Behavioral Neuroscience 112, 762–771. issn: 1939-0084 (1998).

79. Gao, J., Su, Y., Tomonaga, M. & Matsuzawa, T. Learning the rules of the rock–paper–scissors game: chimpanzees versus children. en. Primates 59, 7–17. issn: 1364-6613. 10.1007/s10329-017-0620-0(2022) (Jan. 2018).

80. LeCun, Y., Bengio, Y. & Hinton, G. Deep learning. en. Nature 521, 436–444. issn: 1364-6613. https://www.nature.com/articles/nature14539(2023) (May 2015).

81. Richards, B. A. et al. A deep learning framework for neuroscience. en. Nature Neuroscience 22, 1761–1770. issn: 1364-6613. https://www.nature.com/articles/s41593-019-0520-2 (2023) (Nov. 2019).

82. Saxe, A., Nelli, S. & Summerfield, C. If deep learning is the answer, what is the question? en. Nature Reviews Neuroscience 22, 55–67. issn: 1364-6613. https://www.nature.com/articles/s41583-020-00395-8 (2023) (Jan. 2021).

83. Chizat, L., Oyallon, E. & Bach, F. On Lazy Training in Differentiable Programming in Advances in Neural Information Processing Systems 32 (2019). https://proceedings.neurips.cc/paper/2019/hash/ae614c557843b1df326cb29c57225459-Abstract.html(2023).

84. Woodworth, B. et al. Kernel and Rich Regimes in Overparametrized Models en. in Conference on Learning Theory (July 2020), 3635–3673. https://proceedings.mlr.press/v125/woodworth20a.html (2023).

85. Jacot, A., Gabriel, F. & Hongler, C. Neural tangent kernel: Convergence and generalization in neural networks.Advances in neural information processing systems 31 (2018).

86. Lee, J. et al. Wide neural networks of any depth evolve as linear models under gradient descent. Advances in neural information processing systems 32 (2019).

87. Fort, S. et al. Deep learning versus kernel learning: an empirical study of loss landscape geometry and the time evolution of the Neural Tangent Kernel in Advances in Neural Information Processing Systems 33 (2020), 5850–5861. https://proceedings.neurips.cc/paper/2020/hash/405075699f065e43581f27d67bb68478-Abstract.html(2023).

88. Vyas, N., Bansal, Y. & Nakkiran, P. Limitations of the NTK for Understanding Generalization in Deep Learning arXiv:2206.10012 [cs]. June 2022. http://arxiv.org/abs/2206.10012 (2023).

89. Weinberger, N. Dynamic regulation of receptive fields and maps in the adult sensory cortex. Annual review of neuroscience 18, 129–158 (1995).

90. Poort, J. et al. Learning Enhances Sensory and Multiple Non-sensory Representations in Primary Visual Cortex. en. Neuron 86, 1478–1490. issn: 1364-6613. https://www.sciencedirect.com/science/article/pii/S0896627315004766(2023) (June 2015).

91. Flesch, T., Juechems, K., Dumbalska, T., Saxe, A. & Summerfield, C. Orthogonal representations for robust context-dependent task performance in brains and neural networks. en. Neuron 110, 1258–1270.e11. issn: 1364-6613. https://www.sciencedirect.com/science/article/pii/S0896627322000058 (2023) (Apr. 2022).

92. Van Opstal, F., Fias, W., Peigneux, P. & Verguts, T. The neural representation of extensively trained ordered sequences. en. NeuroImage 47, 367–375. issn: 1364-6613. https://www.sciencedirect.com/science/article/pii/S1053811909003838(2023) (Aug. 2009).

93. Hinton, E. C., Dymond, S., von Hecker, U. & Evans, C. J. Neural correlates of relational reasoning and the symbolic distance effect: involvement of parietal cortex. en. Neuroscience 168, 138–148. issn: 1364-6613. https://www.sciencedirect.com/science/article/pii/S0306452210004471 (2023) (June 2010).

94. Constantinescu, A. O., O’Reilly, J. X. & Behrens, T. E. J. Organizing conceptual knowledge in humans with a gridlike code. Science 352, 1464–1468. https://www.science.org/doi/full/10.1126/science.aaf0941 (2023) (June 2016).

95. Munoz, F. et al. Learned representation of implied serial order in posterior parietal cortex. Scientific Reports 10. Publisher: Springer, 1–14 (2020).

96. Ramawat, S. et al. Different Contribution of the Monkey Prefrontal and Premotor Dorsal Cortex in Decision Making During a Transitive Inference Task. en. Neuroscience 485, 147–162. issn: 1364-6613. https://www.sciencedirect.com/science/article/pii/S0306452222000276(2023) (Mar. 2022).

97. Whittington, J. C. et al. The Tolman-Eichenbaum machine: unifying space and relational memory through generalization in the hippocampal formation. Cell 183, 1249–1263 (2020).

98. Nacson, M. S., Gunasekar, S., Lee, J., Srebro, N. & Soudry, D. Lexicographic and depth-sensitive margins in homogeneous and non-homogeneous deep models in International Conference on Machine Learning (PMLR, 2019), 4683–4692.

99. Lyu, K. & Li, J. Gradient Descent Maximizes the Margin of Homogeneous Neural Networks in International Conference on Learning Representations (2020). https://openreview.net/forum?id=SJeLIgBKPS.

100. Chizat, L. & Bach, F. Implicit Bias of Gradient Descent for Wide Two-layer Neural Networks Trained with the Logistic Loss en. in Proceedings of Thirty Third Conference on Learning Theory ISSN: 2640-3498 (PMLR, July 2020), 1305–1338. https://proceedings.mlr.press/v125/chizat20a.html (2023).

101. Razin, N. & Cohen, N. Implicit Regularization in Deep Learning May Not Be Explainable by Norms in Advances in Neural Information Processing Systems 33 (2020), 21174–21187. https://proceedings.neurips.cc/paper/2020/hash/f21e255f89e0f258accbe4e984eef486-Abstract.html(2024).

102. Savarese, P., Evron, I., Soudry, D. & Srebro, N. How do infinite width bounded norm networks look in function space? in Proceedings of the Thirty-Second Conference on Learning Theory 99 (June 2019), 2667–2690. https://proceedings.mlr.press/v99/savarese19a.html.

103. Ongie, G., Willett, R., Soudry, D. & Srebro, N. A Function Space View of Bounded Norm Infinite Width ReLU Nets: The Multivariate Case in International Conference on Learning Representations (2020).

104. Glymour, C. Learning, prediction and causal Bayes nets. en. Trends in Cognitive Sciences 7, 43–48. issn: 1364-6613. https://www.sciencedirect.com/science/article/pii/S1364661302000098 (2022) (Jan. 2003).

105. Kemp, C. & Tenenbaum, J. B. The discovery of structural form. Proceedings of the National Academy of Sciences 105, 10687–10692. https://www.pnas.org/doi/10.1073/pnas.0802631105 (2022) (Aug. 2008).

106. Tenenbaum, J. B., Kemp, C., Griffiths, T. L. & Goodman, N. D. How to Grow a Mind: Statistics, Structure, and Abstraction. Science 331. Publisher: American Association for the Advancement of Science, 1279–1285. 10.1126/science.1192788(2022) (Mar. 2011).

107. Jacobs, L. F. From movement to transitivity: the role of hippocampal parallel maps in configural learning. eng. Reviews in the Neurosciences 17, 99–109. issn: 0334-1763 (2006).

108. De Soto, C. B., London, M. & Handel, S. Social reasoning and spatial paralogic. Journal of Personality and Social Psychology 2, 513–521. issn: 1939-1315 (1965).

109. Wilson, R. C., Takahashi, Y. K., Schoenbaum, G. & Niv, Y. Orbitofrontal Cortex as a Cognitive Map of Task Space. en. Neuron 81, 267–279. issn: 1364-6613. https://www.sciencedirect.com/science/article/pii/S0896627313010398(2022) (Jan. 2014).

110. Behrens, T. E. J. et al. What Is a Cognitive Map? Organizing Knowledge for Flexible Behavior. en. Neuron 100, 490–509. issn: 1364-6613. https://www.sciencedirect.com/science/article/pii/S0896627318308560 (2022) (Oct. 2018).

111. McClelland, J. L. et al. Letting structure emerge: connectionist and dynamical systems approaches to cognition. en. Trends in Cognitive Sciences 14, 348–356. issn: 1364-6613. https://www.sciencedirect.com/science/article/pii/S1364661310001245(2022) (Aug. 2010).

112. Griffiths, T. L., Chater, N., Kemp, C., Perfors, A. & Tenenbaum, J. B. Probabilistic models of cognition: exploring representations and inductive biases. English. Trends in Cognitive Sciences 14. Publisher: Elsevier, 357–364. issn: 1364-6613. https://www.cell.com/trends/cognitive-sciences/abstract/S1364-6613(10)00112-9(2022) (Aug. 2010).

113. Saxe, A. M., McClelland, J. L. & Ganguli, S. A mathematical theory of semantic development in deep neural networks. Proceedings of the National Academy of Sciences 116, 11537–11546 (2019).

114. McClelland, J. L., McNaughton, B. L. & O’Reilly, R. C. Why there are complementary learning systems in the hippocampus and neocortex: Insights from the successes and failures of connectionist models of learning and memory. Psychological Review 102, 419–457. issn: 1939-1471 (1995).

115. Kumaran, D., Hassabis, D. & McClelland, J. L. What Learning Systems do Intelligent Agents Need? Complementary Learning Systems Theory Updated. English. Trends in Cognitive Sciences 20, 512–534. issn: 1364-6613, 1879-307X. https://www.cell.com/trends/cognitive-sciences/abstract/S1364-6613(16)30043-2 (2024) (July 2016).

116. Jensen, G., Munoz, F., Meaney, A., Terrace, H. S. & Ferrera, V. P. Transitive inference after minimal training in rhesus macaques (Macaca mulatta). Journal of Experimental Psychology: Animal Learning and Cognition 47, 464 (2021).

117. Zeithamova, D., Schlichting, M. & Preston, A. The hippocampus and inferential reasoning: building memories to navigate future decisions. Frontiers in Human Neuroscience 6. issn: 1364-6613. https://www.frontiersin.org/articles/10.3389/fnhum.2012.00070(2024) (2012).

118. Shohamy, D. & Daw, N. D. Integrating memories to guide decisions. Current Opinion in Behavioral Sciences 5, 85–90 (2015).

119. Tambini, A. & Davachi, L. Awake Reactivation of Prior Experiences Consolidates Memories and Biases Cognition. eng. Trends in Cognitive Sciences 23, 876–890. issn: 1879-307X (Oct. 2019).

120. Kurth-Nelson, Z. et al. Replay and compositional computation. Neuron 111, 454–469 (2023).

121. Kumaran, D. & McClelland, J. L. Generalization through the recurrent interaction of episodic memories: A model of the hippocampal system. Psychological Review 119, 573–616. issn: 1939-1471(Electronic),0033-295X(Print) (2012).

122. Russin, J., Zolfaghar, M., Park, S. A., Boorman, E. & O’Reilly, R. C. Complementary Structure-Learning Neural Networks for Relational Reasoning. CogSci … Annual Conference of the Cognitive Science Society. Cognitive Science Society (U.S.). Conference 2021, 1560–1566. https://www.ncbi.nlm.nih.gov/pmc/articles/PMC8491570/(2024) (July 2021).

123. Miconi, T. & Kay, K. An active neural mechanism for relational learning and fast knowledge reassembly. bioRxiv. https://www.biorxiv.org/content/early/2023/09/04/2023.07.27.550739 (2023).

124. Wang, J. X. et al. Prefrontal cortex as a meta-reinforcement learning system. Nature Neuroscience 21, 860–868 (2018).

125. Whittington, J. C. R., McCaffary, D., Bakermans, J. J. W. & Behrens, T. E. J. How to build a cognitive map. en. Nature Neuroscience 25, 1257–1272. issn: 1364-6613. https://www.nature.com/articles/s41593-022-01153-y(2024) (Oct. 2022).

126. Ambrogioni, L. & Ólafsdottir, H. F. Rethinking the hippocampal cognitive map as a meta-learning computational module. English. Trends in Cognitive Sciences 27, 702–712. issn: 1364-6613. https://www.cell.com/trends/cognitive-sciences/abstract/S1364-6613(23)00128-6(2024) (Aug. 2023).

127. Winocur, G., Moscovitch, M. & Bontempi, B. Memory formation and long-term retention in humans and animals: Convergence towards a transformation account of hippocampal–neocortical interactions. Neuropsychologia. Animal Models of Amnesia 48, 2339–2356. issn: 1364-6613. https://www.sciencedirect.com/science/article/pii/S0028393210001624(2024) (July 2010).

128. Ramawat, S. et al. The transitive inference task to study the neuronal correlates of memory-driven decision making: A monkey neurophysiology perspective. Neuroscience & Biobehavioral Reviews 152, 105258. issn: 1364-6613. https://www.sciencedirect.com /science/article/pii/S0149763423002270(2024) (Sept. 2023).

129. Telgarsky, M. & Singer, Y. A Primal-Dual Convergence Analysis of Boosting. Journal of Machine Learning Research 13 (2012).

130. Kumar, A. et al. DR3: Value-Based Deep Reinforcement Learning Requires Explicit Regularization in International Conference on Learning Representations (2022). https://openreview.net/forum?id=POvMvLi91f.

131. Bond, A. B., Kamil, A. C. & Balda, R. P. Social complexity and transitive inference in corvids. en. Animal Behaviour 65, 479–487. issn: 1364-6613. https://www.sciencedirect.com/science/article/pii/S0003347203921014(2022) (Mar. 2003).

132. MacLean, E. L., Merritt, D. J. & Brannon, E. M. Social complexity predicts transitive reasoning in prosimian primates. en. Animal Behaviour 76, 479–486. issn: 1364-6613. https://www.sciencedirect.com/science/article/pii/S0003347208002030(2023) (Aug. 2008).

133. Harris, M. & McGonigle, B. A model of transitive choice. The Quarterly Journal of Experimental Psychology Section B 47. Publisher: Routledge eprint: https://www.tandfonline.com/doi/pdf/10.1080/14640749408401362,319-348. xissn: 1364-6613. https://www.tandfonline.com/doi/abs/10.1080/14640749408401362 (2023) (Aug. 1994).

134. Bryson, J. J. & Leong, J. C. S. Primate errors in transitive ‘inference’: a two-tier learning model. en. Animal Cognition 10, 1–15. issn: 1364-6613. 10.1007/s10071-006-0024-9(2023) (Jan. 2007).

135. Halford, G. S. Can young children integrate premises in transitivity and serial order tasks? Cognitive Psychology 16, 65–93. issn: 1364-6613. https://www.sciencedirect.com/science/article/pii/0010028584900045 (2023) (Jan. 1984).

136. Treichler, F. R. & Van Tilburg, D. Concurrent conditional discrimination tests of transitive inference by macaque monkeys: list linking. Journal of Experimental Psychology: Animal Behavior Processes 22, 105 (1996).

137. Gazes, R. P., Chee, N. W. & Hampton, R. R. Cognitive mechanisms for transitive inference performance in rhesus monkeys: Measuring the influence of associative strength and inferred order. Journal of Experimental Psychology: Animal Behavior Processes 38. Place: US Publisher: American Psychological Association, 331–345. issn: 1939-2184(Electronic),0097-7403(Print) (2012).

138. Lazareva, O. F. & Wasserman, E. A. Transitive inference in pigeons: Measuring the associative values of Stimuli B and D. en. Behavioural Processes 89, 244–255. issn: 1364-6613. https://www.sciencedirect.com/science/article/pii/S0376635711002403(2023) (Mar. 2012).

139. Jensen, G., Alkan, Y., Muñoz, F., Ferrera, V. P. & Terrace, H. S. Transitive inference in humans (Homo sapiens) and rhesus macaques (Macaca mulatta) after massed training of the last two list items. Journal of Comparative Psychology 131. Place: US Publisher: American Psychological Association, 231–245. issn: 1939-2087(Electronic),0735-7036(Print) (2017).

140. Eichenbaum, H. The cognitive neuroscience of memory: An introduction (Oxford University Press, 2011).

141. Kriegeskorte, N., Mur, M. & Bandettini, P. A. Representational similarity analysis — connecting the branches of systems neuroscience. Frontiers in systems neuroscience, 4 (2008).

142. Dusek, J. A. & Eichenbaum, H. The hippocampus and memory for orderly stimulus relations. Proceedings of the National Academy of Sciences 94. Publisher: National Acad Sciences, 7109–7114 (1997).

143. Basile, B. M., Templer, V. L., Gazes, R. P. & Hampton, R. R. Preserved visual memory and relational cognition performance in monkeys with selective hippocampal lesions. Science Advances 6, eaaz0484. 10.1126/sciadv.aaz0484(2024) (July 2020).

144. Bussey, T. J., Warburton, E. C., Aggleton, J. P. & Muir, J. L. Fornix Lesions Can Facilitate Acquisition of the Transverse Patterning Task: A Challenge for “Configural” Theories of Hippocampal Function. en. Journal of Neuroscience 18. Publisher: Society for Neuroscience Section: ARTICLE, 1622–1631. issn: 0270-6474, 1529-2401. https://www.jneurosci.org/content/18/4/1622 (2023) (Feb. 1998).

145. Alvarado, M. C. & Rudy, J. W. Rats with damage to the hippocampal-formation are impaired on the transverse-patterning problem but not on elemental discriminations. Behavioral Neuroscience 109. Place: US Publisher: American Psychological Association, 204–211. issn: 1939-0084 (1995).

146. Rickard, T. C. & Grafman, J. Losing Their Configural Mind: Amnesic Patients Fail on Transverse Patterning. Journal of Cognitive Neuroscience 10, 509–524. issn: 1364-6613. 10.1162/089892998562915(2022) (July 1998).

147. Alvarado, M. C. & Bachevalier, J. Selective neurotoxic damage to the hippocampal formation impairs performance of the transverse patterning and location memory tasks in rhesus macaques. fr. Hippocampus 15. eprint: 10.1002/hipo.20037, 118–131. issn: 1364-6613. https://onlinelibrary.wiley.com/doi/abs/10.1002/hipo.20037(2022) (2005).

148. Fodor, J. A. & Pylyshyn, Z. W. Connectionism and cognitive architecture: A critical analysis. Cognition 28, 3–71 (1988).

149. Lake, B. M., Salakhutdinov, R. & Tenenbaum, J. B. Human-level concept learning through probabilistic program induction. Science 350, 1332–1338. https://www.science.org/doi/10.1126/science.aab3050 (2022) (Dec. 2015).

150. Shalev-Shwartz, S. & Shashua, A. On the Sample Complexity of End-to-end Training vs. Semantic Abstraction Training arXiv:1604.06915 [cs]. Apr. 2016. http://arxiv.org/abs/1604.06915 (2022).

151. Bender, E. M. & Koller, A. Climbing towards NLU: On Meaning, Form, and Understanding in the Age of Data in Proceedings of the 58th Annual Meeting of the Association for Computational Linguistics July 2020), 5185–5198. https://aclanthology.org/2020.acl-main.463(2022).

152. Klinger, T. et al. A study of compositional generalization in neural models. arXiv preprint arXiv:2006.09437 (2020).

153. Summerfield, C., Luyckx, F. & Sheahan, H. Structure learning and the posterior parietal cortex. en. Progress in Neurobiology 184, 101717. issn: 1364-6613. https://www.sciencedirect.com/science/article/pii/S0301008219303351(2023) (Jan. 2020).

154. Yuille, A. L. & Liu, C. Deep Nets: What have They Ever Done for Vision? en. International Journal of Computer Vision 129, 781–802. issn: 1364-6613. 10.1007/s11263-020-01405-z(2022) (Mar. 2021).

155. Kaplan, J. et al. Scaling laws for neural language models. arXiv preprint arXiv:2001.08361 (2020).

156. Brown, T. et al. Language Models are Few-Shot Learners in Advances in Neural Information Processing Systems 33 (Curran Associates, Inc., 2020), 1877–1901. https://papers.nips.cc/paper/2020/hash/1457c0d6bfcb4967418bfb8ac142f64a-Abstract.html(2022).

157. Mahowald, K. et al. Dissociating language and thought in large language models: a cognitive perspective arXiv:2301.06627 [cs]. Jan. 2023. http://arxiv.org/abs/2301.06627 (2023).

158. Saphra, N. & Lopez, A. Understanding learning dynamics of language models with SVCCA. arXiv preprint arXiv:1811.00225 (2018).

159. Tirumala, K., Markosyan, A. H., Zettlemoyer, L. & Aghajanyan, A. Memorization Without Overfitting: Analyzing the Training Dynamics of Large Language Models arXiv:2205.10770 [cs]. Nov. 2022. http://arxiv.org/abs/2205.10770(2023).

160. Kandpal, N., Deng, H., Roberts, A., Wallace, E. & Raffel, C. Large language models struggle to learn long-tail knowledge in International Conference on Machine Learning (PMLR, 2023), 15696–15707.

161. Grosse, R. et al. Studying Large Language Model Generalization with Influence Functions arXiv:2308.03296 [cs, stat]. Aug. 2023. http://arxiv.org/abs/2308.03296 (2023).

162. Smolensky, P. On the proper treatment of connectionism. en. Behavioral and Brain Sciences 11, 1–23. issn: 1364-6613. https://www.cambridge.org/core/journals/behavioral-and-brain-sciences/article/abs/on-the-proper-treatment-of-connectionism/4B8871A82A932DB96D183AAC9C0CF037(2023) (Mar. 1988).

163. Lake, B. M., Ullman, T. D., Tenenbaum, J. B. & Gershman, S. J. Building machines that learn and think like people. Behavioral and Brain Sciences 40 (2017).

164. Kipf, T., Fetaya, E., Wang, K.-C., Welling, M. & Zemel, R. Neural Relational Inference for Interacting Systems en. in Proceedings of the 35th International Conference on Machine Learning ISSN: 2640-3498 (PMLR, July 2018), 2688–2697. https://proceedings.mlr.press/v80/kipf18a.html (2022).

165. Hill, F., Santoro, A., Barrett, D., Morcos, A. & Lillicrap, T. Learning to Make Analogies by Contrasting Abstract Relational Structure in International Conference on Learning Representations (2019).

166. Saxe, A., Sodhani, S. & Lewallen, S. J. The Neural Race Reduction: Dynamics of Abstraction in Gated Networks en. in Proceedings of the 39th International Conference on Machine Learning (PMLR, June 2022), 19287–19309. https://proceedings.mlr.press/v162/saxe22a.html (2023).

167. Craik, K. J. W. The Nature of Explanation (1943).

168. Frankland, S. M. & Greene, J. D. Concepts and Compositionality: In Search of the Brain’s Language of Thought. Annual Review of Psychology 71, 273–303. 10.1146/annurev-psych-122216-011829(2023) (2020).

169. Johnson, J. et al. CLEVR: A Diagnostic Dataset for Compositional Language and Elementary Visual Reasoning in (2017), 2901–2910. https://openaccess.thecvf.com/content_cvpr_2017/html/Johnson_CLEVR_A_Diagnostic_CVPR_2017_paper.html(2023).

170. Ito, T. et al. Compositional generalization through abstract representations in human and artificial neural networks arXiv:2209.07431 [q-bio]. Sept. 2022. http://arxiv.org/abs/2209.07431 (2023).

171. Lake, B. M. & Baroni, M. Human-like systematic generalization through a meta-learning neural network. en. Nature 623. Number: 7985 Publisher: Nature Publishing Group, 115–121. issn: 1364-6613. https://www.nature.com/articles/s41586-023-06668-3(2023) (Nov. 2023).

172. Lepori, M., Serre, T. & Pavlick, E. Break it down: Evidence for structural compositionality in neural networks. Advances in Neural Information Processing Systems 36 (2024).

173. Fukushima, K. Visual Feature Extraction by a Multilayered Network of Analog Threshold Elements. IEEE Transactions on Systems Science and Cybernetics 5, 322–333. issn: 2168-2887 (Oct. 1969).

174. Maas, A. L., Hannun, A. Y., Ng, A. Y., et al. Rectifier nonlinearities improve neural network acoustic models in Proc. icml 30. Issue: 1 (Atlanta, Georgia, USA, 2013), 3.

175. Tsuchida, R., Roosta-Khorasani, F. & Gallagher, M. Invariance of Weight Distributions in Rectified MLPs arXiv:1711.09090 [cs, stat]. May 2018. http://arxiv.org/abs/1711.09090 (2022).

176. Tsuchida, R., Roosta, F. & Gallagher, M. Richer priors for infinitely wide multi-layer perceptrons arXiv:1911.12927 [cs, stat]. Nov. 2019. http://arxiv.org/abs/1911.12927 (2022).

177. Han, I. et al. Fast Neural Kernel Embeddings for General Activations arXiv:2209.04121 [cs, stat]. Sept. 2022. http://arxiv.org/abs/2209.04121 (2022).

178. Schöolkopf, B. The Kernel Trick for Distances in Advances in Neural Information Processing Systems 13 (MIT Press, 2000). https://proceedings.neurips.cc/paper/2000/hash/4e87337f366f72daa424dae11df0538c-Abstract.html(2024).

179. Hu, G. Y. & O’Connell, R. F. Analytical inversion of symmetric tridiagonal matrices. Journal of Physics A: Mathematical and General 29, 1511. 10.1088/0305-4470/29/7/020 (Apr. 1996).

180. He, K., Zhang, X., Ren, S. & Sun, J. Delving Deep into Rectifiers: Surpassing Human-Level Performance on ImageNet Classification in (2015), 1026–1034. https://openaccess.thecvf.com/content_iccv_2015/html/He_Delving_Deep_into_ICCV_2015_paper.html(2023).

181. R Core Team. R: A Language and Environment for Statistical Computing https://www.R-project.org/ (R Foundation for Statistical Computing, Vienna, Austria, 2022).

182. Wickham, H. ggplot2: Elegant Graphics for Data Analysis isbn: 978-3-319-24277-4. https://ggplot2.tidyverse.org(Springer-Verlag New York, 2016).

183. Pedersen, T. L. patchwork: The Composer of Plots https://CRAN.R-project.org/package=patchwork(2022).

184. Paszke, A. et al. in Advances in Neural Information Processing Systems 32 8024–8035 (Curran Associates, Inc., 2019). http://papers.neurips.cc/paper/9015-pytorch-an-imperative-style-high-performance-deep-learning-library.pdf.

185. Knuth, D. E. Johann Faulhaber and sums of powers. en. Mathematics of Computation 61, 277–294. issn: 1364-6613. https://www.ams.org/mcom/1993-61-203/S0025-5718-1993-1197512-7/(2024) (1993).

186. Böottcher, A. & Grudsky, S. M. Spectral properties of banded Toeplitz matrices (SIAM, 2005).

187. Gill, J. The use of the sequence Fn(z)=fnf1(z) in computing fixed points of continued fractions, products, and series. en. Applied Numerical Mathematics 8, 469–476. issn: 1364-6613. https://www.sciencedirect.com/science/article/pii/016892749190109D(2023) (Dec. 1991).

